# A combined RNA-Seq and comparative genomics approach identifies 1,085 candidate structured RNAs expressed in human microbiomes

**DOI:** 10.1101/2020.03.31.018887

**Authors:** Brayon J. Fremin, Ami S. Bhatt

**Affiliations:** Department of Genetics, Stanford University, CA, 94305, USA; Department of Medicine (Hematology), Stanford University, CA, 94305, USA

## Abstract

Structured RNAs play varied bioregulatory roles within microbes. To date, hundreds of candidate structured RNAs have been predicted using informatic approaches by searching for motif structures in genomic sequence data. However, only a subset of these candidate structured RNAs, those from culturable, well-studied microbes, have been shown to be transcribed. As the human microbiome contains thousands of species and strains of microbes, we sought to apply both informatic and experimental approaches to these organisms to identify novel transcribed structured RNAs. We combine an experimental approach, RNA-Seq, with an informatic approach, comparative genomics across the human microbiome project, to discover 1,085 candidate, conserved structured RNAs that are actively transcribed in human fecal microbiomes. These predictions include novel tracrRNAs that associate with *Cas9* and RNA structures encoded in overlapping regions of the genome that are in opposing orientations. In summary, this combined experimental and computational approach enables the discovery of thousands of novel candidate structured RNAs.

## Introduction

Microbial structured RNAs are involved in biological processes ranging from cis-antisense regulation to targeted regulation of diverse sets of genes ^1^. Given their exciting demonstrated roles in bioregulation, to date, there has been enthusiasm for developing methods to more comprehensively identify candidate structured RNAs. Previously, comparative genomics has been applied for this purpose to high effect, revealing hundreds of novel candidate structured RNA motifs ^2^. In general, these comparative genomic approaches involve extensive clustering of subsequences of intergenic regions from many organisms followed by structural motif prediction ^3,4^. In the last decade, hundreds of structured RNAs in bacteria have been predicted using comparative genomics ^2,5^. The small subset of these RNAs that have been validated and carefully characterized have displayed an exciting array of activities, like self-cleaving ribozymes ^6,7^.

While powerful, a drawback of this comparative genomics approach is that in any given experiment, only a select set of intergenic regions are considered; this is done to accommodate computational limitations. For example, comparative genomics approaches often subset intergenic regions to bias towards known classes of structured RNAs, like ribozymes or specific types of riboswitches, which have a tendency to be found in specific genomic contexts ^2^. As a consequence of computational limitations and subsetting of intergenic regions, much of the intergenic search space in microbes remains unexplored and many new structured RNAs likely remain to be discovered.

Another caveat of a comparative genomics strategy is that it does not directly show that a given motif is a structured RNA as opposed to, for example, a single-stranded DNA motif ^2^. Demonstrating that the motif is transcribed makes a stronger case that the motif is a structured RNA. However, validating hundreds of structured RNAs identified in tens to hundreds of different organisms at an RNA-level is technically challenging using traditional culturing approaches of each organism. Additionally, most existing RNA-Seq datasets in bacteria bias against small structured RNAs as small transcripts are typically removed in the process of size selection in standard RNA-Seq protocols ^8^. It then follows that RNA-Seq protocols that do not size select may be useful both to predict and demonstrate transcription of new structured RNAs in bacteria ^8^.

To overcome these limitations, here, we carried out a combined approach leveraging both RNA-Seq without fragment size selection on fecal samples and comparative genomics. Our finalized list of 1,085 novel, candidate structured RNAs are all found in transcribed, intergenic regions. Additionally, they are conserved across the Human Microbiome Project phase 2 (HMP2) ^9^ with strong evidence of computationally predicted secondary structure. We use this approach to identify novel tracrRNAs and structured RNAs that occupy the same genomic region in opposing orientations. Overall, we show that this combined approach of RNA-Seq and comparative genomics is a powerful way to discover transcribed human host-associated structured RNAs from microbes in fecal samples.

## Results

### RNA-Seq along with comparative genomics enables targeted, high-throughput predictions of structured RNAs

To identify novel structured RNAs, we created the following workflow (Figure 1). In this study, we analyzed metagenomics and RNA-seq without fragment size selection on four taxonomically diverse fecal samples from four different human subjects (A – healthy adult, B – patient with hematological disorder undergoing treatment, C – patient with cancer undergoing treatment, D – patient with Alzheimer’s disease). These are the same samples used in a previous study ^10^. First, metagenomic DNA sequencing data from four human stool samples was obtained; these data had previously been subjected to computational assembly and the resultant contigs (longer contiguous DNA sequences that are “in silico” assembled from smaller DNA fragments) were collected. RNA-Seq without fragment size selection was performed on these same samples and the resultant reads were aligned to the *de novo* assemblies. The intergenic regions in these *de novo* assemblies that had RNA-Seq signal were searched against HMP2 ^9^ metagenomes. For each intergenic region that was found to be transcribed, we searched for conserved regions across HMP2 ^9^ to test if we could make strong comparative genomics predictions of RNA structure. Finally, we structurally aligned these regions using CMfinder (version 0.4.1) ^2,11^ and retained sequences with high scoring structures (see Methods, Figure 1). Motifs predicted independently multiple times were removed. We further filtered these putative structured RNAs to exclude any regions that had significant BLASTx ^12^ hits to the nr database or significant potential to be coding determined via RNAcode ^13^. We reassessed if these specific structures were expressed (using a threshold of Reads Per Kilobase Million (RPKM) > 20). To propose only novel candidate structured RNAs, we next determined which of our predictions were not found in Rfam 14.1, a database of known structured RNAs ^14^. This enabled highly stringent predictions of novel structured RNAs (Figure 1).

**Figure 1:**
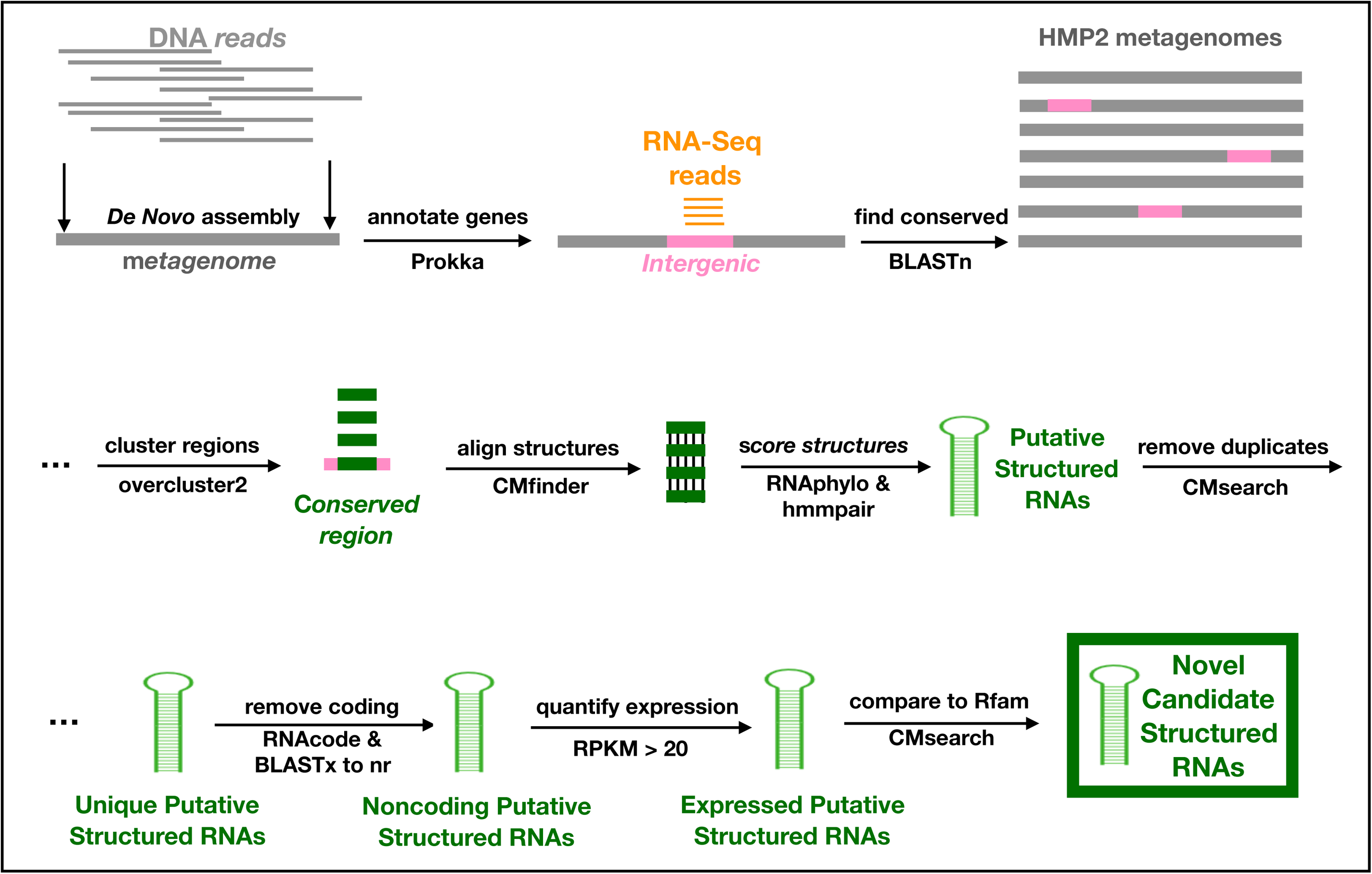
Candidate structured RNA prediction workflow. First, genes in *de novo* assembled metagenomes are annotated with Prokka ^17^. Non-coding genomic regions are next scanned for RNA-Seq reads that align to these intergenic regions. Homologs of these transcribed intergenic regions are identified by homology searching in the HMP2 ^9^ metagenome dataset via BLASTn ^12^. Conserved regions, defined as sequences for which there are homologs in the HMP2 dataset, are clustered. This near-final list of putative structured RNAs was filtered to exclude duplicates, coding regions, and those already found in Rfam 14.1^2,14^.

Taking this approach, we found that of the 157,995 intergenic regions in our four fecal samples tested, 9,261 regions contained RNA-Seq signal (Figure 2A). Using BLASTn ^12^ to search for these regions across HMP2 ^9^ and overcluster2 to identify conserved regions within these intergenic regions (see Methods), we identified 13,377 conserved regions among these 9,261 intergenic regions. From these conserved regions, we identified 2,023 putative structured RNAs. We include a stringent filter against these putative structured RNAs being coding by requiring no significant hits to the nr database via BLASTx ^12^ and no significant p-values from RNAcode ^13^, which narrowed our list of putative structured RNAs to 1,862. To avoid proposing the same structure multiple times, we filtered this list to 1,525 unique putative structured RNAs by ensuring the same structures do not match the same regions using CMsearch ^15^. To ensure that the predicted structure motifs were transcribed, we calculated RPKM values specifically for the structure motifs (instead of the entire intergenic region), and found that 1,213 of these structures contained strong RNA-Seq signal (RPKM > 20).

**Figure 2:**
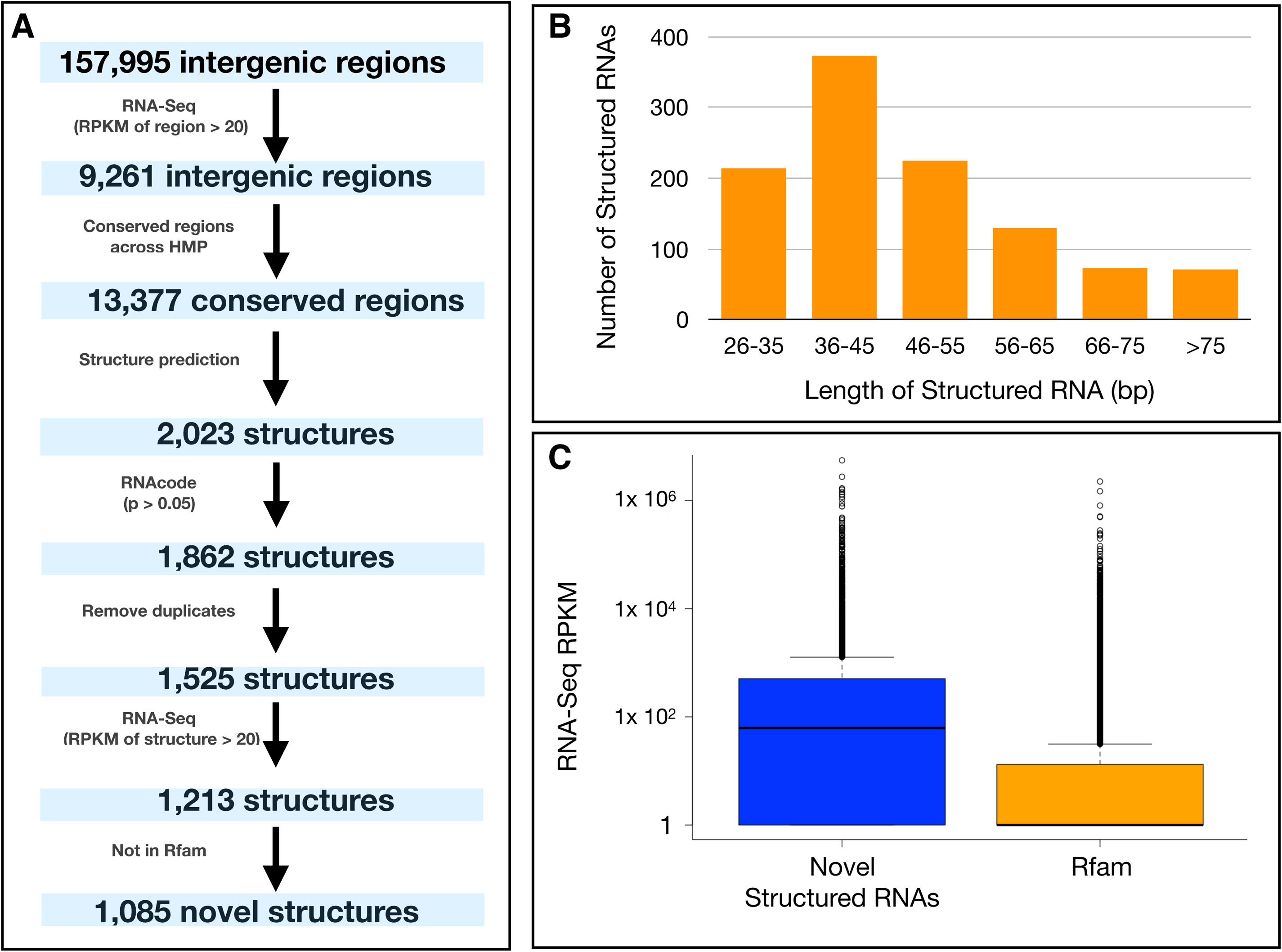
Assessment of novel, candidate structured RNAs. (A) Quantitative flowchart of our combined approach in which we start with 157,995 intergenic regions in our four fecal samples and identify a final list of 1,085 novel, candidate structured RNAs. (B) Length distribution of the 1,085 candidate structured RNAs. (C) Boxplot of RNA-Seq RPKM values for the novel 1,085 candidate structured RNAs compared to known structured RNAs that currently exist in Rfam 14.1^2,14^.

Of these 1,213 predicted and transcribed structured RNAs, 128 were previously known structured RNAs present in Rfam 14.1 ^14^ (CMsearch ^15^ e-value < 1 x 10 ^5^). Included in these known structured RNAs, for example, were tRNA and rRNA predictions that were overlooked by Aragorn ^16^ and barnnap ^17^, respectively, but were present in Rfam 14.1 ^14^ (Table S1). Eight of these were proposed by recent literature ^2^ (Table S1). Our ability to rediscover these 128 previously known structured RNAs with our approach lended support to the overall workflow; the remaining 1,085 predicted structured RNAs discovered by this approach were deemed to be novel. We characterize these novel candidate structured RNAs by their taxonomy, genomic neighborhood, genomic position relative to other structured RNAs, and expression levels in fecal microbiomes (Figure 2A, Table S2). All candidate structured RNAs we predict are given the following naming system: SampleX_Y, where X is the sample we initially predicted the structure in (A,B,C or D) and Y is a unique number given to each prediction. These novel candidate structured RNAs range from 29 to 134 base pairs in length (Figure 2B, Table S2) and are expressed in the fecal microbiome (Figure 2C, Table S3). SampleBMeta_586 was the most highly expressed structured RNA across the samples at 887,778 RPKM (Table S3). The second most highly expressed was a previously identified structured RNA, Bacteroidales-1 ^5^ (Table S3).

### RNA-Seq evidence of structured RNAs can be made simultaneously across diverse microbes

Next, we tested if our RNA-Seq based approach captured structured RNAs across the different taxa present in host-associated microbiomes. Within each of the four diverse fecal samples with RNA-Seq data, we taxonomically display all the novel 1,085 structured RNAs that are expressed (RPKM > 20) in the fecal microbiome (Figure 2A-D). Each structured RNA can occur multiple times in a sample or in multiple samples. We found these 1,085 structured RNAs 4,351 times in the Samples A-D in total. Of these 4,351 predictions, 2,582 were expressed at a level of RPKM > 20. We detect 694, 722, 684, and 482 expressed, novel structured RNAs in samples A, B, C, and D, respectively (Figure 2A-D). For each of these novel structured RNA predictions, its taxonomy across HMP2 ^9^ can be more extensive than the taxa shown here (Figure 2, Table S2). While 862 of the candidate structured RNAs are exclusive to one phyla, the remaining 223 can be found in two or more different phyla (Table S2).

Of note, 120 of these 1,085 novel structured RNAs share structural similarity with known structured RNAs (CMsearch ^15^ E-value > 1 x 10 ^5^). These structured RNAs are found in different taxa and genomic contexts than the known structured RNAs they are similar to; however, they may still perform similar functions in these organisms. For example, there are strong structural similarities between *DsrA* and our prediction, SampleCMeta_143 (Table S1). *DsrA* is a trans-acting structured RNA that regulates transcription and translation of distant genes, including, for example, those involved in stress response ^18^. SampleCMeta_143 is found in Clostridia, whereas *DsrA* is more specific to Proteobacteria (Table S1). Additionally, SampleAMeta_243 and SampleBMeta_667 are structurally similar to *Spot_42* (Table S1). *Spot_42* regulates carbohydrate metabolism and uptake ^19^ and is quite specific to Proteobacteria ^14^, while SampleAMeta_243 and SampleCMeta_667 are found in Bacteroidetes and Firmicutes, respectively (Table S2).

### Novel tracrRNAs occur directly upstream of *Cas9*

In our genomic neighborhood analysis, one interesting pattern we observed was the prediction of structured RNAs directly upstream of *Cas9* in a diverse set of microbes. Both the specific secondary structure and orientation of the structured RNA prediction relative to *Cas9* varied across taxa (Figure 3). We observed stranded RNA-Seq signal for these RNA structures even in cases where the structured RNA was antisense to the predicted mRNA transcript of *Cas9*. We find these four distinct structures directly adjacent to *Cas9* in *Bifidobacterium* (Figure 4A), *Bacteroides* (Figure 3B), *Subdoligranulum* (Figure 4C), and *Lachnospiraceae* (Figure 4D). TracrRNAs are commonly encoded directly upstream of *Cas9*, suggesting these are novel tracrRNAs being expressed in the fecal microbiome. These are four additional structures of tracrRNAs that can be added to the two structures that currently exist in Rfam 14.1 ^14^. Previous work, which performed small RNA-Seq on isolates, suggests that tracrRNAs often have no conservation of structure across diverse taxa and can have different sequences, localization, and even orientation within type II CRISPR-Cas systems ^20^. RNA-Seq without size selection in the microbiome was critical to identifying the transcription and strand orientation of these new tracrRNAs in uncultured microbes.

**Figure 3:**
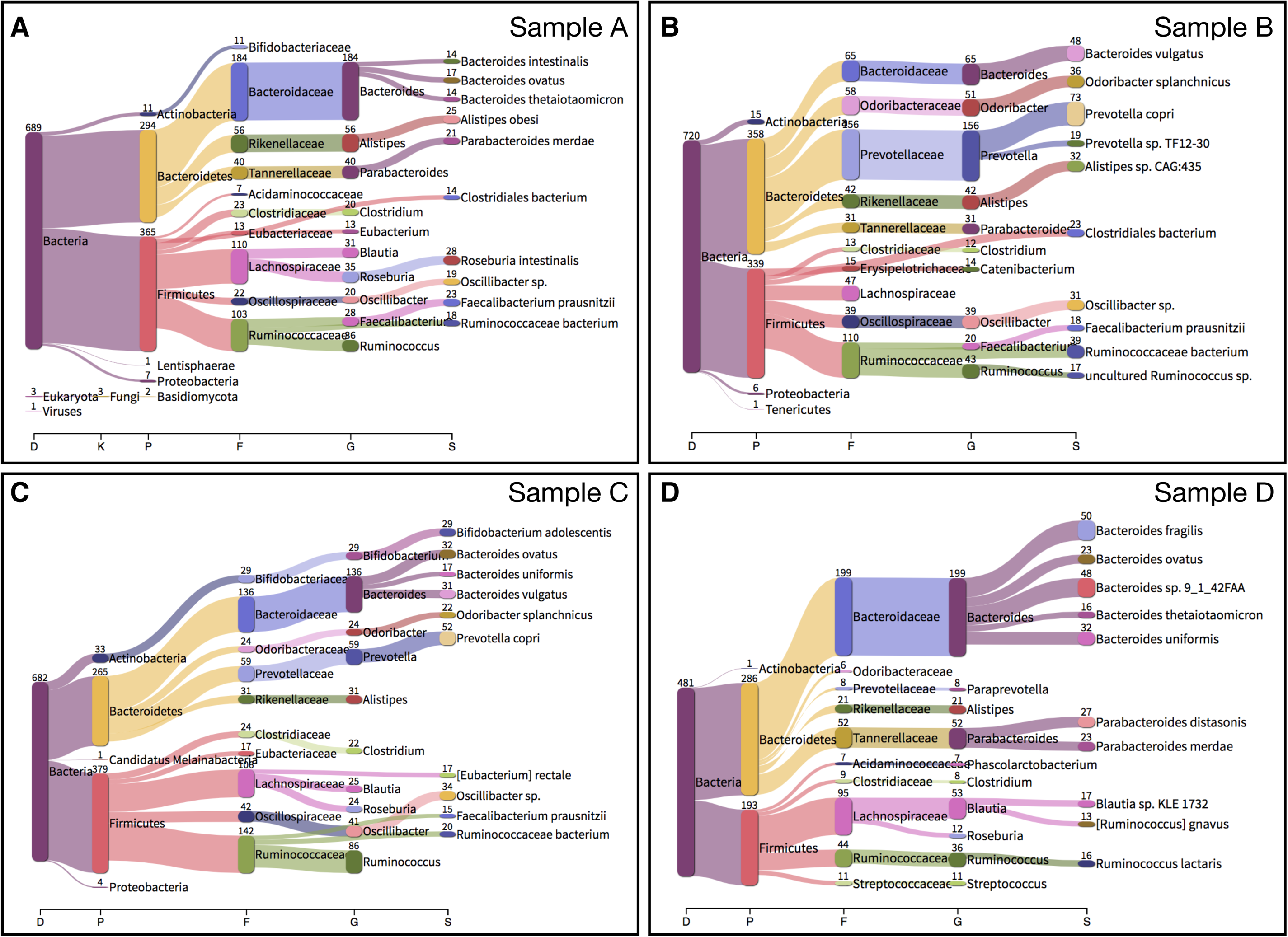
Structured RNA predictions can be made across diverse taxa simultaneously. (A-D) Taxonomic classifications of the contigs on which all of the predicted structured RNAs exist in Samples A, B, C, and D. This is shown at multiple taxonomic levels. D = Domain, K = Kingdom, P = Phylum, F = Family, G = Genus, S = Species.

**Figure 4:**
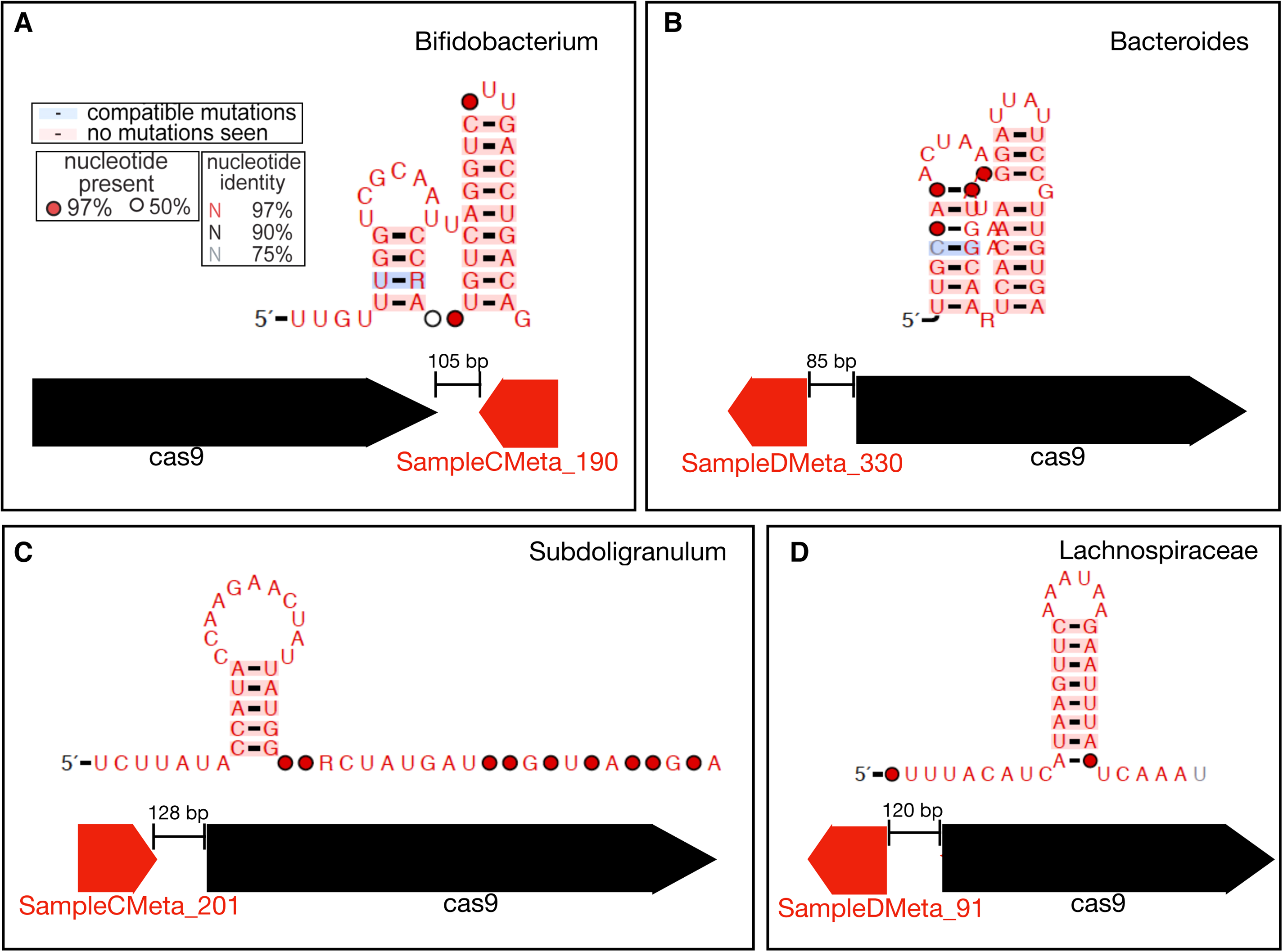
Novel tracrRNAs have different conserved stem-loop structures. (A-D) Structure diagrams, showing the consensus features, of novel tracrRNAs associated with *Cas9*. Below the diagrams, we show the genomic localization and RNA-Seq signal distribution across the tracrRNA. In panels A, B, C, and D, we show novel tracrRNAs predicted in *Bifidobacterium, Bacteroides, Subdoligranulum*, and *Lachnospiraceae*, respectively.

**Figure 5:**
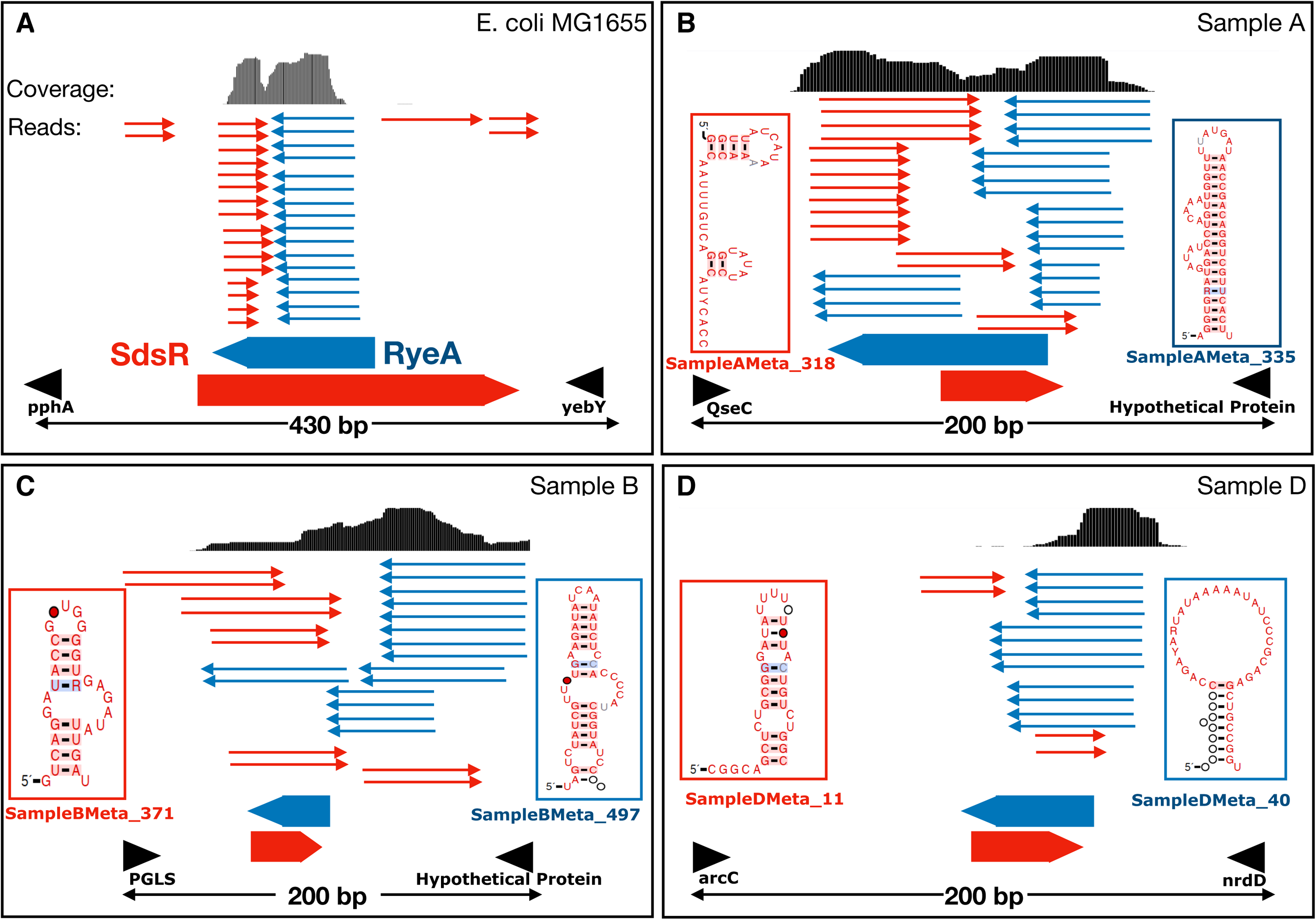
Structured RNAs can be encoded in an overlapping manner in opposing strand orientations. (A) IGV coverage plot of the known toxin-antitoxin system *SdsR* and *RyeA* in *E. coli* MG1655. To our knowledge, this is the only previously characterized instance of overlapping structured RNAs in opposing orientations. We observe stranded RNA-Seq signal on both strands, which we indicate using reads representing the directionality of the coverage. (B-D) IGV coverage plots of additional overlapping structured RNAs in the fecal microbiome. Like *SdsR* and *RyeA*, we observe stranded RNA-Seq signal on both strands in these examples. Consensus diagrams for these structures are also provided.

### RNA-Seq identifies novel structured RNAs that overlap in opposing strand orientations

Unexpectedly, we found many instances of overlapping structured RNA predictions on opposite strands. We found that 80 of our predictions, or 40 pairs of structured RNAs, overlapped other predictions in the opposing strand orientation (Figure S2, Table S2). Cis-antisense RNAs are common in bacteria; however, they are often found as a structured RNA overlapping a coding transcript ^2,21,22^. Structured RNAs that overlap one another are not unprecedented ^23–25^, but have not been extensively reported in bacteria. To our knowledge, the only such instance of this occurs in *E. coli* and *Salmonella enterica* serovar Typhimurium - structured RNAs *ryeA* and *sdsR*^*26,27*^ (Figure 4A). In previous literature, there are cases in which stable RNA secondary structures can be computationally predicted in both orientations, especially for miRNAs, and only the “most stable” is reported as the correct orientation ^28^. Here, we present many cases of overlapping secondary structures across both strands with RNA-Seq signal supporting both structures. For example, SampleAMeta_318 and SampleAMeta_335 (Figure 4B), SampleBMeta_371 and SampleBMeta_497 (Figure 4C), and SampleDMeta_11 and SampleDMeta_40 (Figure 4D) are distinct structured RNA pairs that overlap each other in opposing orientations. We show the coverage across the intergenic region each of these are found in, as well as the direction of reads, demonstrating transcription of both predicted structured RNAs. This phenomenon of overlapping structured RNAs was previously thought of as uncommon in bacteria.

## Discussion

Even though structured RNAs in microbes play key roles in bioregulation, many structured RNAs remain undiscovered. Applying a method like a modified RNA-Seq approach without fragment size selection to human fecal samples enables the quantification of many host-associated, predicted structured RNAs; furthermore, this can be done in a culture-free manner and can be performed across multiple taxa simultaneously. Thus, this approach greatlyc expands our ability to predict novel structured RNAs. We use RNA-Seq to reduce the intergenic search space of structured RNA predictions to relevant, transcribed regions (5.9% of intergenic regions in our samples) and also to support that structured motifs predicted via comparative genomics are likely transcribed RNA motifs.

Here, we build a resource of 1,085 novel host-associated structured RNAs founded both on comparative genomics and evidence of transcription in the fecal microbiome. Among these predictions, we identify distinct structured RNAs found directly upstream of *Cas9* across different taxa. The structure and genomic neighborhood suggests that these are novel tracrRNAs expressed in human fecal microbiomes. Moreover, we identify 80 structured RNAs that also contain structured RNAs within the same region on the opposite strand. This suggests that such phenomenon of overlapping structured RNAs may not be as rare as it once seemed in bacteria. In the one known case in bacteria, some evidence suggests that *SdsR* and *RyeA* act as a novel toxin-antitoxin system ^29,30^. Given this finding, some of these 40 structured RNA pairs may be additional members of this novel type of toxin-antitoxin system. However, they may regulate each other as part of a different type of system. Future experiments will be needed to elucidate the functions and mechanisms of these structured RNAs.

While this approach is high in throughput and generates many intriguing predictions, there are several limitations to this approach. First, this method relies heavily on transcription levels of the structured RNAs. Certain RNAs may serve regulatory functions and their transcription may be tightly regulated. Here, we cannot predict structured RNAs that are found within rare taxa of the gut microbiome, nor do we detect those that are lowly expressed in this niche. Of course, structured RNAs in bacteria that do not inhabit the gut are entirely overlooked by this approach. This limitation can be overcome by applying the entire approach we describe in samples from other microbiomes of interest (such as ocean sediment, soil, etc.). Additionally, some structured RNAs within the fecal microbiome may not be expressed in the four fecal samples we performed RNA-Seq on; however, they may be expressed in the microbiomes of other individuals or under different conditions. Moreover, we are predicting structured RNAs in fragmented metagenomic assemblies as opposed to complete, isolated genomes; this means that the taxonomic classification of these structured RNAs may not be highly resolved (e.g., sometimes we can only trace a predicted RNA down to the family or order level). This is in part due to limitations in the reference databases we use for taxonomic classification, and in part due to the limited sensitivity and occasional misclassification that happens when classifying contigs as opposed to entire genomes. Finally, unlike previous structured RNA discovery approaches, which use comparative genomics along with genomic neighborhood information to select for specific predicted functions such as riboswitches, the structured RNAs we predict in this work can be less straightforward to functionally characterize. Because we are not subsetting intergenic regions to enrich for specific types of structured RNAs, it becomes more difficult to make informed inferences on the functions of our predictions. For example, these structured RNAs could be trans-regulatory and regulate genes in the organism found on different contigs in the sample. Further investigation is needed to determine functions.

While we are limited in the number of existing bacterial small transcriptomics datasets, which tend to be few and limited to model organisms ^31^, we have a plethora of assembled bacterial genomes both from isolates ^32^ and metagenomic samples ^9^. This is why comparative genomics approaches, despite their computational and experimental shortcomings have been among the most powerful means to predict hundreds of new candidate structured RNAs in bacteria ^2,5^. By applying RNA-Seq to the fecal microbiome, we can simultaneously obtain transcriptomic information across many taxa without culturing restrictions. Using RNA-Seq as a means to subset, we can predict with greater confidence that the motifs we find are indeed structured RNAs. This also allows us to predict rarer and more diverse structured RNAs that will be overlooked by common subsetting tactics in comparative genomics approaches alone. Thus, the combined strengths of large-scale RNA-Seq and comparative genomics have yielded a powerful approach to rigorously predict over a thousand novel host-associated structured RNAs.

## Supporting information

RNA Structures

Alignment Files

RPKM values

RNA Structure Information

RNA Structure Overlaps

## Acknowledgements

The authors would like to thank Tessa M. Andermann, Ekaterina Tkachenko, and Joyce B. Kang for major contributions to the fecal biobank of blood and marrow transplantation patients at Stanford hospital, especially Dr. David Miklos who oversees the biobanking protocol and Dr. Andrew Rezvani. We would like to thank the physicians, patients, and nurses involved in collection. We thank Christina Wyss-Coray for collecting samples from patients with Alzheimer’s disease, some of which were used in this study. We appreciate sample collection feedback from Victor Henderson and Tony Wyss-Coray. Sequencing costs were supported via NIH S10 Shared Instrumentation Grant (1S10OD02014101) and Damon Runyon Clinical Investigator Award to ASB, Stanford ADRC grant # P50AG047366. B.J.F is supported by National Science Foundation Graduate Research Fellowship DGE-114747.

## Methods

### Subject Recruitment

RNA-Seq was performed on stool samples from individuals in a variety of health states. Informed consent was obtained for all participants. All subjects were recruited at Stanford University as a part of one of three IRB-approved protocols for tissue biobanking and clinical metadata collection (PIs: Dr. Ami Bhatt, Dr. Victor Henderson, Dr. David Miklos).

### Fecal Sample Storage

Stool was immediately stored in 2 mL cryovials and frozen at −80 °C, and the samples were not thawed until lysis. For RNA extraction applications, 1.3 grams of fecal samples were preserved in 700 mL of RNALater at −80 °C.

### Cell Lysis for Transcriptomics in Fecal Samples

Stool (150 mg) was suspended in 600 μL of Qiagen RLT lysis buffer supplemented with one percent beta-mercaptoethanol and 0.3 U/μL Superase-In (Invitrogen). The suspension was subjected to bead beating for 3 minutes using 1.0 mm Zirconia/Silica beads. This was performed with a MiniBeadBeater-16, Model 607 (Biospec). The lysed solution was centrifuged at room temperature for 3 minutes at 21,000 x g to pellet cellular debris and the supernatant was decanted into 2 mL tubes.

### Cell Lysis for Transcriptomics in E. coli

NR-2653 *Escherichia coli* K-12 MG1655 was obtained from BEI Resources. *E. coli* was grown in LB broth to an OD600 of 0.4 at 37 °C. Aliquots (10 mL) of the community were centrifuged in 15 mL tubes at 4000 x g for 30 minutes at room temperature. Bacterial pellets were suspended in 600 μL Qiagen RLT lysis buffer supplemented with one percent beta-mercaptoethanol and 0.3 U/μL Superase-In (Invitrogen). The suspension was subjected to bead beating for 3 minutes using 1.0 mm Zirconia/Silica beads. This was performed with a MiniBeadBeater-16, Model 607 (Biospec). The lysed solution was centrifuged at room temperature for 3 minutes at 21,000 x gto pellet cellular debris and the supernatant was decanted into 2 mL tubes.

### Metagenomics

DNA was extracted from each fecal sample with the DNA Stool Mini Kit (Qiagen) according to manufacturer’s protocols. Samples were then exposed to bead beating for 3 minutes. 1 ng of DNA was used to create Nextera XT libraries according to manufacturer’s instructions (Illumina).

### Transcriptomics Without Fragment Size Selection

We performed metatranscriptomics as follows: 15 uL of proteinase K (Ambion, 20 mg/mL) was added to 600 μL of lysate. After incubation for 10 minutes at room temperature, samples were centrifuged at 21,000 x g for 3 minutes and the supernatant was collected. An equal volume of Phenol/Chloroform/Isoamyl Alcohol 25:24:1 (pH. 5.2) was applied and was vortexed for three minutes. The mixture was centrifuged at 21,000 x g for three minutes. The aqueous phase was extracted. This phenol chloroform step was repeated once more and the aqueous phase was extracted. This final aqueous phase was ethanol precipitated with 0.1 percent volume 3M sodium acetate and 2.5 volumes of 100 percent ethanol. The RNA was further purified using the RNAeasy Mini plus Kit (Qiagen) using manufacturer’s protocols. Any remaining DNA was degraded via Baseline-ZERO-DNase (Epicentre) using manufacturer’s protocols. RNA was fragmented for 15 minutes at 70 °C using RNA Fragmentation Reagent (Ambion) using manufacture protocols. The fragmented RNA was purified with miRNAeasy Mini Kit (Qiagen) using manufacturer’s protocols. Elution was performed in 15 mL of water. rRNA was depleted using RiboZero-rRNA Removal Kit for Bacteria (Illumina) using half reactions of manufacture protocol. This was purified with RNAeasy MinElute Cleanup Kit (Qiagen), eluting in 20 uL of water. The fragments, in 18 mL volume, were subjected to T4 PNK Reaction (NEB M0201S) with addition of 1μL Superase-In (Invitrogen), 2.2 mL 10X T4 PNK Buffer, and 1 mL T4 PNK (10U/mL). This reaction was purified again with RNAeasy MinElute Cleanup (Qiagen) using manufacturer’s protocols. The concentration was determined with Qubit RNA HS Assay Kit (Invitrogen). With 100 ng as input, libraries were prepared using NEBNext Small RNA Library Prep Set for Illumina (NEB, E7330), using manufacturer’s protocols. The resulting DNA libraries were purified using MinElute PCR Purification Kit (Qiagen) using manufacturer’s protocols. Libraries were sequenced with 1×75 bp reads on a NextSeq 500.

### De Novo Assembly

Quality trimmed metagenomic reads were assembled using metaSPAdes 3.7.0 ^33^ as previously performed ^10^. For all samples, a maximum of 60 million metagenomic reads were used to generate assemblies. Samples sequenced to higher depth were randomly subsetted to 60 million reads for assembly purposes to both ensure relatively similar numbers of gene predictions and limit computational requirements in assembly and downstream predictions.

### Read Mapping

RNA-seq reads were trimmed with trim galore version 0.4.0 using cutadapt 1.8.1 ^34^ with flags –q 30 and –illumina. RNA-seq reads were mapped to the annotated metagenomic assembly using bowtie version 1.1.1 ^35^. IGV ^36^ was used to visualize coverage.

### Known Structured RNA Predictions

Prokka v1.12 ^17^ was used to call open reading frames from the metagenomics assemblies using the –meta option. Annotations were facilitated by many dependencies ^12,16,37,38^. Transfer RNAs were predicted by Aragorn ^16^. Ribosomal RNAs were predicted with barnnap ^17^. Structured RNAs were predicted by comparing to a database of known structured RNAs called Rfam 14.1 ^14^ with the following restrictions to include: CRISPR, antisense, antitoxin, miRNA, ribozyme, RNA, snRNA, Intron, Cis-reg, IRES, frameshift_element, leader, riboswitch, thermoregulator, and bacteria. Certain predictions, such as tRNAs and rRNAs not detected by Aragorn ^16^ and Barnnap ^17^, respectively, or structured RNAs not previously identified in bacteria, were not included in this search. Intergenic regions were determined based on these predictions from Prokka ^17^.

### RNA-Seq and Comparative Genomics Analysis Workflow

To identify structured RNAs, we employed a similar comparative genomics approach to established protocols ^2,5^; however, we also incorporated information gained from RNA-Seq into this pipeline. For Samples A, B, C, and D, we identified all intergenic regions with an RNA-Seq RPKM greater than 20 RPKM. We performed nucleotide BLASTn ^12^ on these intergenic regions to compare these sequences with those from 1,773 human metagenomic assemblies within the HMP2 dataset ^9^. The HMP2 dataset consists of metagenomic sequencing from four different body sites in hundreds of healthy human subjects. We clustered similar subsequences based on BLASTn scores ^2,5^ using a single-linkage clustering algorithm called overcluster2 (Weinberg,Z., unpublished open-source software, available at http://weinberg-overcluster2.sourceforge.io). RNA structural alignments were inferred from these clusters using CMfinder version 0.4.1 ^2,11^. RNA structures were drawn using R2R ^39^. Scores of these structures were calculated using updated RNAphylo and hmmpair, as performed previously ^2^. The best scoring structure for each intergenic region was drawn and the score for this structure was recorded. Duplicate structures that were independently predicted were identified by searching across the same regions (CMsearch ^15^ e value < 1 × 10 ^5^) and were discarded. We removed any regions with significant BLASTx ^12^ evalues (< 0.000001) to nr. Additionally, these clusters were aligned with ClustalW ^33,40^ using default parameters, and tested for coding potential using RNAcode ^13^ version 3 ^13^. Sequences with significant RNACode p values (< 0.05), which suggests that they are more likely to be coding regions of the genome, were discarded. We discarded any structured RNA that did not contain RNA-Seq signal above 20 RPKM, considering these narrowed boundaries and strand information. Finally, we discarded any prediction homologous to known RNA structures by removing those sequences that had a significant hit to Rfam 14.1 ^14^ (CMsearch ^15^ e value < 1 × 10 ^5^).

### Motif Searches

In separate searches after our Prokka ^17^ predictions were generated, we then searched for the Rfam ^14^ database against these predictions to determine overlap. CMsearch version 1.1.2 ^15,41^ was used for all searches with an e-value cutoff of 1 × 10 ^5^ considered as a significant match.

### Taxonomic Classification of Technologies

The contigs in which predictions were identified were classified using Kraken ^42^ against a custom database. This database included all bacteria, viral, and fungal genomes in NCBI GenBank of at least scaffold quality as of February 2019. In the case of initial predictions, the entire contigs that regions were predicted within were classified. In the case of HMP2 ^9^ predictions, contigs in which clustered blast hits were identified were classified. These classifications were visualized with Pavian ^43^.

### Genomic Neighborhood Analysis of Small Protein Families

To determine the genes that are present within the vicinity of candidate structured RNAs, amino acid sequences of genes that existed within 5 kb of each predicted structured RNA was identified. We then compared these sequences against the Conserved Domain Database (CDD) ^44^, using RPS-blast ^12^. A hit was considered significant if: e-value ≤ 0.05 and the protein aligns to at least 80% of the PSSM’s length.

## Figure Legends

File S1: Visualization of best scoring secondary structures.

All best scoring structures from structural alignments of all structured RNA predictions are provided.

File S2: Alignment files of the novel candidate structured RNAs.

We provide alignment files for these predicted structured RNAs.

Table S1: Overlap of structured RNA motifs.

We display all overlaps among structured RNAs predicted. We also provide information on the structured RNAs that are significant hits to structures in Rfam 14.1 ^2,14^ database (CMsearch ^15^ e value < 1 × 10 ^5^), and are thus not part of the novel 1,085 predictions. Finally, we show those novel predictions that are similar structures to those found in Rfam 14.1 ^2,14^ database (CMsearch ^15^ e value < 1 × 10 ^5^).

Table S2: Conserved region validation

For every structured motif provided, we provide scores assigned by HMMpair and RNAPhylo ^2^. For every cluster of structured RNAs, we show the best p-value assigned to the alignments via RNAcode ^13^. We provide the number of instances the structure appears in HMP as well as its length. Within a 5kb range for all instances of each structured RNA, we count which protein domains often occur near the structured RNA. Additionally, we report on the taxonomic classifications of the contigs that these structured RNAs are predicted from. Finally, we denote which of these structured RNAs overlap other structured RNAs in the opposing strand orientation.

Table S3: RPKM quantifications of structured RNAs

For all structured RNAs predicted by Prokka ^17^ and our novel predictions in the four diverse samples with RNA-Seq signal, we provide where these motifs are located in the metagenomic assemblies and their RNA-Seq RPKM values.

