## Supplementary material for "A combined RNA-Seq and comparative genomics approach identifies 1,085 candidate structured RNAs expressed in human microbiomes": RNA Structures

### SampleAMeta_10_1

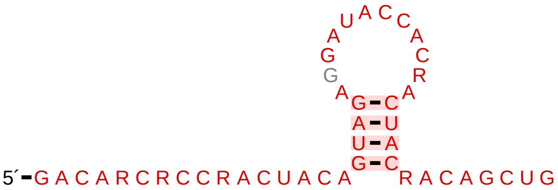

### SampleAMeta_100_1

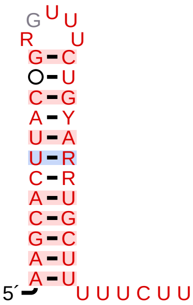

### SampleAMeta_101_1

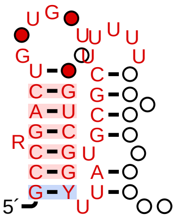

### SampleAMeta_109_1

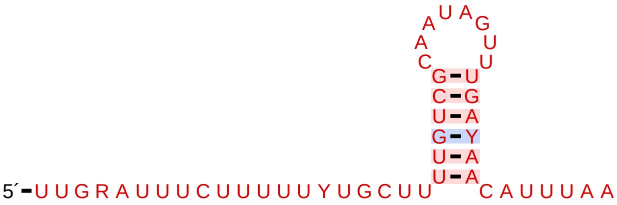

### SampleAMeta_117_1

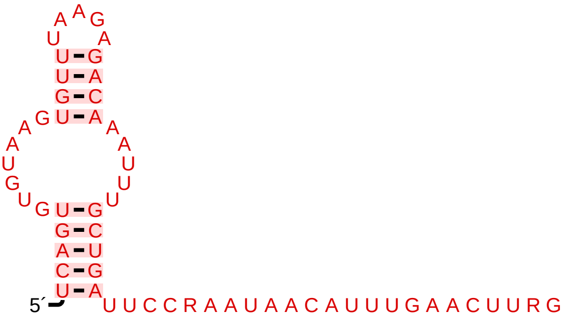

SampleAMeta_118_1

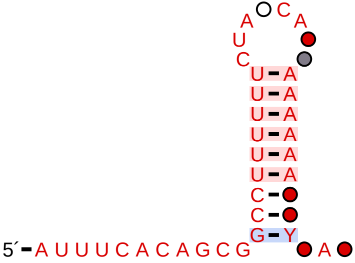

SampleAMeta_119_1

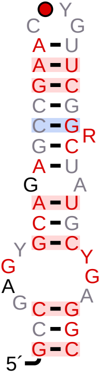

SampleAMeta_120_1

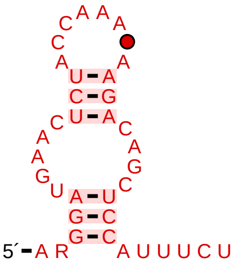

### SampleAMeta_122_1

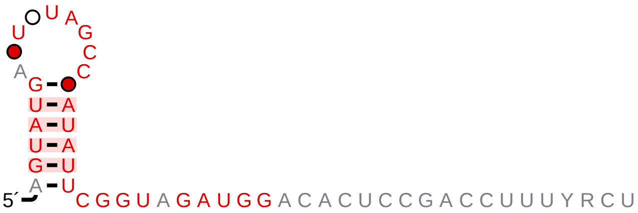

### SampleAMeta_124_1

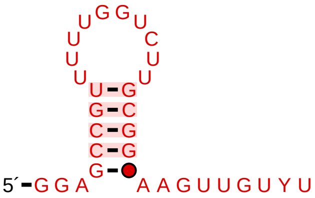

SampleAMeta_125_1

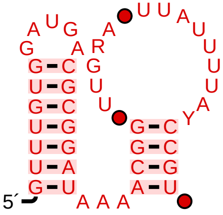

### SampleAMeta_126_1

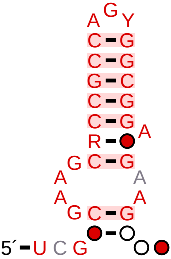

### SampleAMeta_127_1

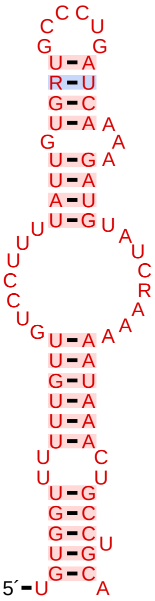

### SampleAMeta_133_1

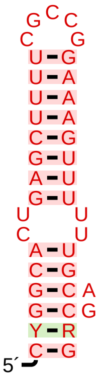

### SampleAMeta_134_1

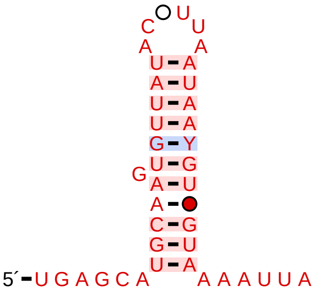

SampleAMeta_143_1

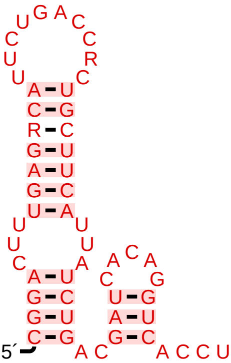

SampleAMeta_144_1

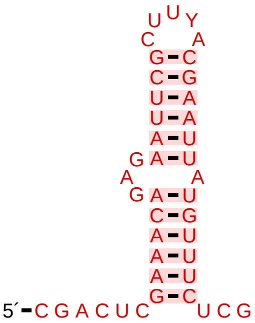

### SampleAMeta_145_1

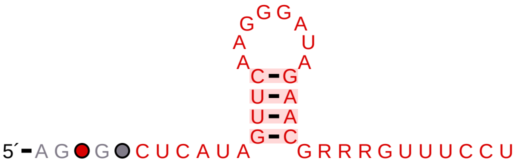

### SampleAMeta_148_1

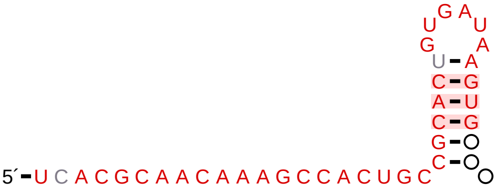

SampleAMeta_149_1

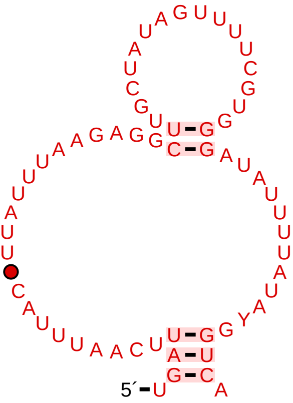

### SampleAMeta_15_1

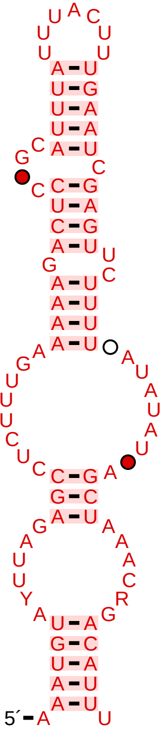

### SampleAMeta_154_1

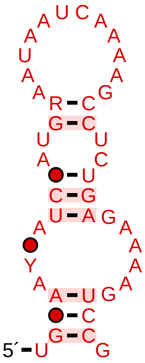

### SampleAMeta_155_1

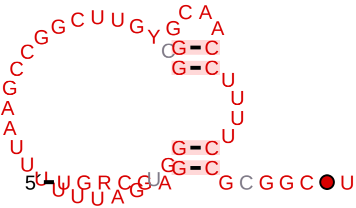

### SampleAMeta_156_1

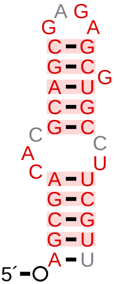

### SampleAMeta_160_1

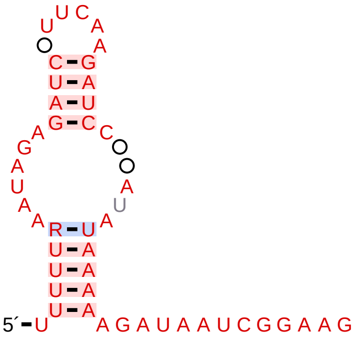

SampleAMeta_161_1

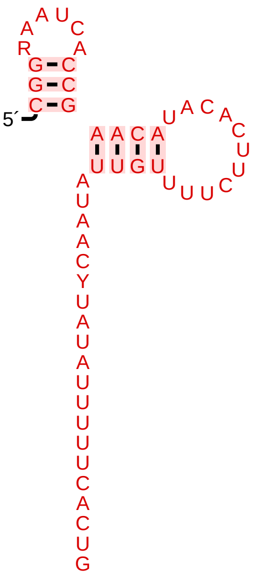

### SampleAMeta_168_1

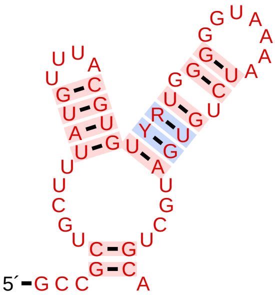

SampleAMeta_169_1

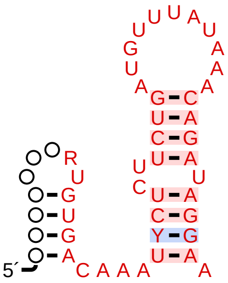

### SampleAMeta_170_1

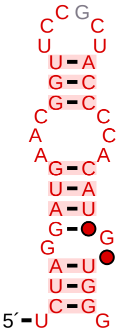

SampleAMeta_182_1

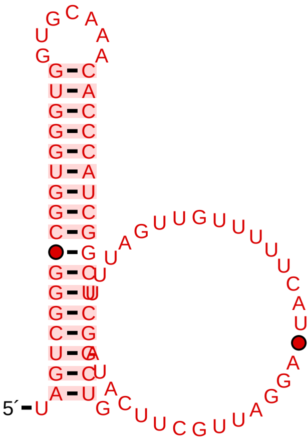

### SampleAMeta_184_1

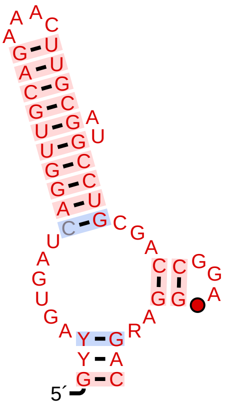

SampleAMeta_186_1

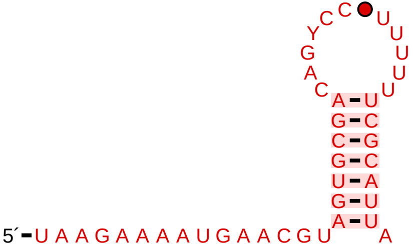

### SampleAMeta_187_1

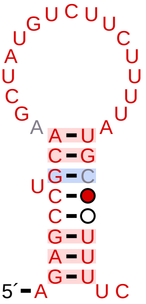

### SampleAMeta_19_1

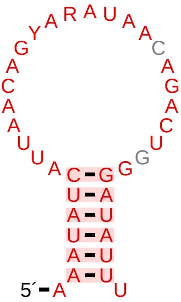

### SampleAMeta_194_1

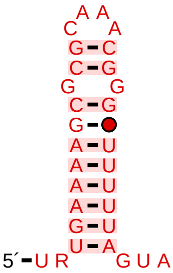

### SampleAMeta_195_1

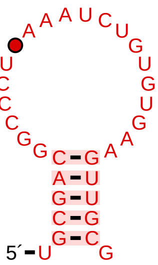

### SampleAMeta_2_1

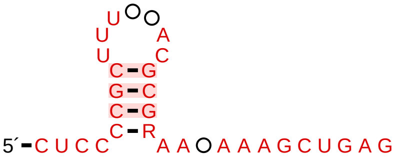

### SampleAMeta_202_1

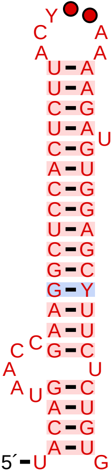

SampleAMeta_203_1

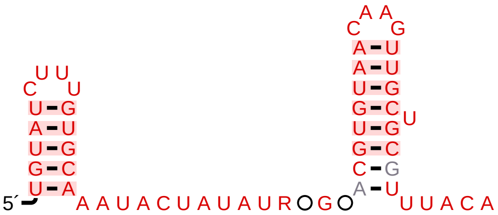

SampleAMeta_206_1

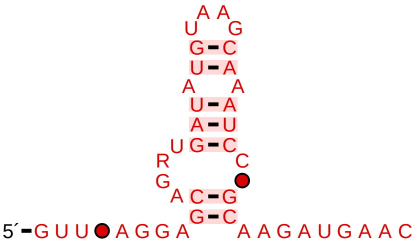

### SampleAMeta_207_1

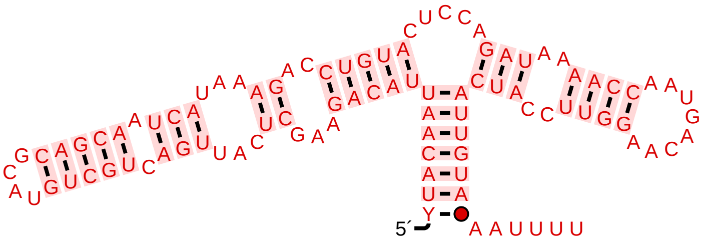

### SampleAMeta_211_1

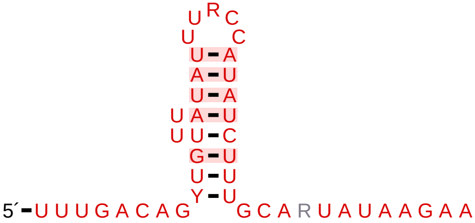

SampleAMeta_212_1

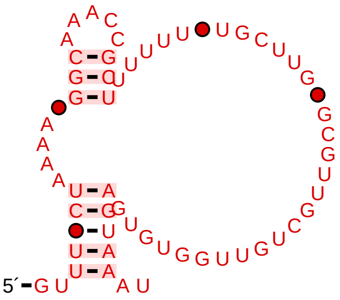

### SampleAMeta_213_1

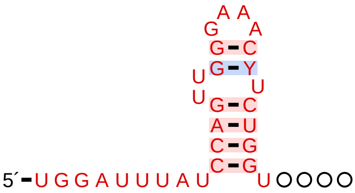

### SampleAMeta_216_1

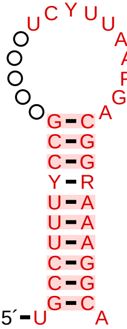

### SampleAMeta_217_1

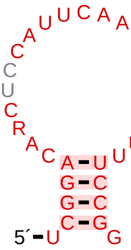

### SampleAMeta_221_1

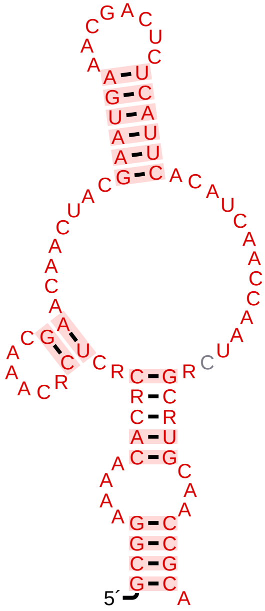

### SampleAMeta_222_1

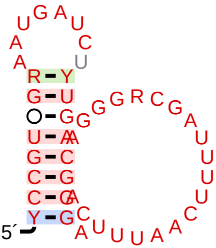

SampleAMeta_225_1

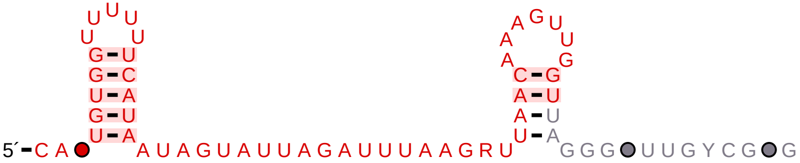

SampleAMeta_226_1

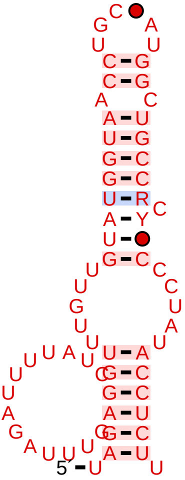

### SampleAMeta_227_1

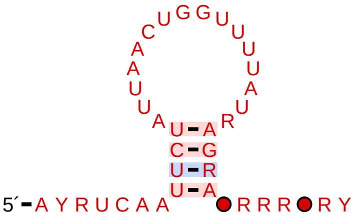

### SampleAMeta_228_1

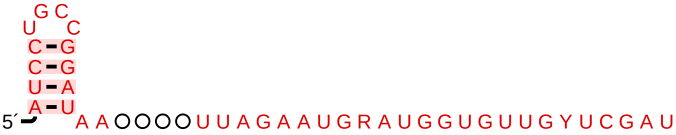

### SampleAMeta_231_1

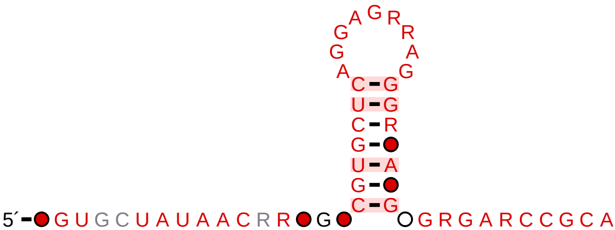

SampleAMeta_234_1

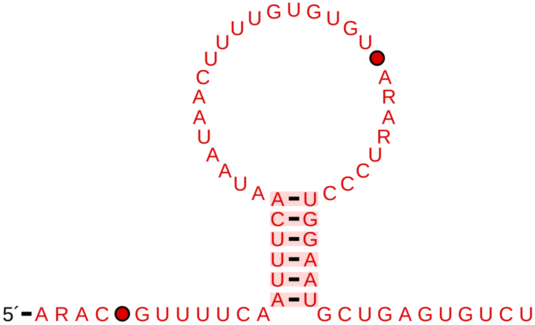

### SampleAMeta_235_1

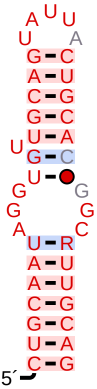

### SampleAMeta_24_1

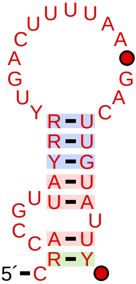

### SampleAMeta_241_1

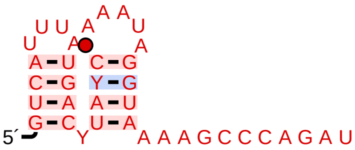

SampleAMeta_242_1

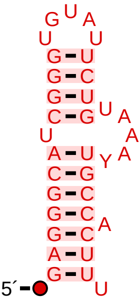

### SampleAMeta_243_1

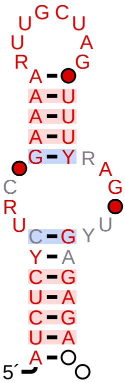

### SampleAMeta_25_1

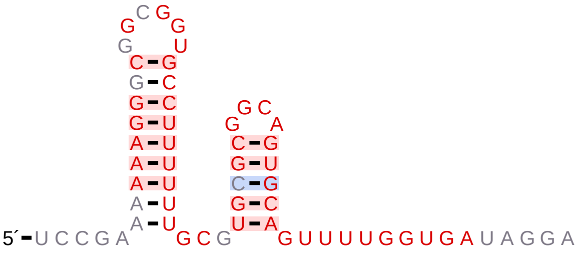

### SampleAMeta_251_1

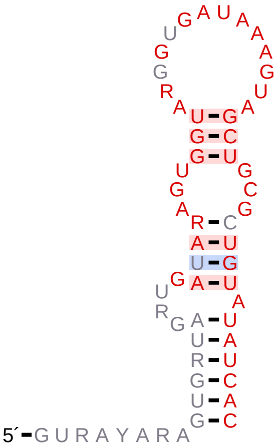

### SampleAMeta_252_1

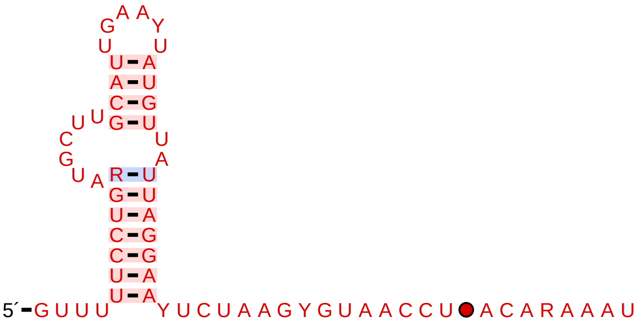

### SampleAMeta_254_1

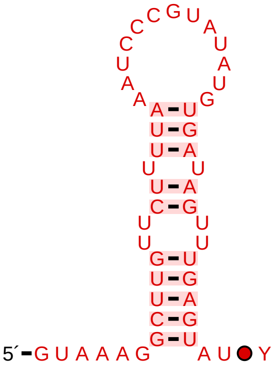

### SampleAMeta_255_1

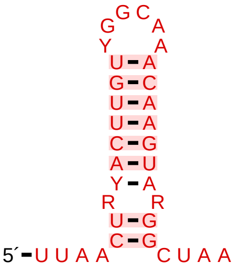

### SampleAMeta_257_1

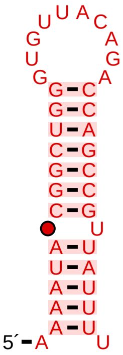

### SampleAMeta_258_1

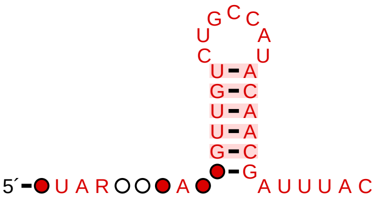

### SampleAMeta_259_1

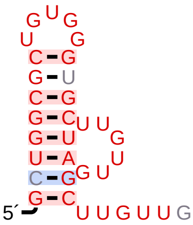

SampleAMeta_260_1

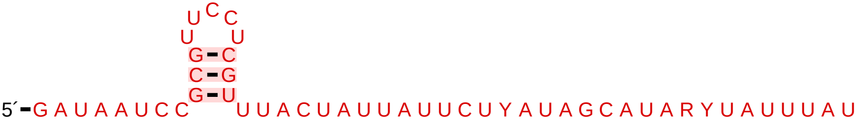

### SampleAMeta_261_1

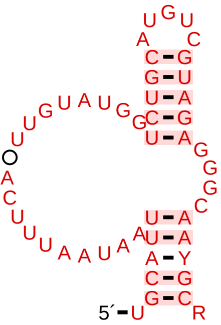

### SampleAMeta_264_1

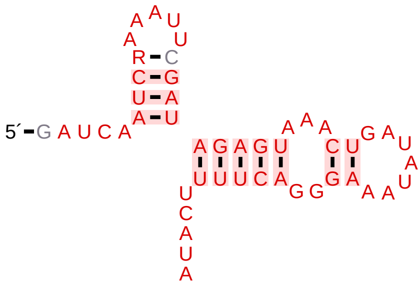

### SampleAMeta_266_1

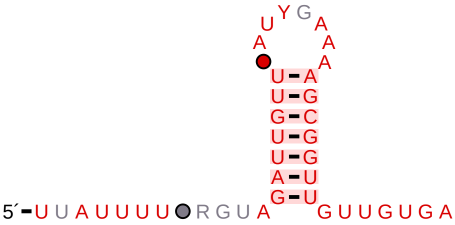

### SampleAMeta_267_1

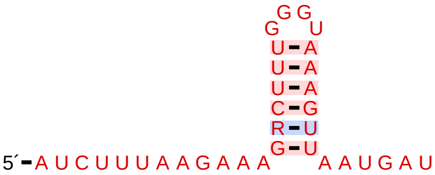

### SampleAMeta_268_1

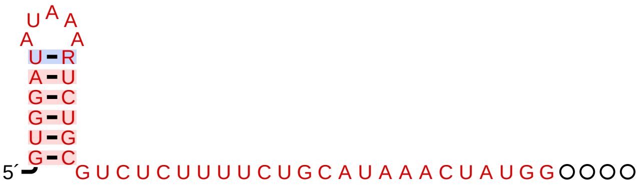

SampleAMeta_269_1

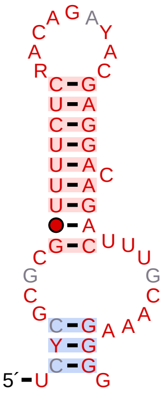

### SampleAMeta_270_1

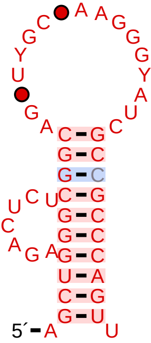

### SampleAMeta_272_1

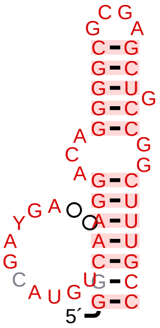

### SampleAMeta_274_1

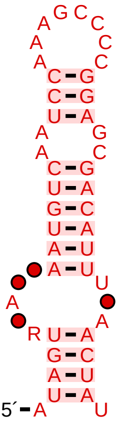

### SampleAMeta_28_1

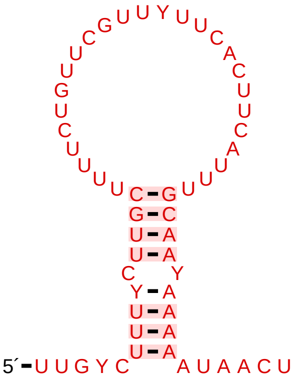

### SampleAMeta_280_1

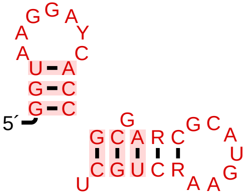

SampleAMeta_284_1

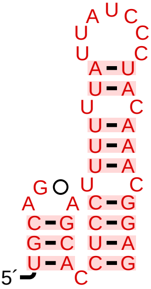

### SampleAMeta_289_1

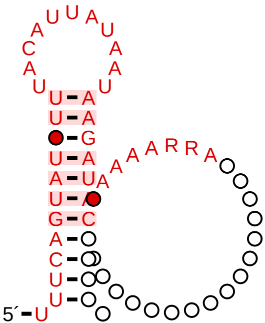

### SampleAMeta_29_1

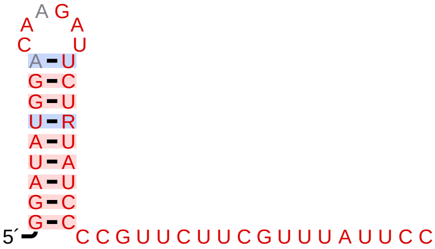

### SampleAMeta_290_1

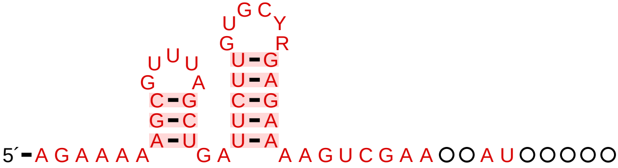

### SampleAMeta_294_1

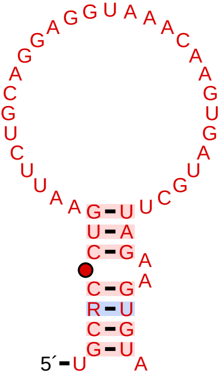

### SampleAMeta_298_1

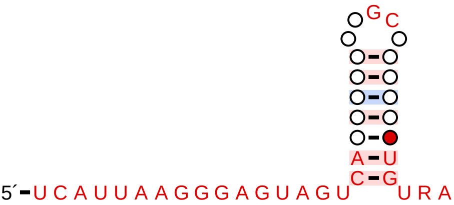

### SampleAMeta_305_1

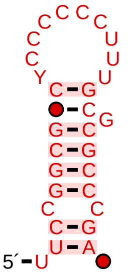

### SampleAMeta_306_1

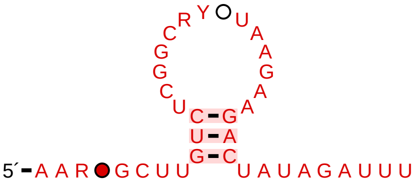

### SampleAMeta_307_1

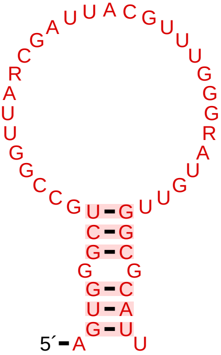

SampleAMeta_310_1

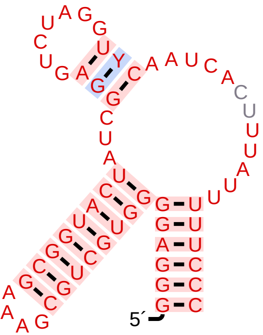

### SampleAMeta_314_1

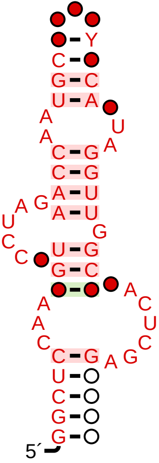

### SampleAMeta_315_1

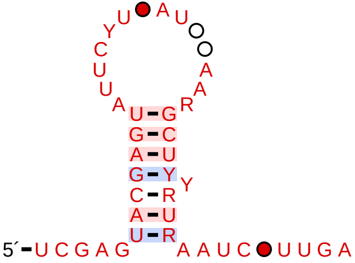

### SampleAMeta_317_1

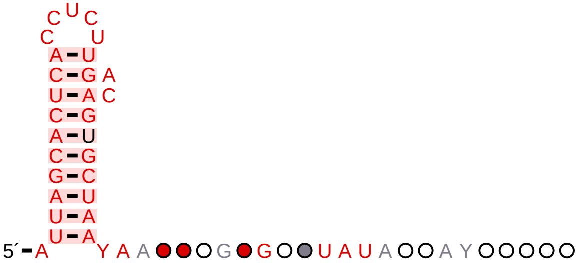

### SampleAMeta_318_1

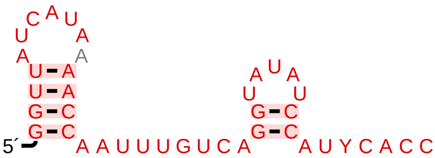

### SampleAMeta_321_1

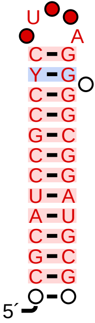

SampleAMeta_322_1

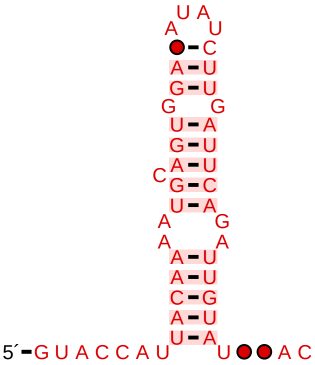

### SampleAMeta_325_1

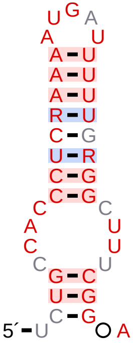

### SampleAMeta_327_1

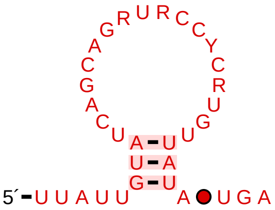

### SampleAMeta_330_1

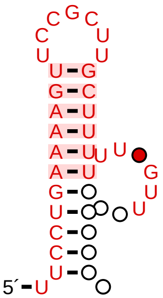

### SampleAMeta_331_1

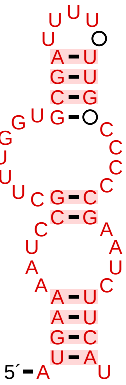

### SampleAMeta_333_1

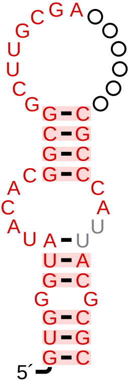

### SampleAMeta_335_1

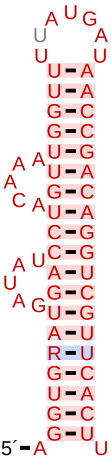

### SampleAMeta_34_1

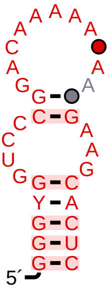

### SampleAMeta_341_1

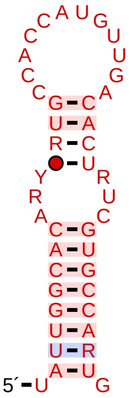

### SampleAMeta_343_1

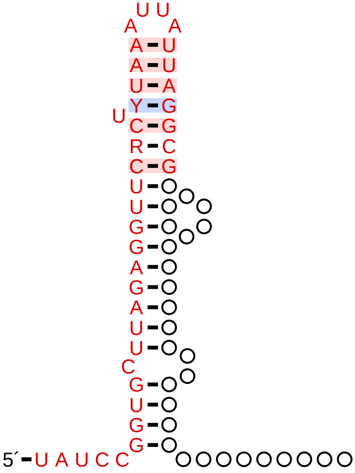

### SampleAMeta_344_1

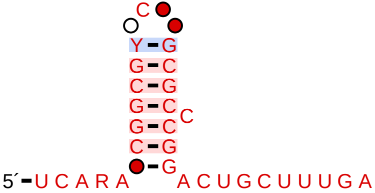

SampleAMeta_346_1

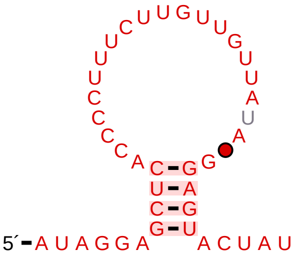

SampleAMeta_347_1

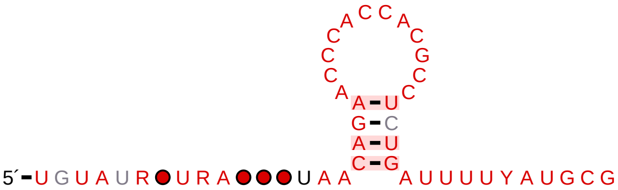

### SampleAMeta_348_1

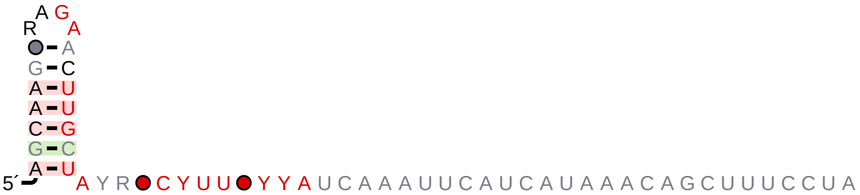

### SampleAMeta_349_1

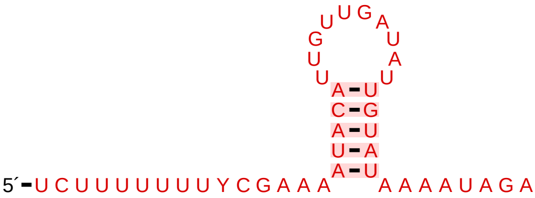

### SampleAMeta_361_1

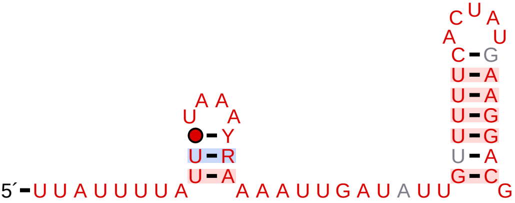

### SampleAMeta_362_1

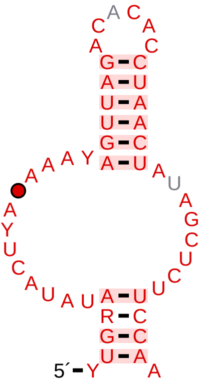

SampleAMeta_365_1

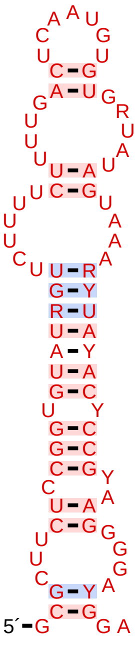

### SampleAMeta_368_1

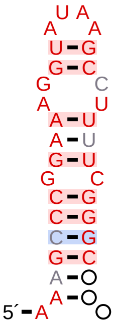

### SampleAMeta_37_1

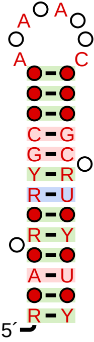

### SampleAMeta_370_1

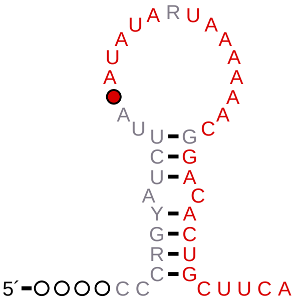

### SampleAMeta_371_1

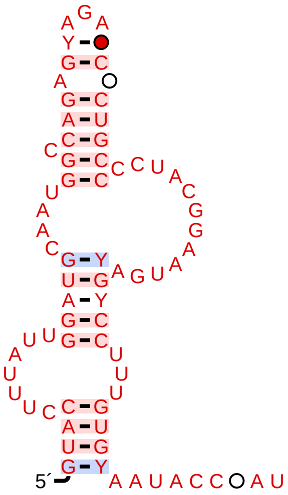

### SampleAMeta_374_1

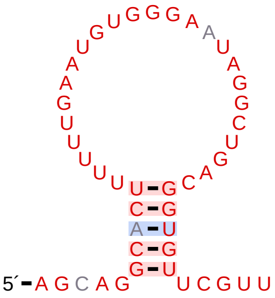

### SampleAMeta_375_1

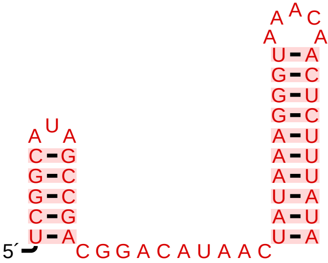

### SampleAMeta_381_1

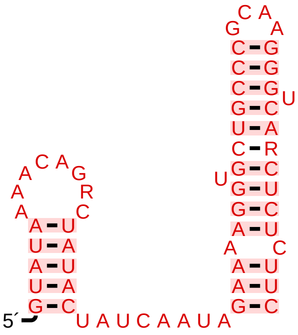

### SampleAMeta_382_1

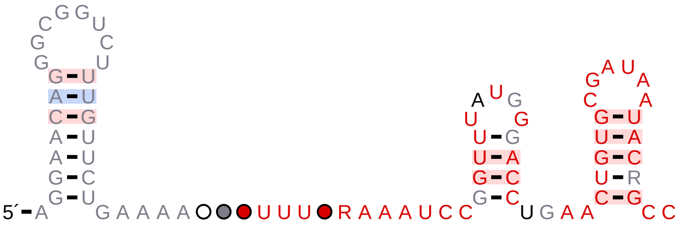

### SampleAMeta_384_1

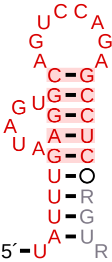

### SampleAMeta_394_1

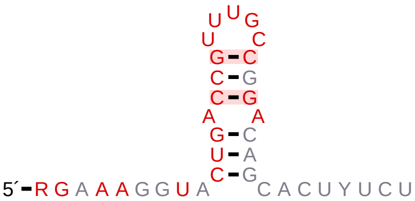

SampleAMeta_396_1

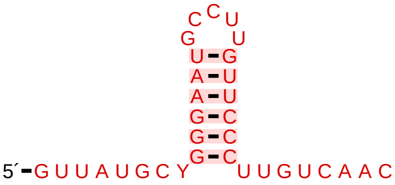

SampleAMeta_397_1

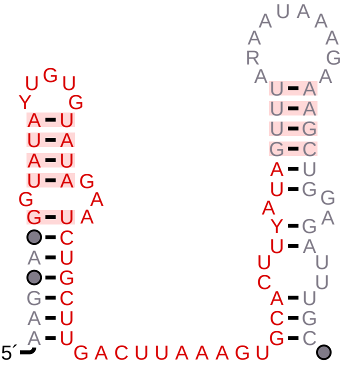

SampleAMeta_398_1

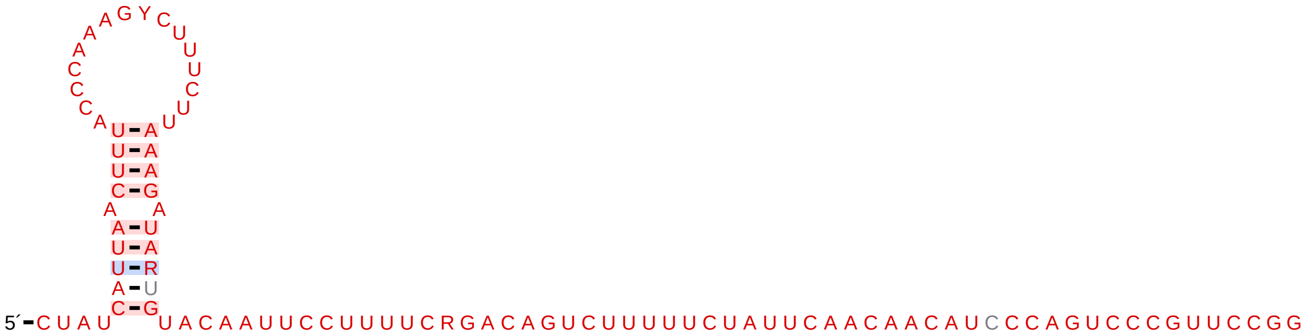

SampleAMeta_399_1

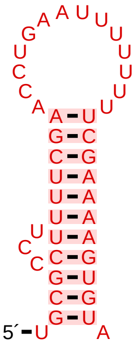

### SampleAMeta_401_1

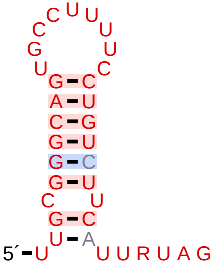

### SampleAMeta_402_1

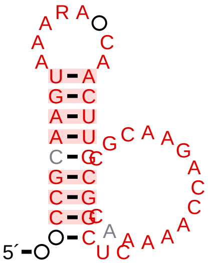

### SampleAMeta_41_1

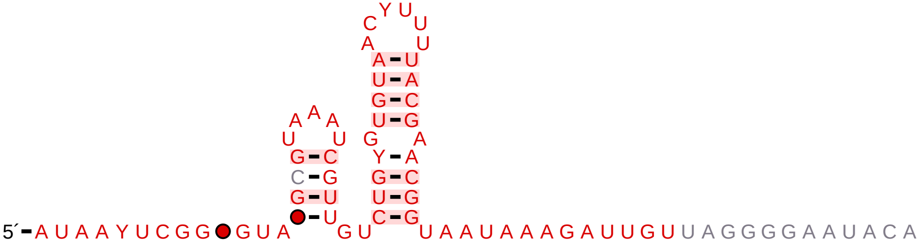

### SampleAMeta_412_1

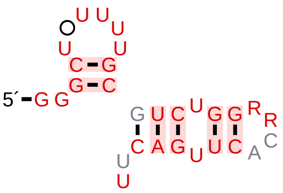

SampleAMeta_413_1

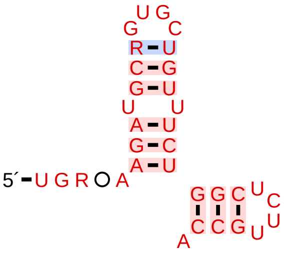

### SampleAMeta_414_1

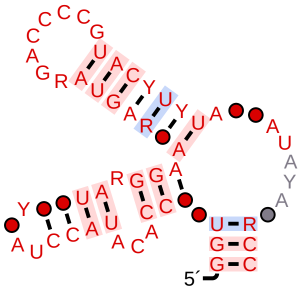

### SampleAMeta_416_1

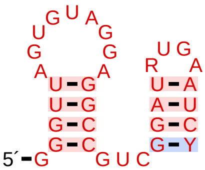

### SampleAMeta_418_1

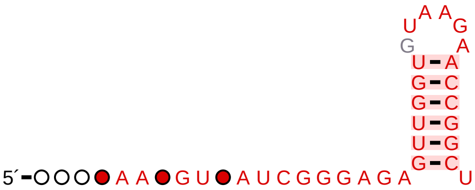

### SampleAMeta_42_1

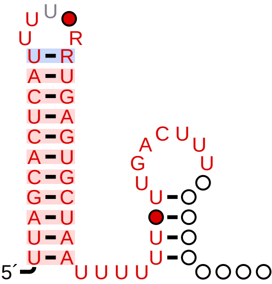

### SampleAMeta_420_1

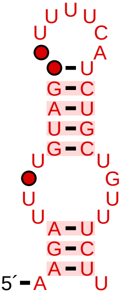

### SampleAMeta_421_1

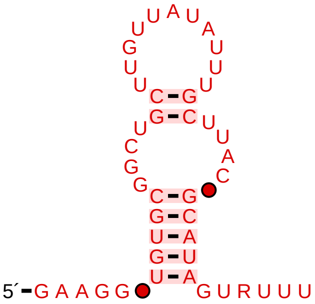

### SampleAMeta_424_1

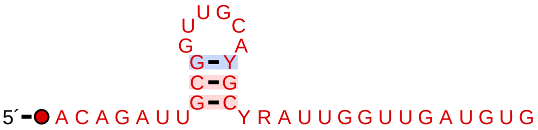

### SampleAMeta_43_1

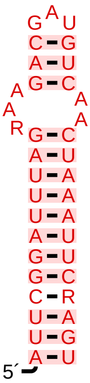

### SampleAMeta_430_1

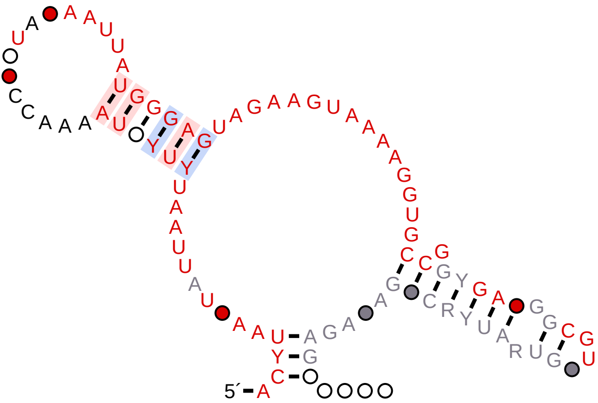

### SampleAMeta_433_1

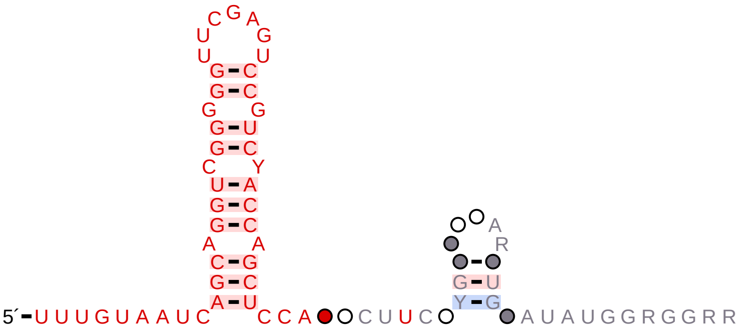

### SampleAMeta_436_1

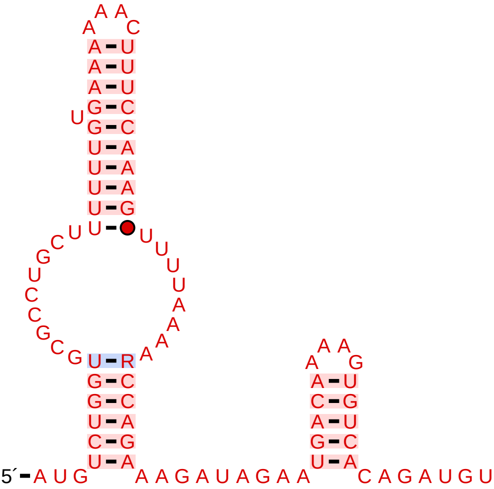

### SampleAMeta_437_1

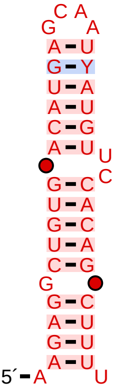

### SampleAMeta_443_1

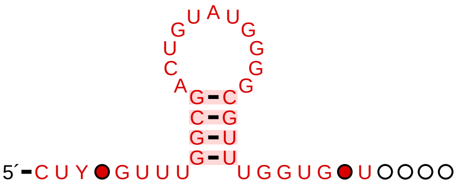

### SampleAMeta_444_1

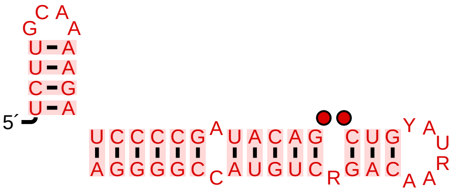

SampleAMeta_445_1

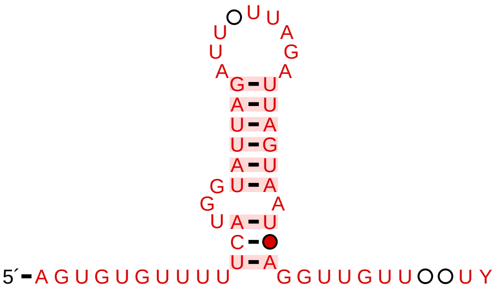

### SampleAMeta_448_1

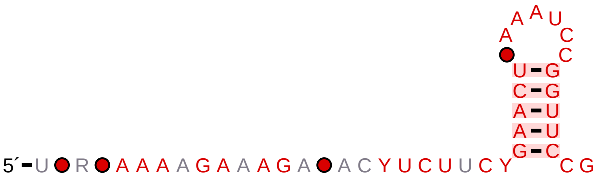

### SampleAMeta_453_1

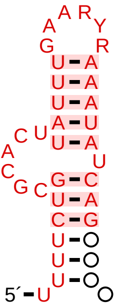

### SampleAMeta_454_1

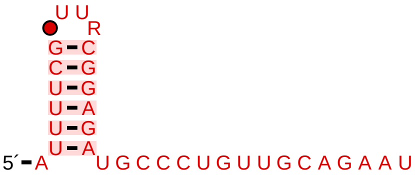

### SampleAMeta_458_1

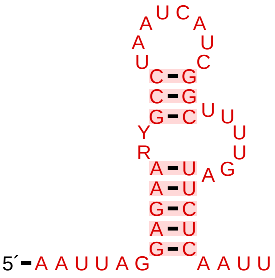

SampleAMeta_46_1

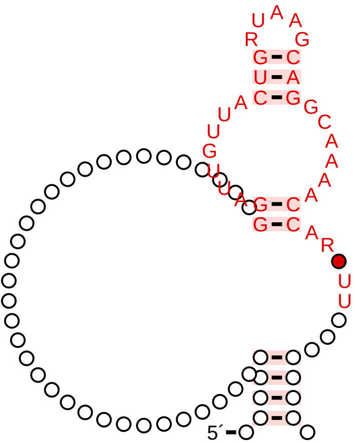

SampleAMeta_460_1

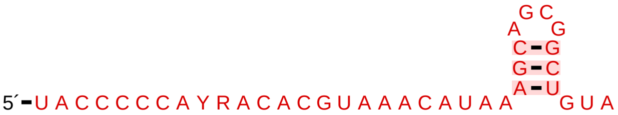

### SampleAMeta_466_1

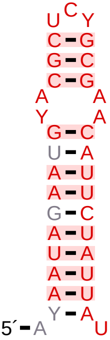

### SampleAMeta_468_1

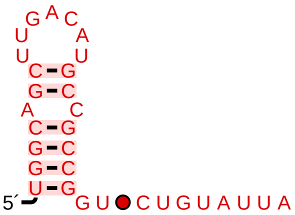

### SampleAMeta_47_1

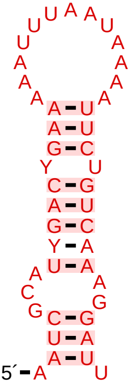

### SampleAMeta_471_1

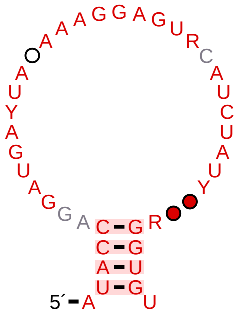

### SampleAMeta_477_1

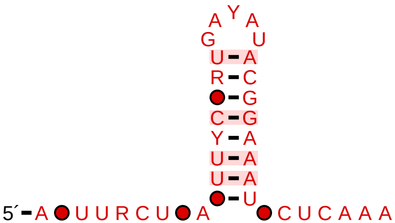

### SampleAMeta_479_1

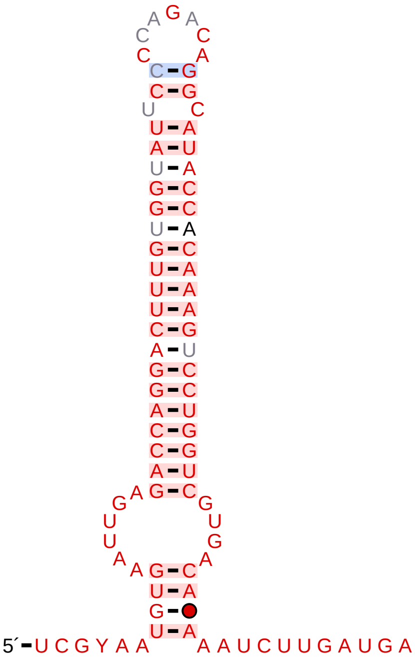

### SampleAMeta_481_1

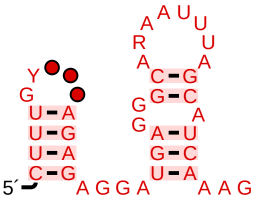

### SampleAMeta_483_1

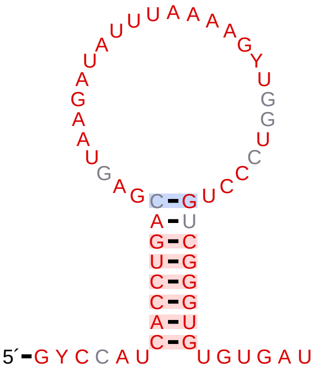

### SampleAMeta_484_1

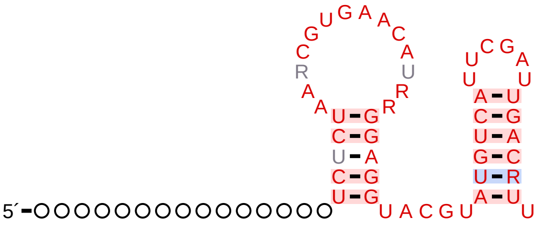

SampleAMeta_485_1

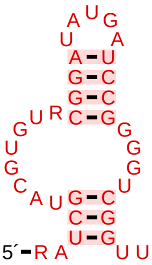

SampleAMeta_489_1

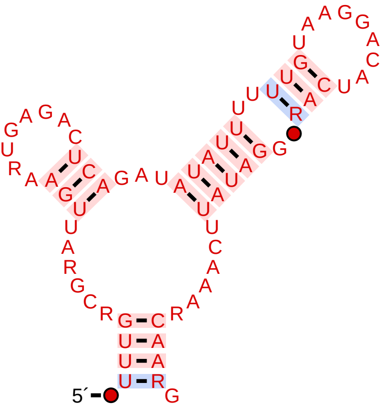

### SampleAMeta_49_1

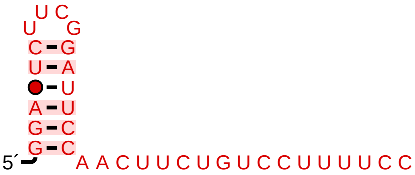

### SampleAMeta_494_1

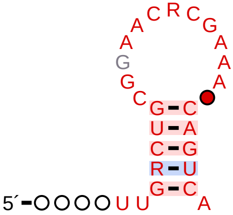

### SampleAMeta_495_1

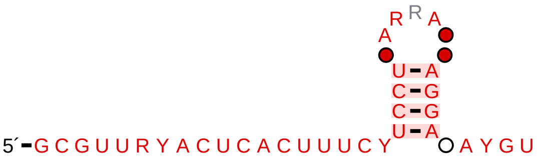

SampleAMeta_499_1

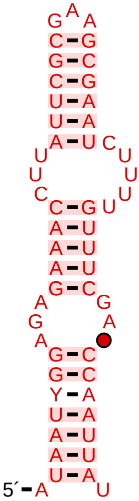

SampleAMeta_502_1

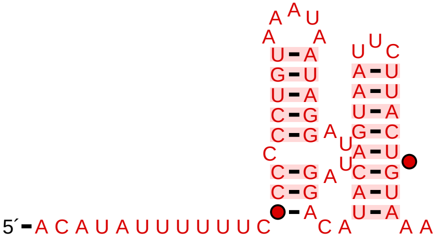

SampleAMeta_503_1

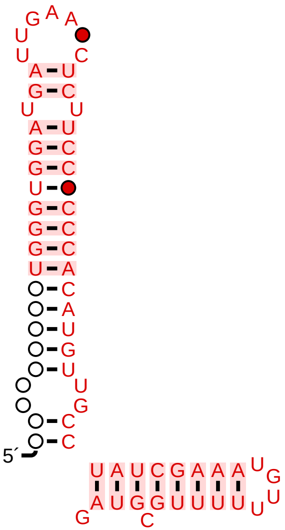

### SampleAMeta_508_1

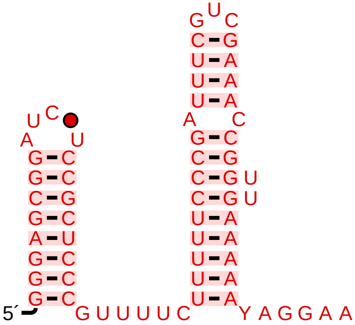

### SampleAMeta_509_1

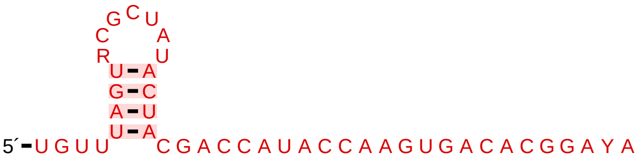

### SampleAMeta_514_1

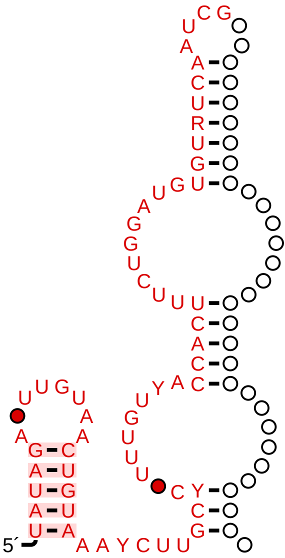

### SampleAMeta_515_1

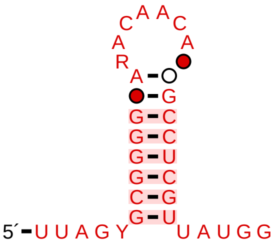

### SampleAMeta_517_1

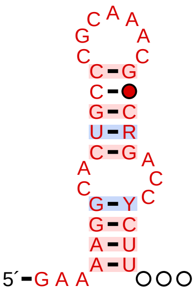

### SampleAMeta_519_1

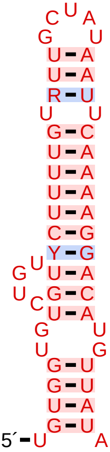

### SampleAMeta_52_1

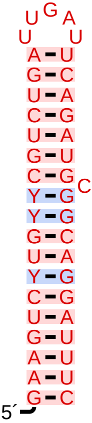

### SampleAMeta_521_1

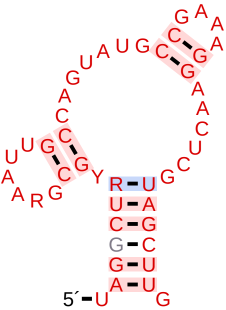

### SampleAMeta_522_1

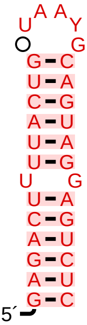

### SampleAMeta_525_1

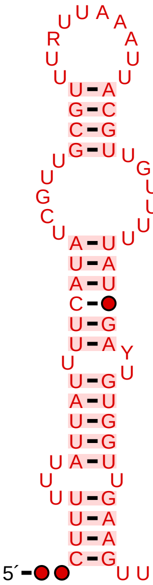

### SampleAMeta_527_1

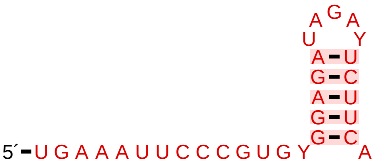

### SampleAMeta_53_1

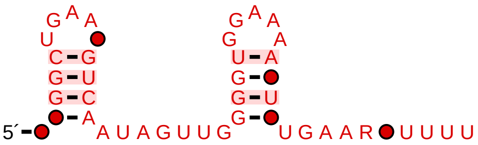

### SampleAMeta_533_1

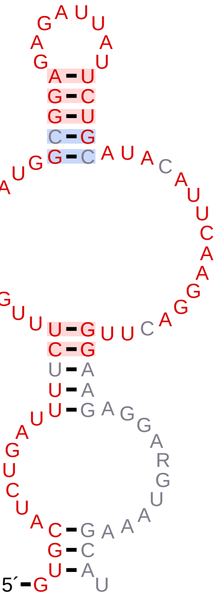

SampleAMeta_535_1

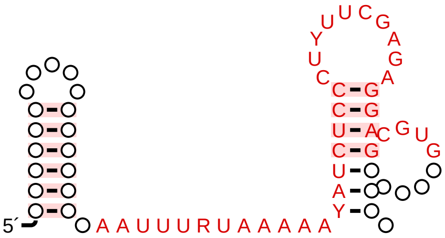

SampleAMeta_537_1

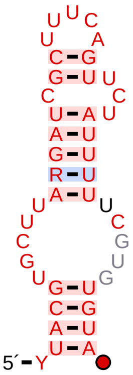

SampleAMeta_54_1

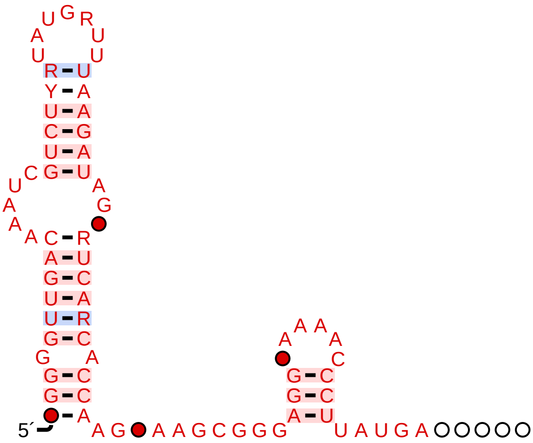

### SampleAMeta_540_1

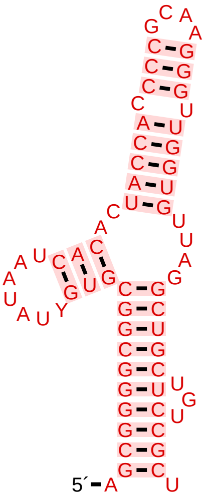

### SampleAMeta_545_1

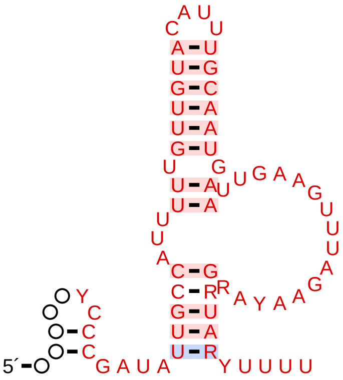

### SampleAMeta_546_1

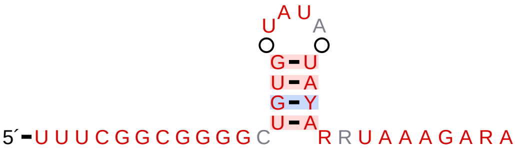

### SampleAMeta_547_1

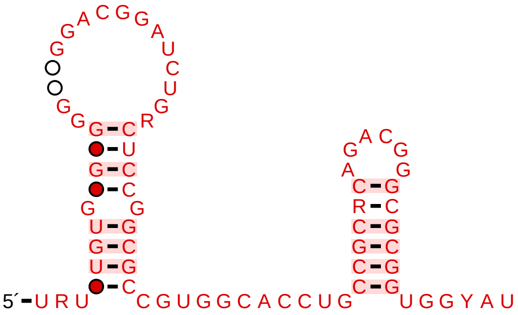

### SampleAMeta_548_1

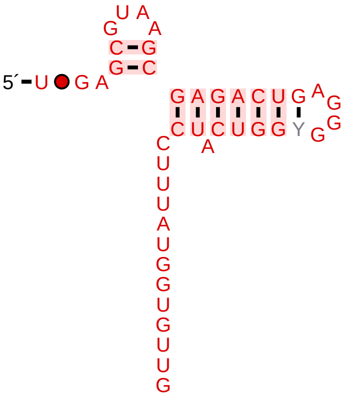

### SampleAMeta_55_1

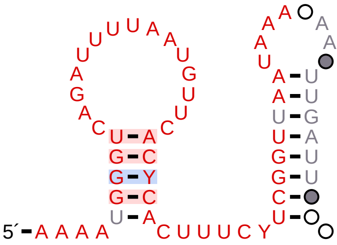

SampleAMeta_553_1

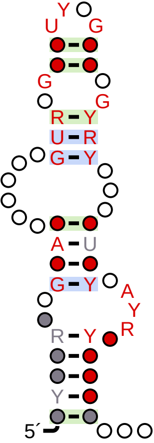

### SampleAMeta_554_1

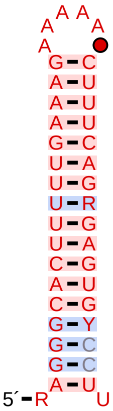

SampleAMeta_555_1

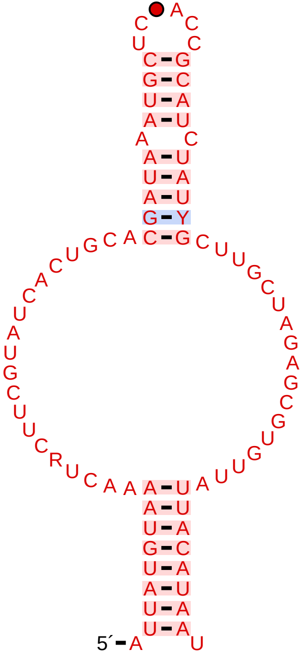

### SampleAMeta_557_1

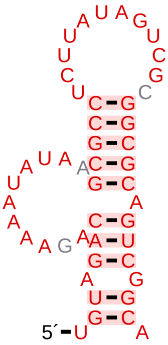

### SampleAMeta_564_1

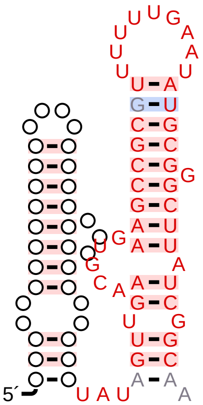

### SampleAMeta_565_1

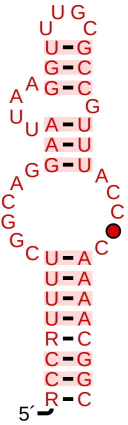

### SampleAMeta_566_1

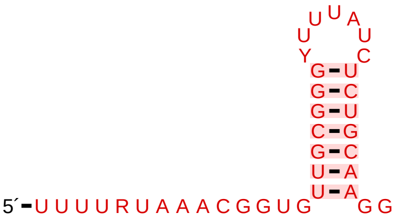

### SampleAMeta_572_1

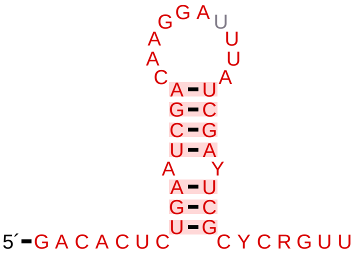

### SampleAMeta_573_1

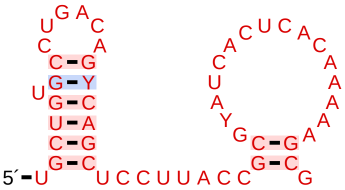

### SampleAMeta_576_1

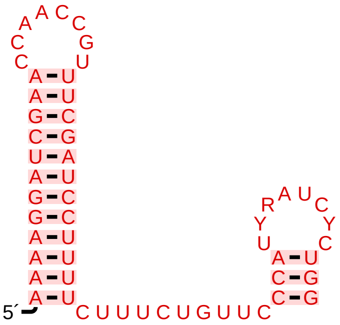

### SampleAMeta_577_1

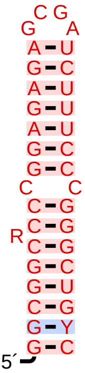

### SampleAMeta_578_1

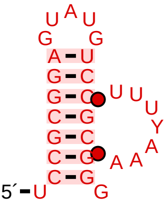

### SampleAMeta_579_1

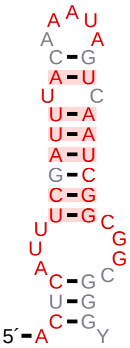

### SampleAMeta_580_1

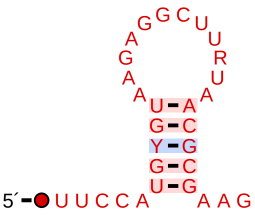

SampleAMeta_588_1

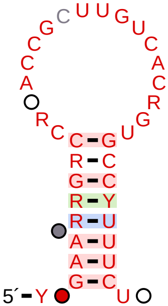

### SampleAMeta_589_1

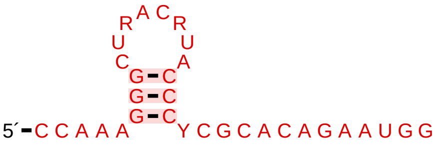

### SampleAMeta_593_1

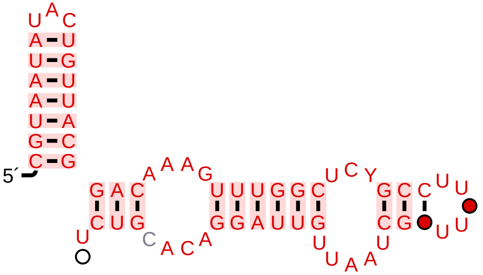

### SampleAMeta_595_1

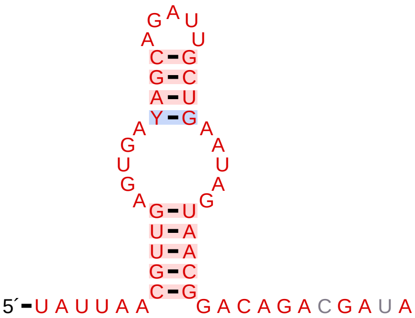

SampleAMeta_6_1

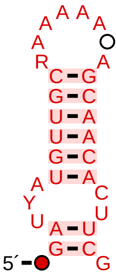

### SampleAMeta_601_1

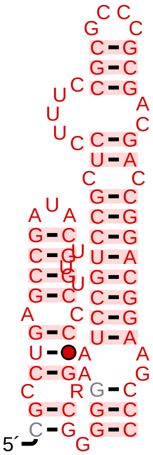

### SampleAMeta_61_1

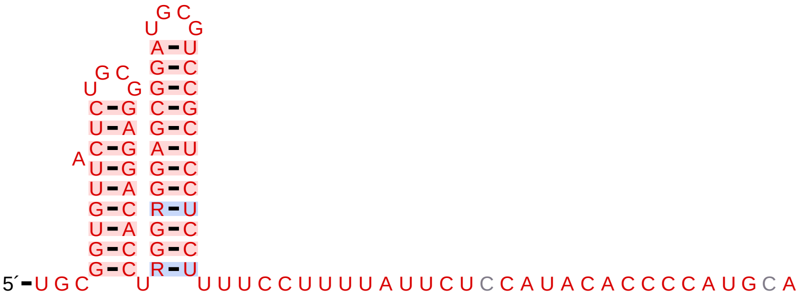

### SampleAMeta_611_1

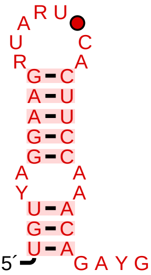

### SampleAMeta_612_1

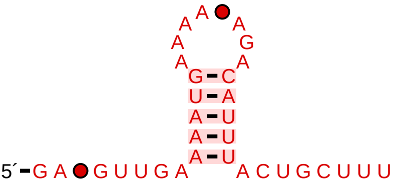

### SampleAMeta_614_1

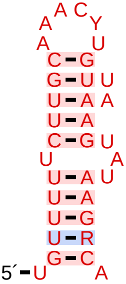

### SampleAMeta_618_1

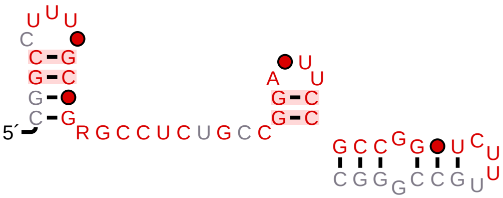

### SampleAMeta_620_1

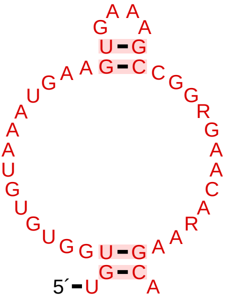

### SampleAMeta_623_1

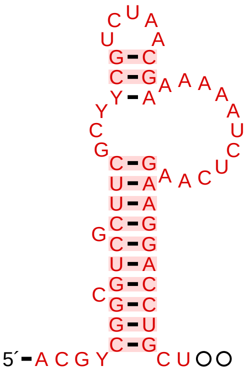

SampleAMeta_625_1

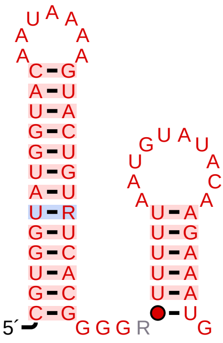

### SampleAMeta_626_1

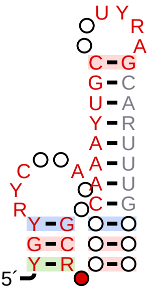

### SampleAMeta_627_1

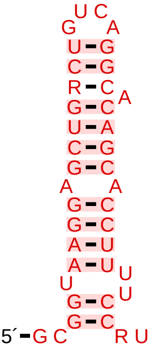

SampleAMeta_628_1

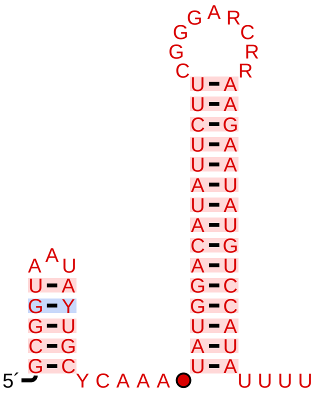

SampleAMeta_630_1

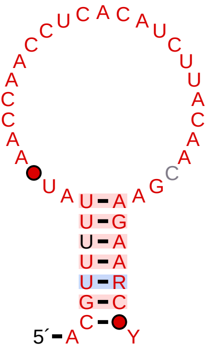

### SampleAMeta_632_1

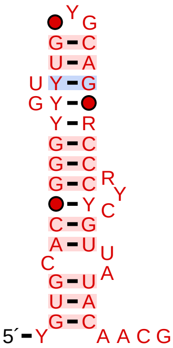

### SampleAMeta_633_1

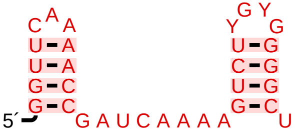

SampleAMeta_636_1

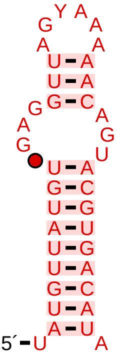

### SampleAMeta_639_1

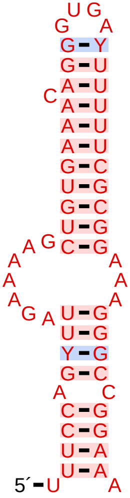

SampleAMeta_642_1

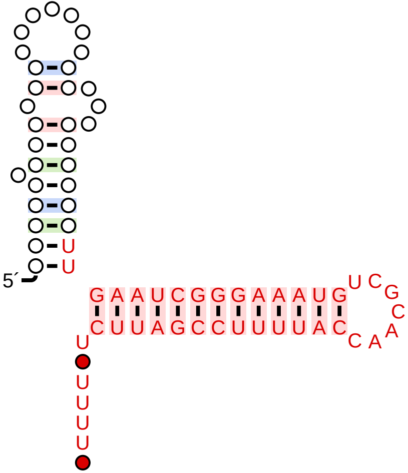

### SampleAMeta_643_1

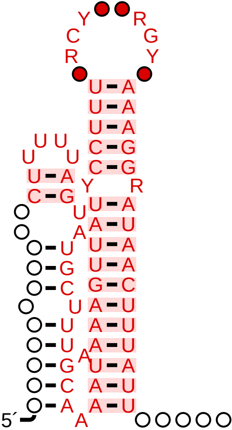

SampleAMeta_644_1

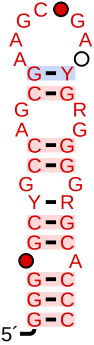

### SampleAMeta_646_1

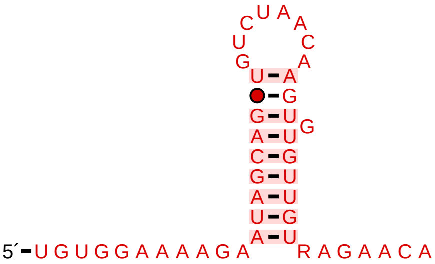

### SampleAMeta_651_1

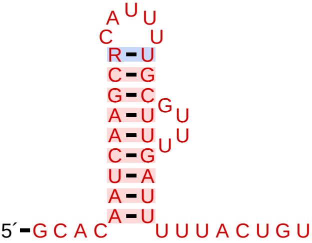

SampleAMeta_653_1

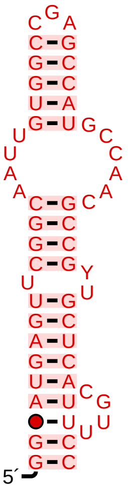

### SampleAMeta_656_1

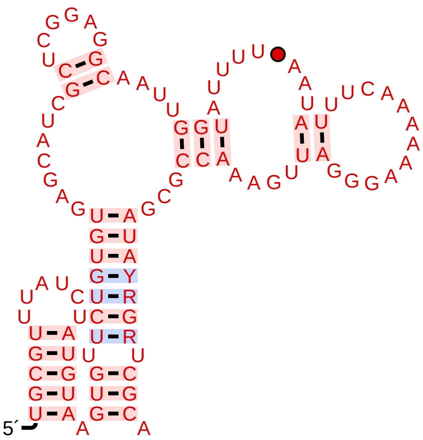

### SampleAMeta_657_1

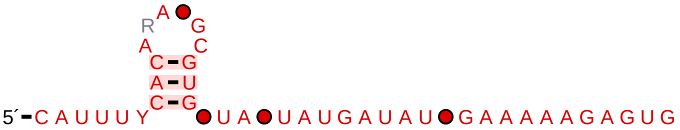

SampleAMeta_662_1

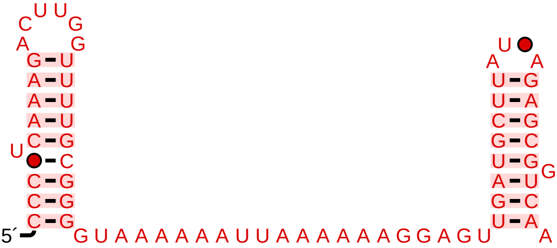

### SampleAMeta_663_1

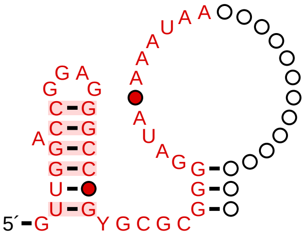

### SampleAMeta_664_1

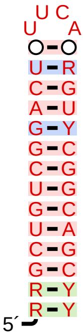

### SampleAMeta_665_1

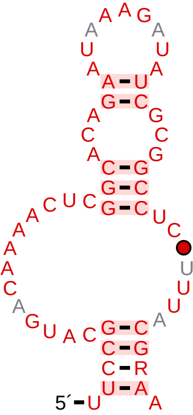

### SampleAMeta_666_1

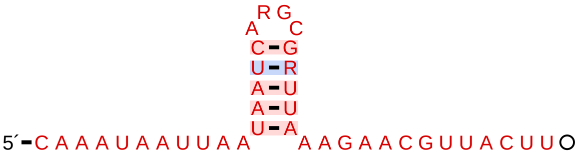

### SampleAMeta_667_1

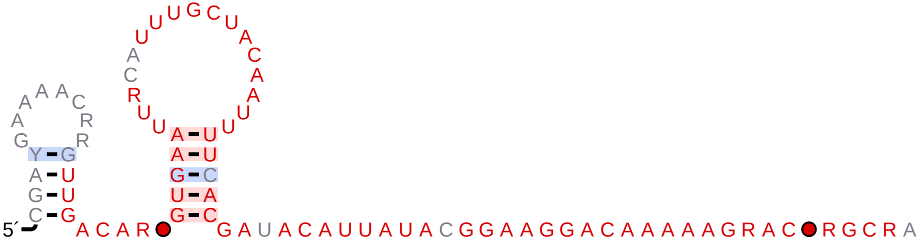

### SampleAMeta_668_1

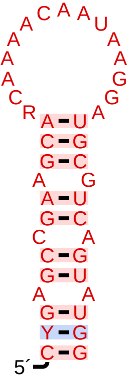

SampleAMeta_669_1

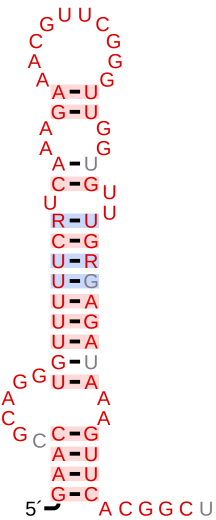

### SampleAMeta_671_1

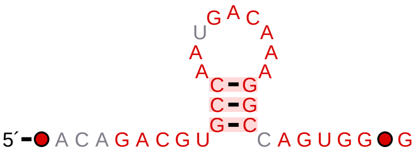

### SampleAMeta_672_1

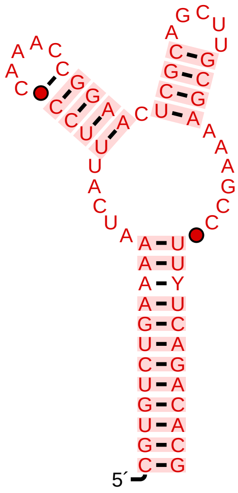

SampleAMeta_676_1

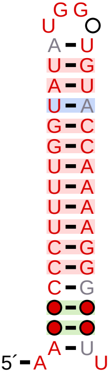

SampleAMeta_677_1

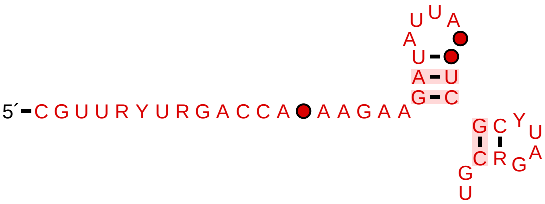

SampleAMeta_680_1

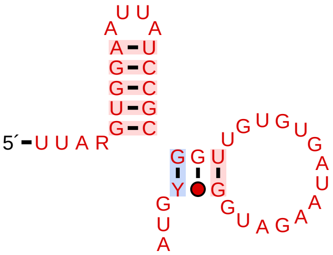

SampleAMeta_685_1

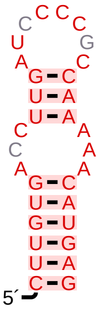

SampleAMeta_688_1

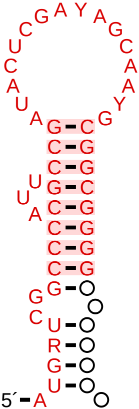

SampleAMeta_689_1

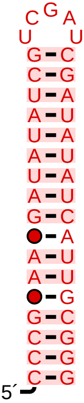

### SampleAMeta_7_1

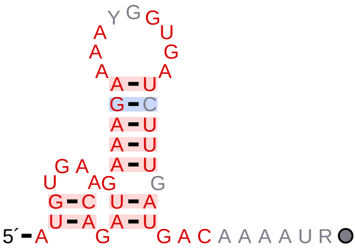

### SampleAMeta_71_1

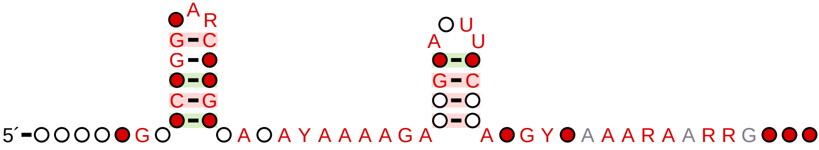

### SampleAMeta_73_1

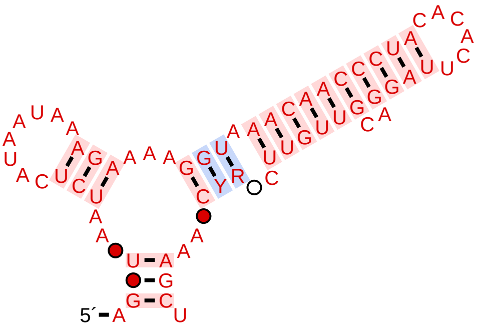

### SampleAMeta_76_1

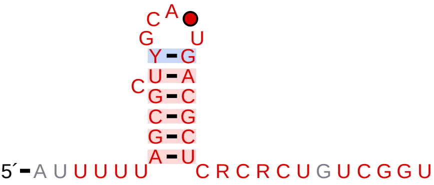

### SampleAMeta_80_1

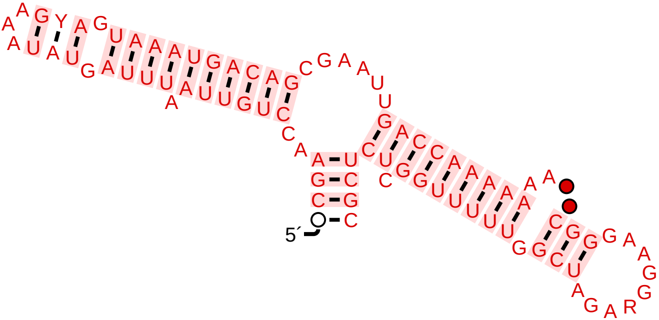

### SampleAMeta_82_1

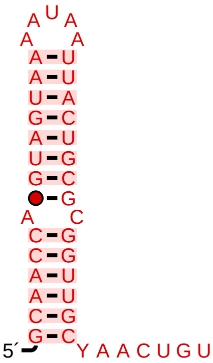

### SampleAMeta_88_1

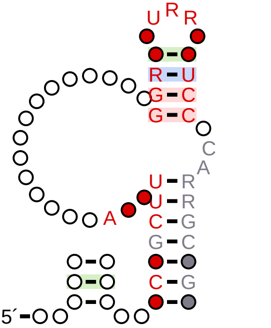

SampleAMeta_89_1

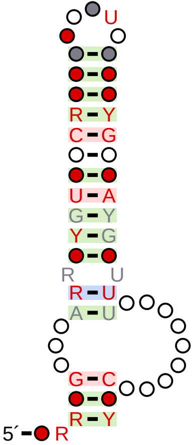

SampleAMeta_92_1

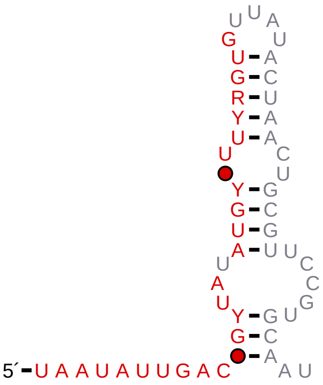

### SampleAMeta_94_1

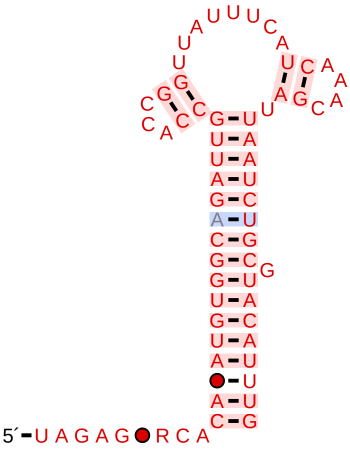

SampleAMeta_95_1

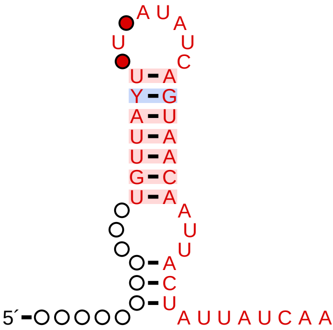

### SampleAMeta_97_1

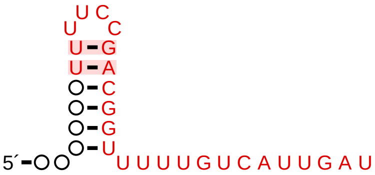

SampleAMeta_99_1

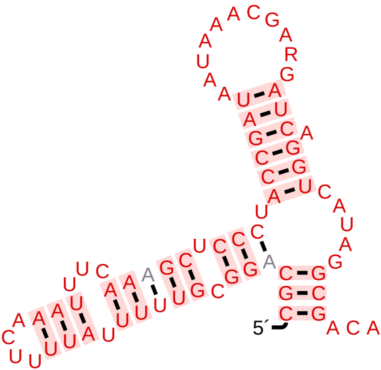

SampleBMeta_1_1

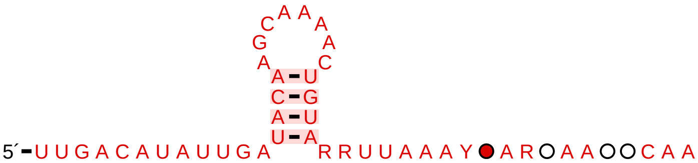

### SampleBMeta_101_1

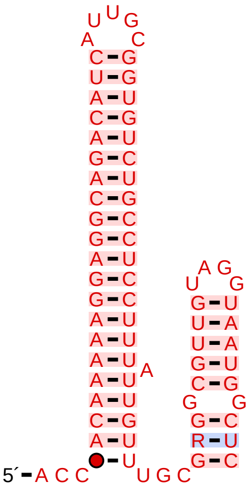

SampleBMeta_102_1

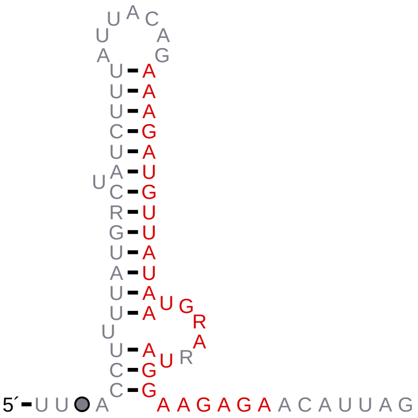

SampleBMeta_104_1

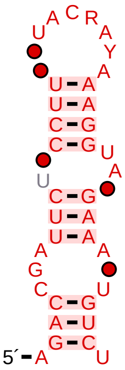

### SampleBMeta_109_1

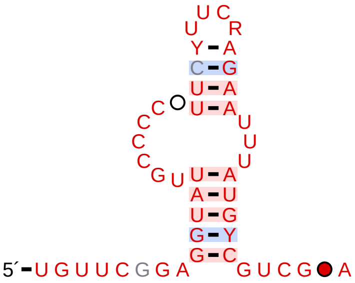

### SampleBMeta_112_1

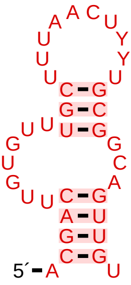

### SampleBMeta_113_1

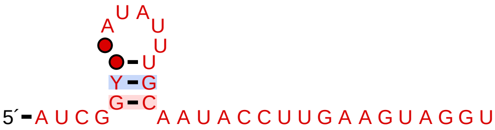

### SampleBMeta_115_1

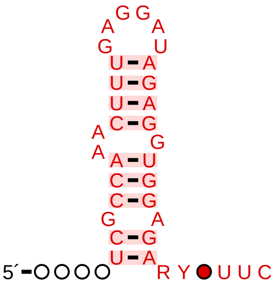

SampleBMeta_116_1

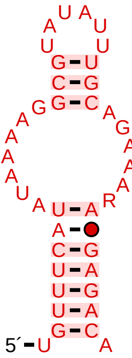

### SampleBMeta_119_1

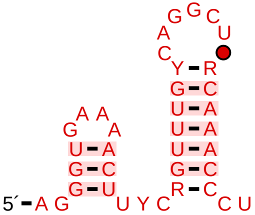

### SampleBMeta_123_1

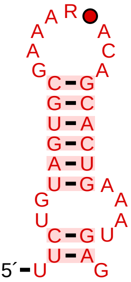

### SampleBMeta_124_1

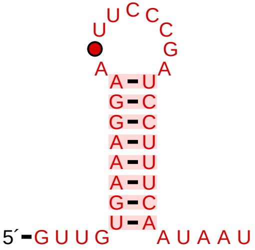

### SampleBMeta_126_1

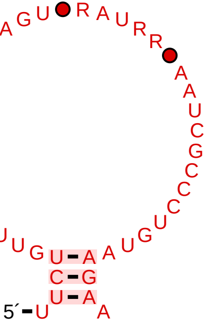

SampleBMeta_127_1

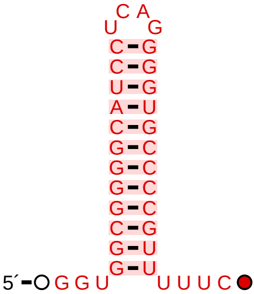

### SampleBMeta_129_1

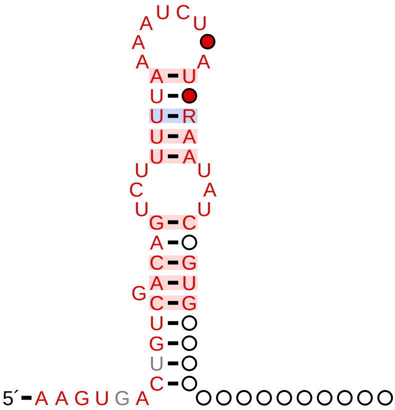

SampleBMeta_131_1

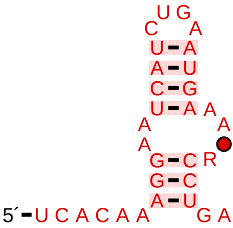

### SampleBMeta_133_1

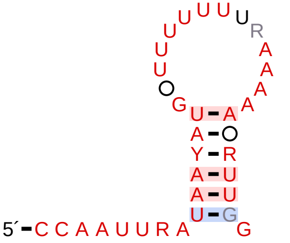

### SampleBMeta_137_1

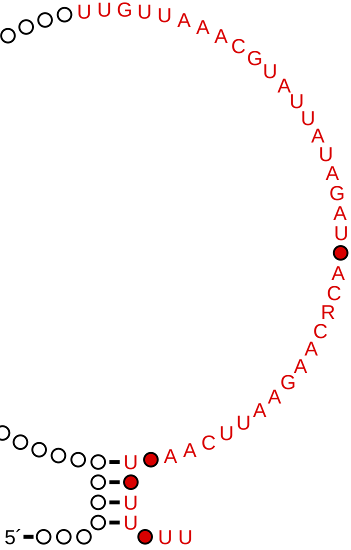

SampleBMeta_139_1

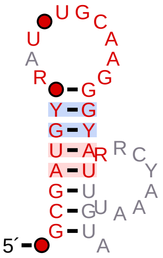

### SampleBMeta_141_1

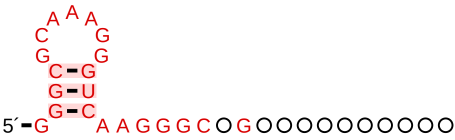

### SampleBMeta_144_1

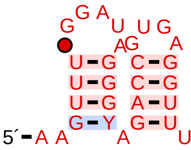

### SampleBMeta_146_1

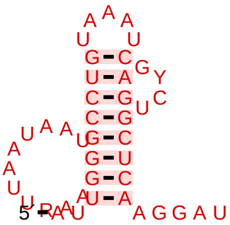

### SampleBMeta_15_1

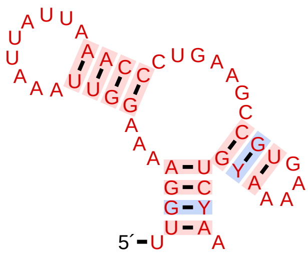

SampleBMeta_150_1

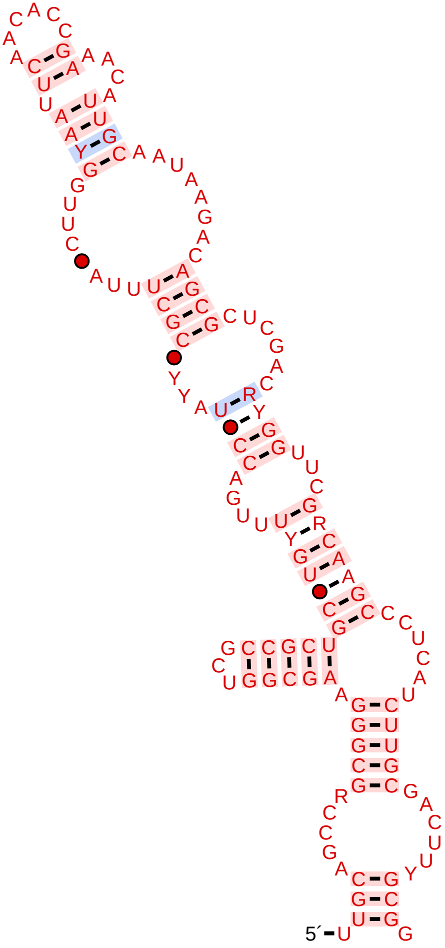

### SampleBMeta_152_1

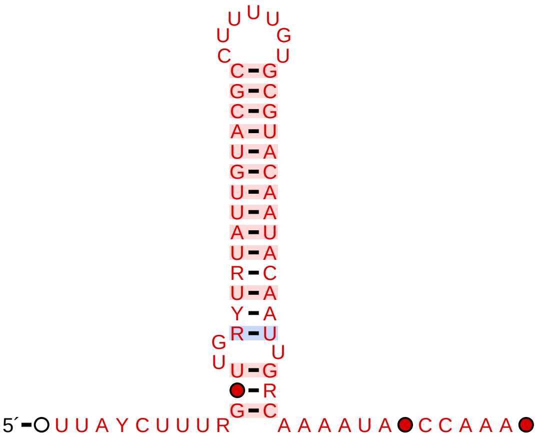

### SampleBMeta_155_1

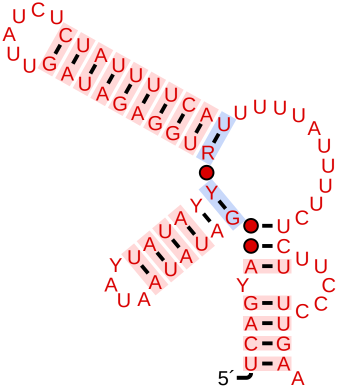

### SampleBMeta_156_1

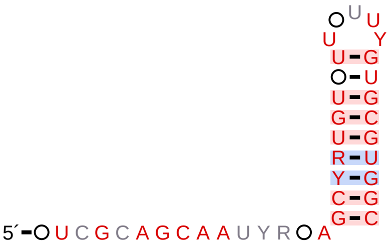

### SampleBMeta_157_1

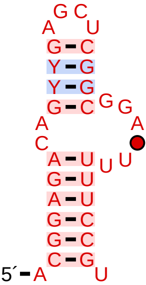

### SampleBMeta_158_1

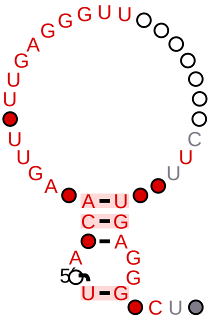

### SampleBMeta_16_1

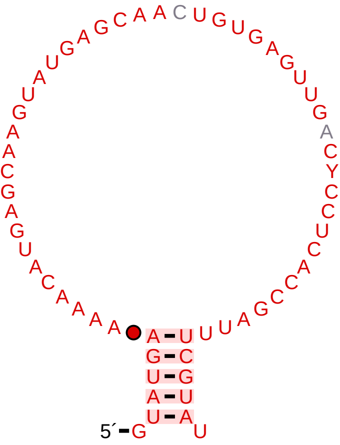

### SampleBMeta_160_1

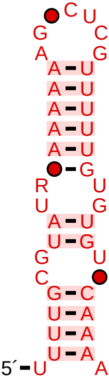

### SampleBMeta_163_1

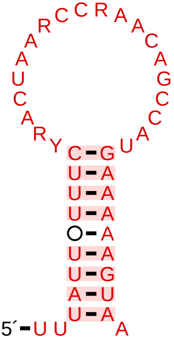

SampleBMeta_165_1

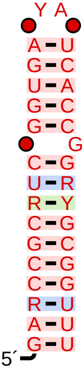

### SampleBMeta_171_1

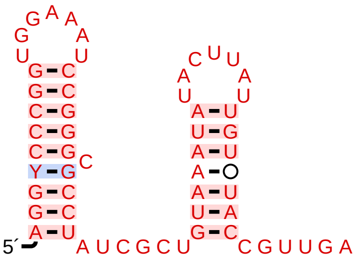

### SampleBMeta_175_1

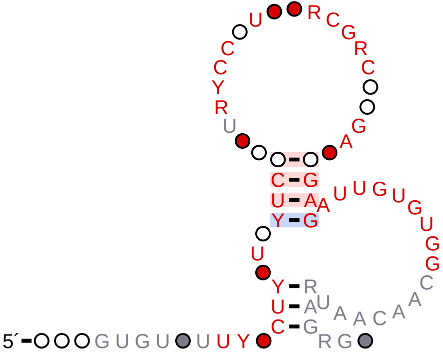

### SampleBMeta_18_1

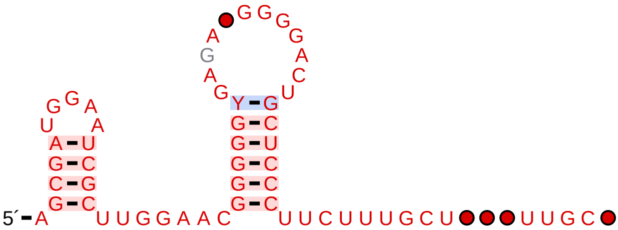

SampleBMeta_183_1

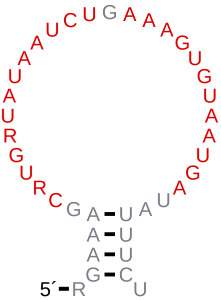

SampleBMeta_184_1

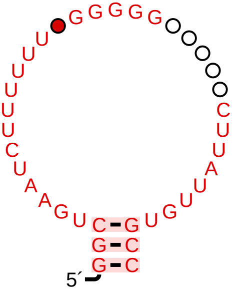

### SampleBMeta_186_1

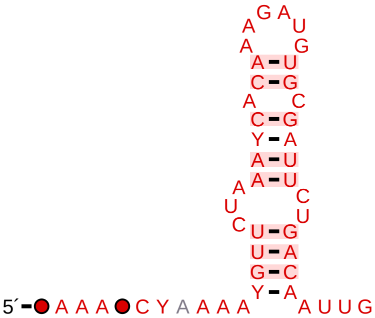

### SampleBMeta_187_1

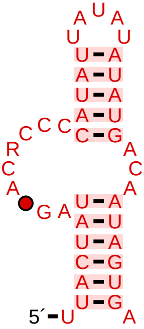

### SampleBMeta_195_1

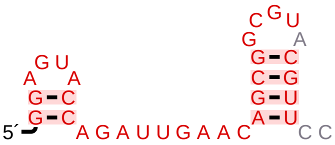

### SampleBMeta_199_1

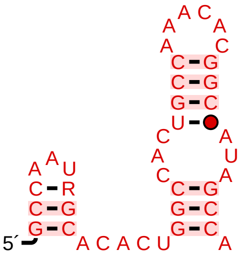

### SampleBMeta_20_1

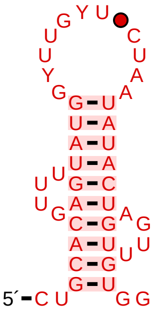

### SampleBMeta_200_1

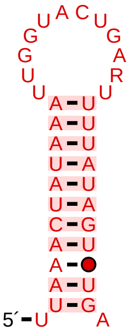

SampleBMeta_202_1

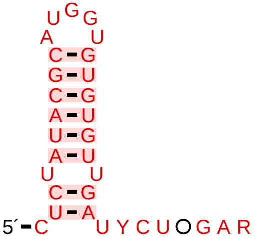

### SampleBMeta_203_1

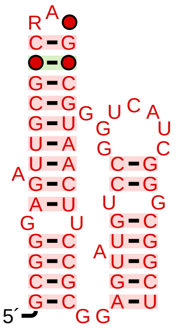

SampleBMeta_204_1

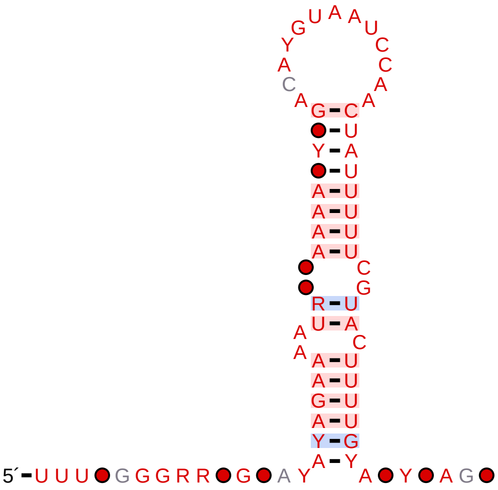

### SampleBMeta_205_1

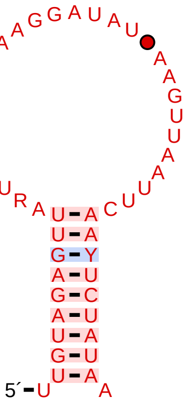

SampleBMeta_206_1

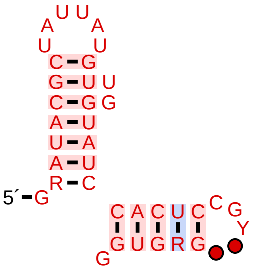

### SampleBMeta_207_1

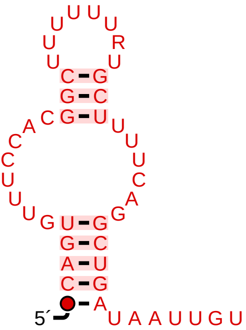

### SampleBMeta_215_1

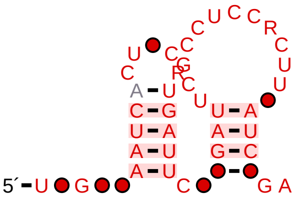

### SampleBMeta_216_1

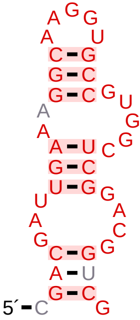

### SampleBMeta_217_1

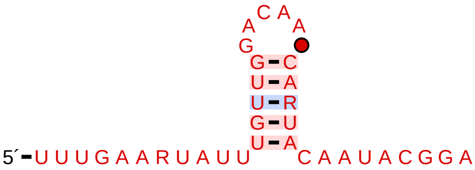

### SampleBMeta_221_1

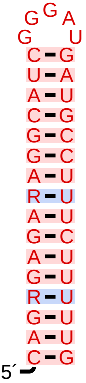

### SampleBMeta_222_1

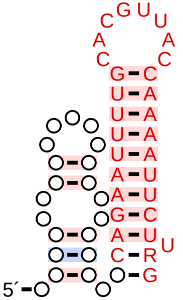

### SampleBMeta_226_1

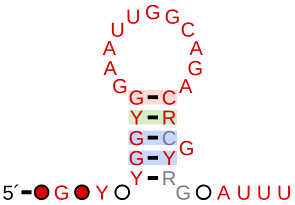

### SampleBMeta_228_1

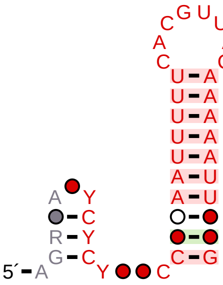

### SampleBMeta_233_1

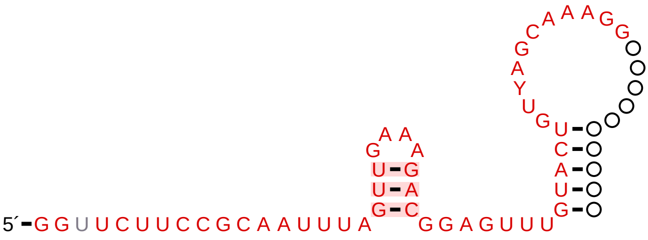

### SampleBMeta_235_1

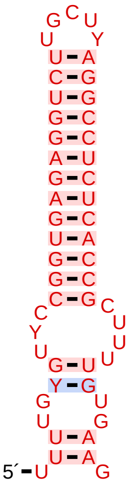

### SampleBMeta_237_1

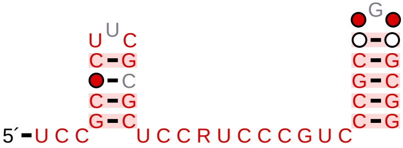

### SampleBMeta_238_1

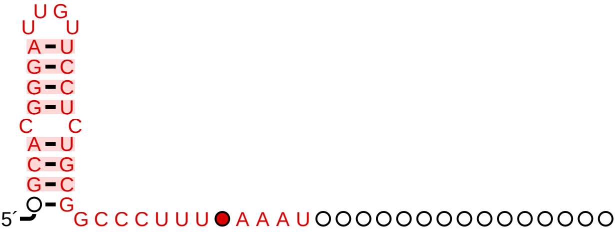

### SampleBMeta_239_1

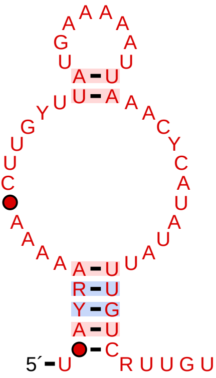

### SampleBMeta_241_1

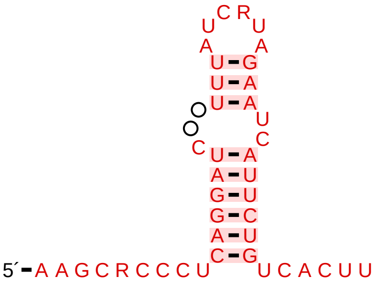

### SampleBMeta_242_1

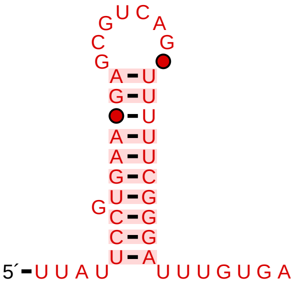

SampleBMeta_245_1

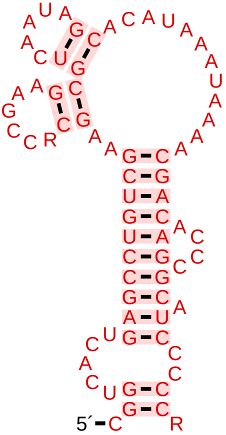

### SampleBMeta_251_1

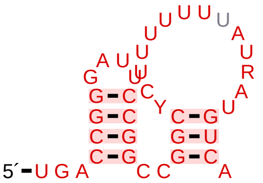

### SampleBMeta_254_1

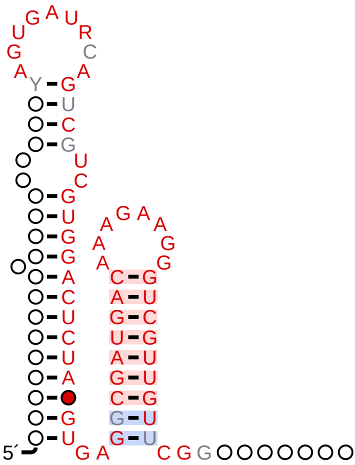

### SampleBMeta_258_1

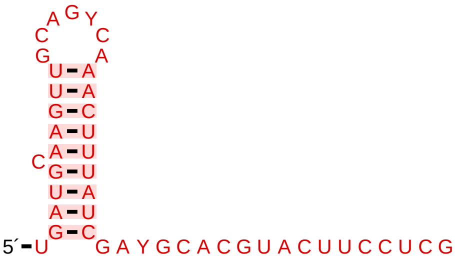

### SampleBMeta_259_1

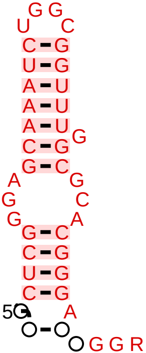

### SampleBMeta_26_1

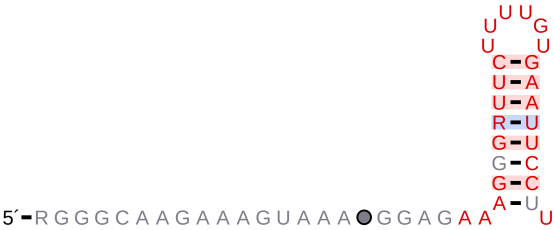

### SampleBMeta_260_1

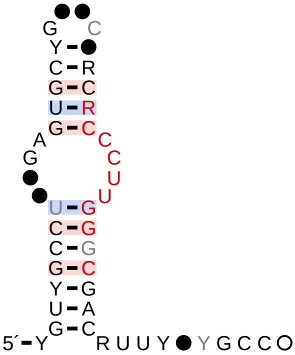

### SampleBMeta_261_1

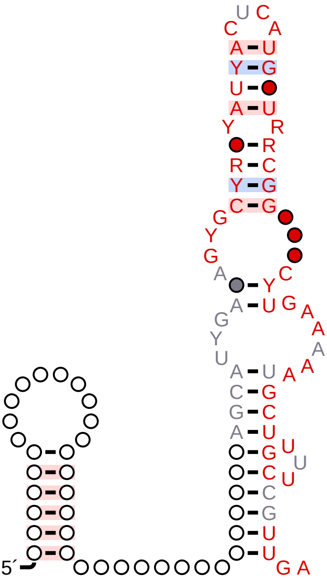

SampleBMeta_263_1

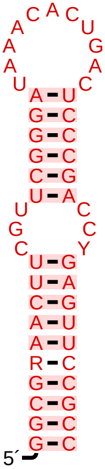

### SampleBMeta_264_1

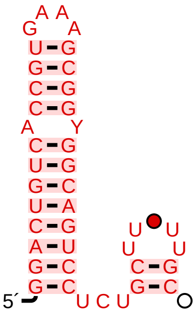

SampleBMeta_265_1

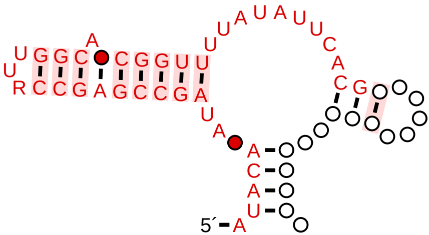

### SampleBMeta_27_1

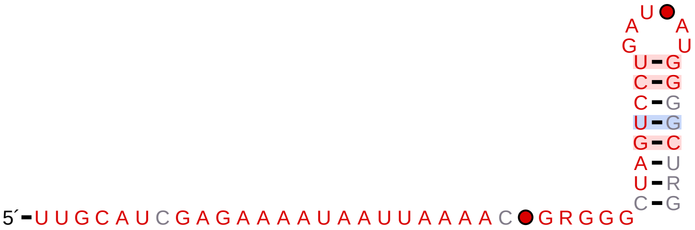

SampleBMeta_272_1

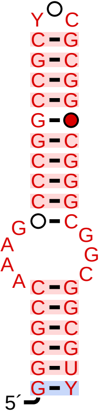

### SampleBMeta_275_1

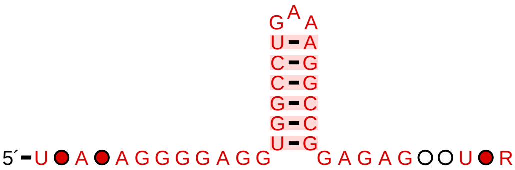

SampleBMeta_278_1

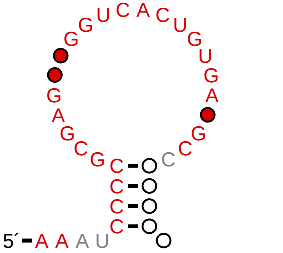

### SampleBMeta_279_1

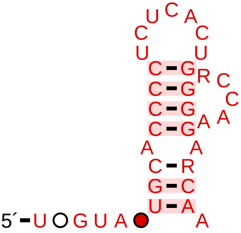

### SampleBMeta_28_1

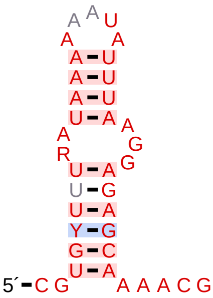

### SampleBMeta_280_1

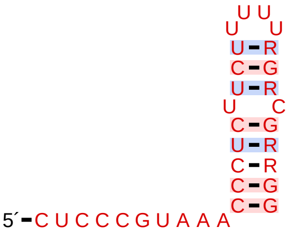

SampleBMeta_282_1

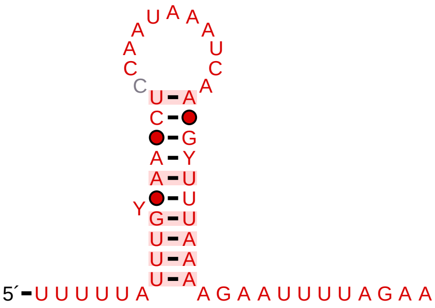

### SampleBMeta_283_1

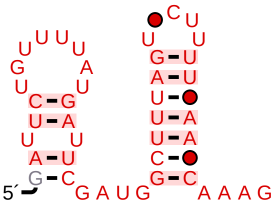

SampleBMeta_285_1

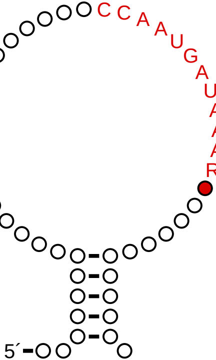

### SampleBMeta_286_1

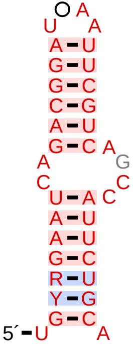

### SampleBMeta_291_1

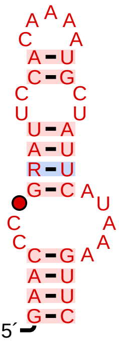

### SampleBMeta_293_1

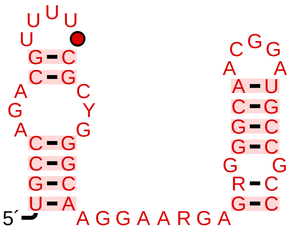

SampleBMeta_294_1

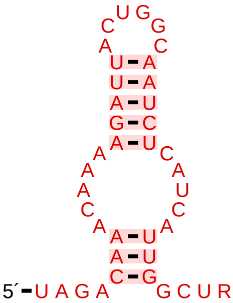

### SampleBMeta_296_1

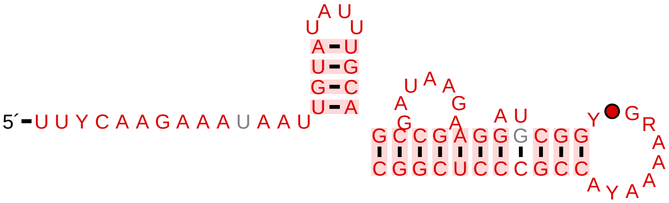

SampleBMeta_297_1

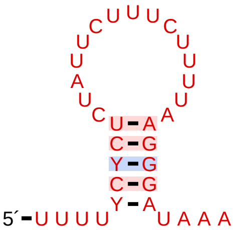

SampleBMeta_3_1

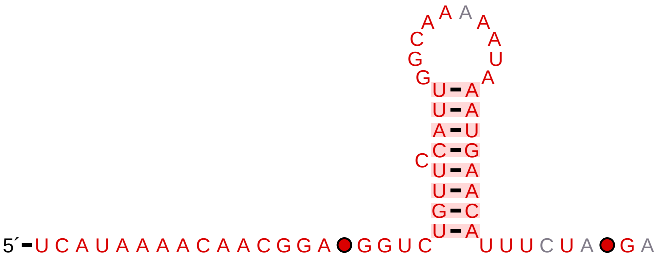

### SampleBMeta_305_1

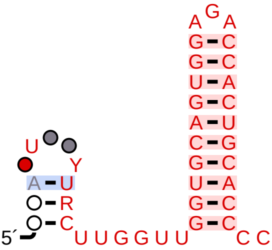

SampleBMeta_307_1

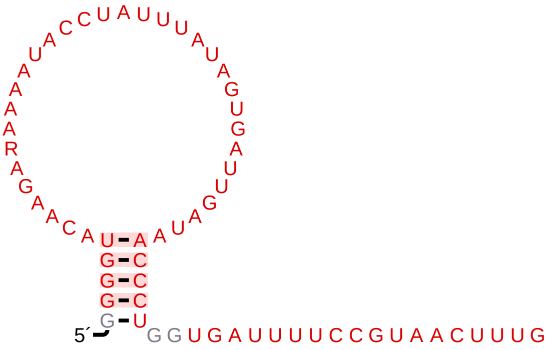

### SampleBMeta_314_1

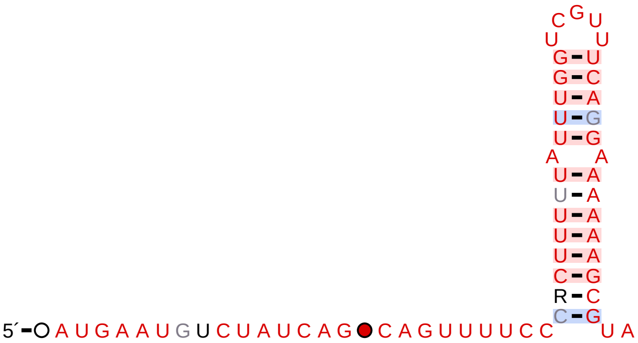

### SampleBMeta_315_1

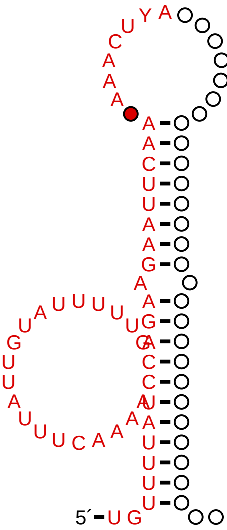

### SampleBMeta_316_1

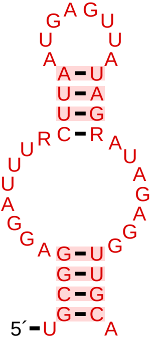

### SampleBMeta_317_1

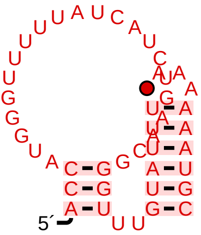

### SampleBMeta_320_1

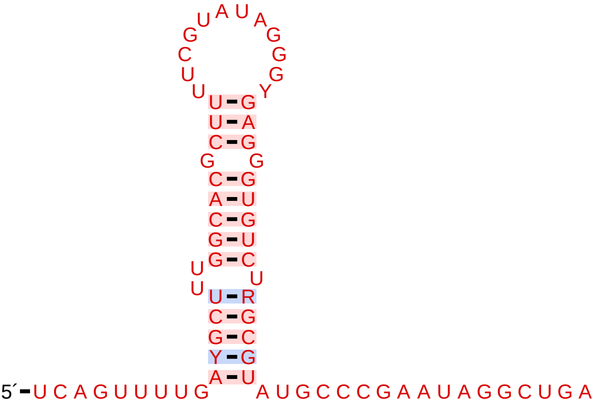

### SampleBMeta_323_1

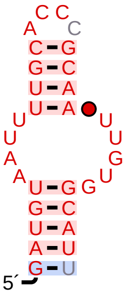

### SampleBMeta_327_1

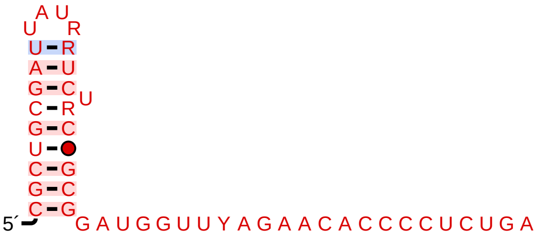

### SampleBMeta_334_1

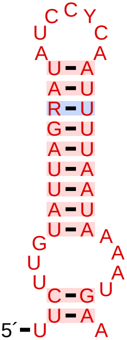

SampleBMeta_335_1

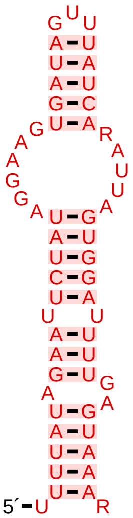

SampleBMeta_337_1

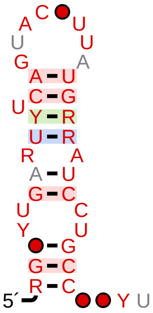

### SampleBMeta_341_1

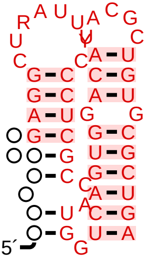

### SampleBMeta_344_1

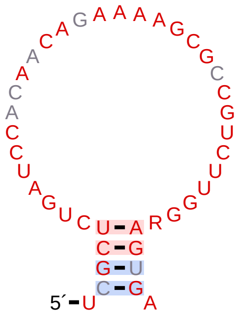

### SampleBMeta_35_1

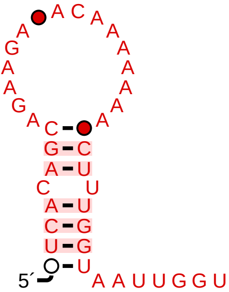

### SampleBMeta_351_1

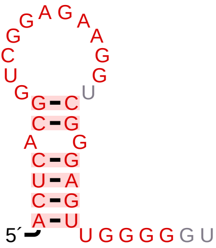

### SampleBMeta_356_1

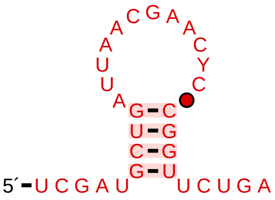

### SampleBMeta_362_1

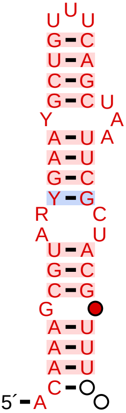

### SampleBMeta_366_1

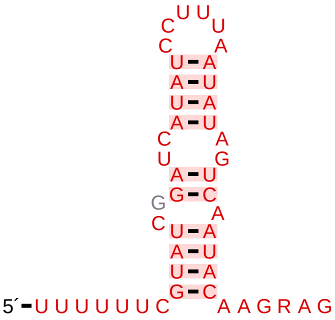

SampleBMeta_367_1

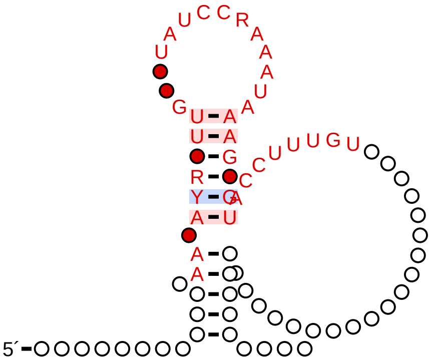

### SampleBMeta_370_1

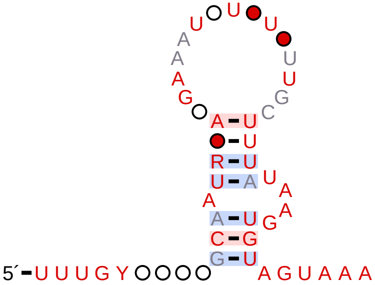

### SampleBMeta_371_1

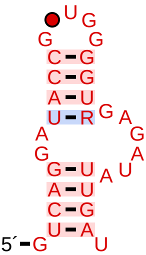

### SampleBMeta_373_1

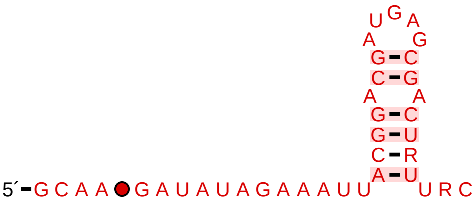

SampleBMeta_379_1

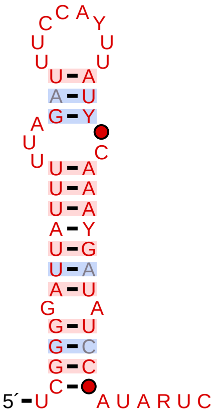

### SampleBMeta_38_1

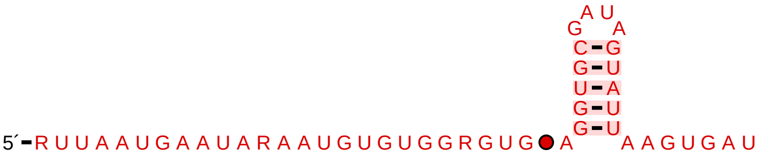

### SampleBMeta_382_1

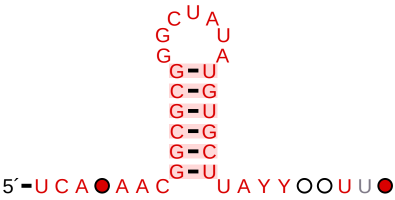

### SampleBMeta_384_1

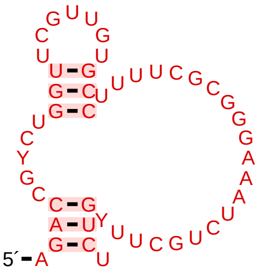

### SampleBMeta_388_1

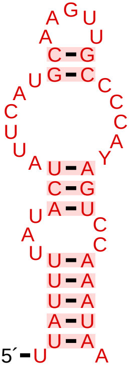

### SampleBMeta_389_1

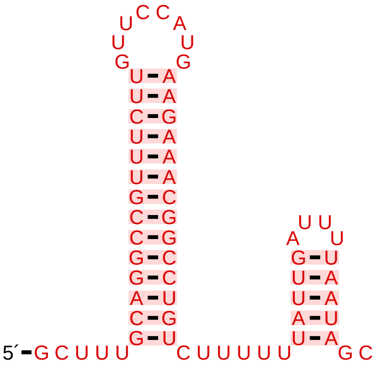

### SampleBMeta_39_1

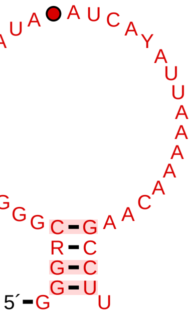

### SampleBMeta_394_1

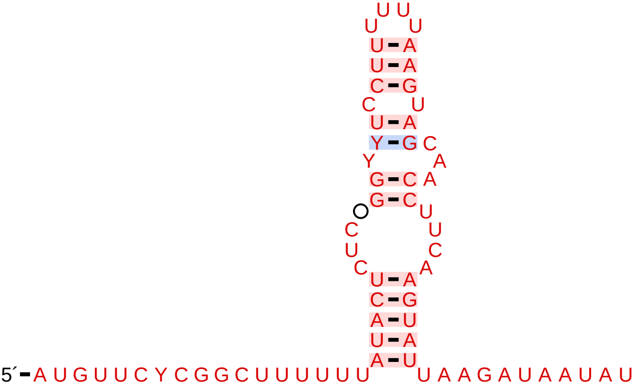

### SampleBMeta_395_1

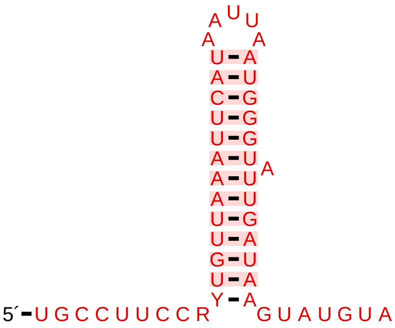

### SampleBMeta_399_1

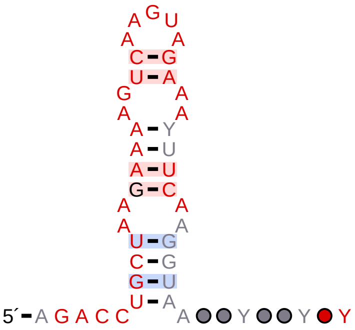

### SampleBMeta_4_1

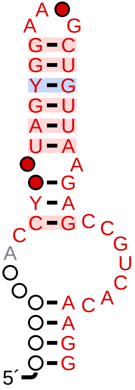

SampleBMeta_40_1

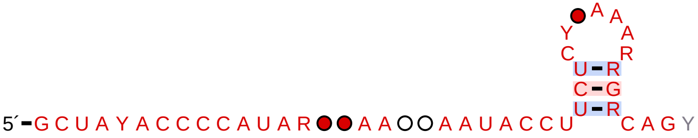

### SampleBMeta_400_1

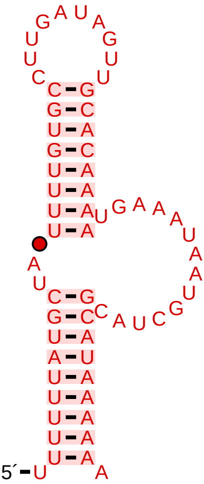

### SampleBMeta_408_1

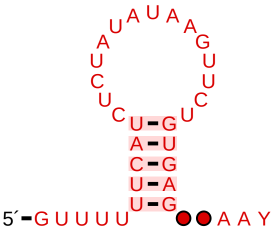

### SampleBMeta_41_1

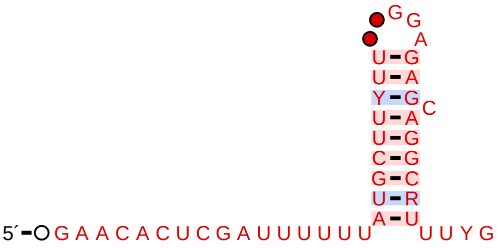

### SampleBMeta_411_1

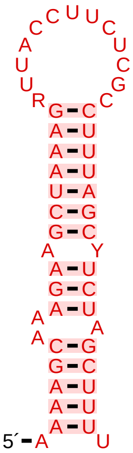

### SampleBMeta_414_1

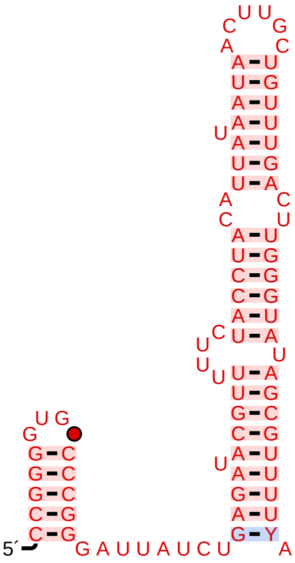

### SampleBMeta_416_1

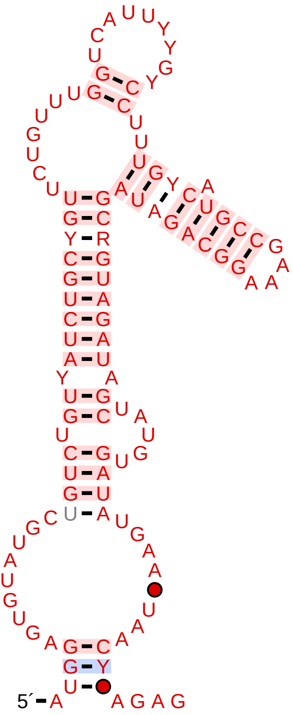

### SampleBMeta_418_1

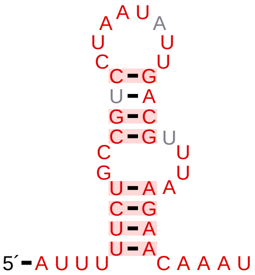

SampleBMeta_419_1

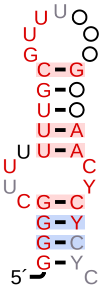

SampleBMeta_420_1

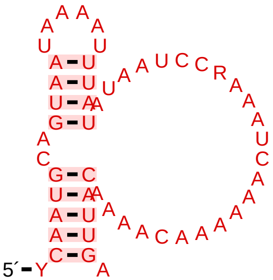

SampleBMeta_421_1

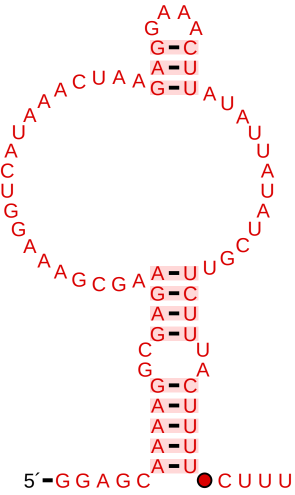

### SampleBMeta_423_1

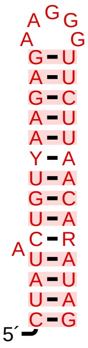

### SampleBMeta_426_1

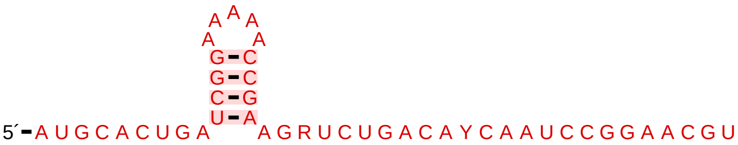

### SampleBMeta_427_1

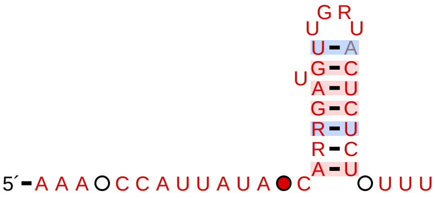

SampleBMeta_429_1

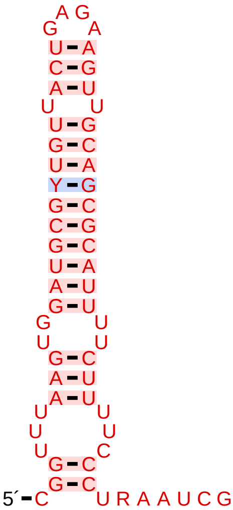

### SampleBMeta_430_1

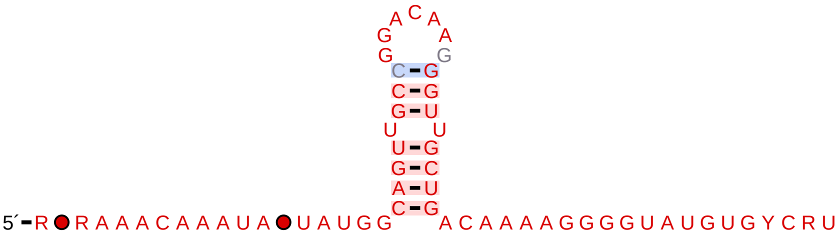

SampleBMeta_431_1

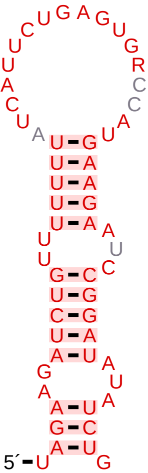

### SampleBMeta_432_1

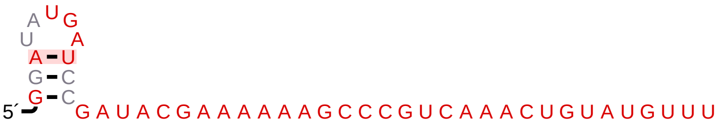

### SampleBMeta_433_1

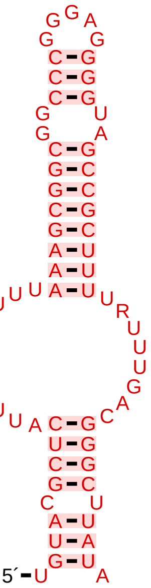

### SampleBMeta_435_1

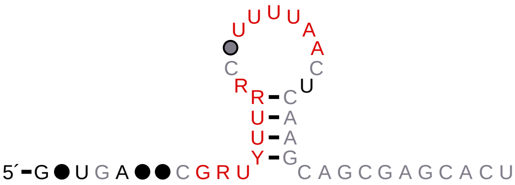

### SampleBMeta_44_1

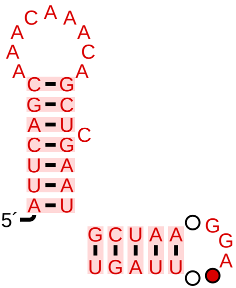

### SampleBMeta_442_1

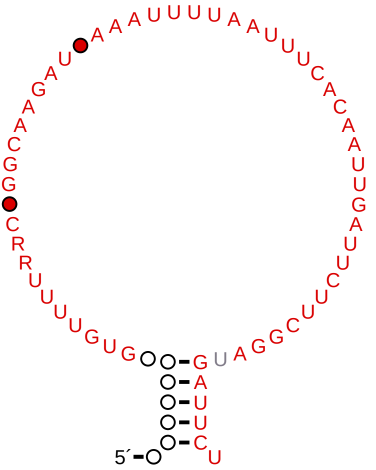

### SampleBMeta_444_1

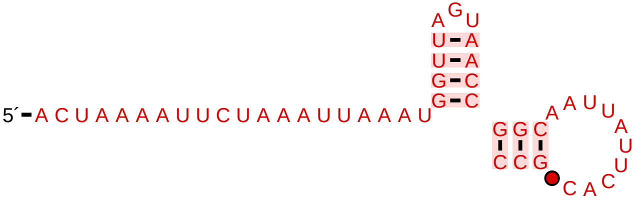

### SampleBMeta_446_1

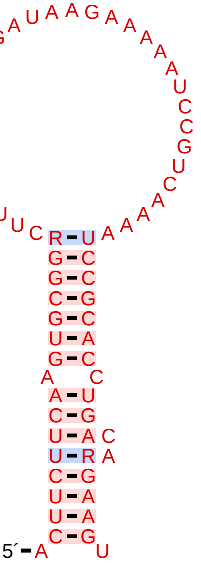

### SampleBMeta_447_1

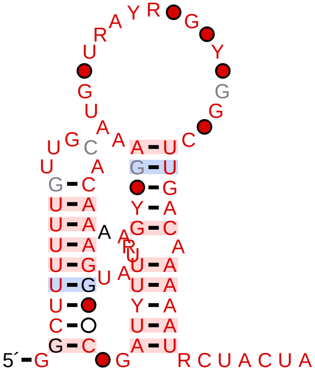

SampleBMeta_450_1

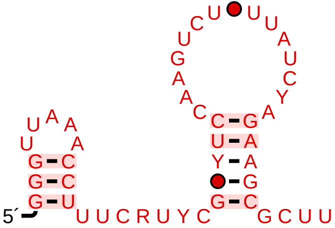

### SampleBMeta_452_1

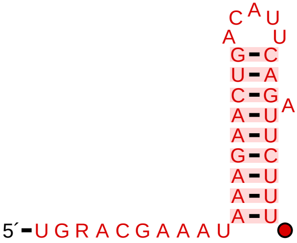

### SampleBMeta_453_1

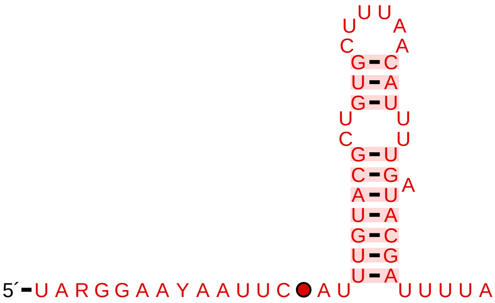

### SampleBMeta_46_1

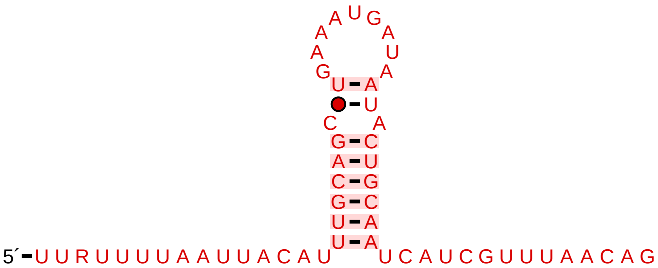

### SampleBMeta_461_1

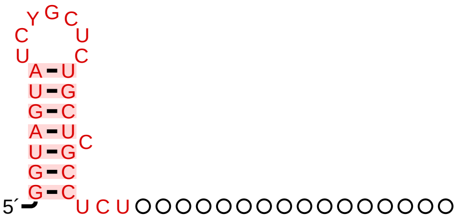

SampleBMeta_462_1

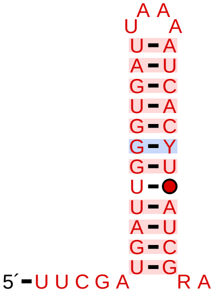

SampleBMeta_464_1

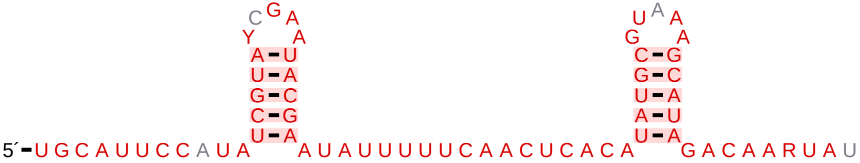

### SampleBMeta_468_1

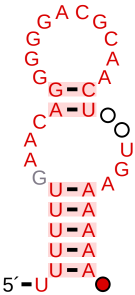

### SampleBMeta_470_1

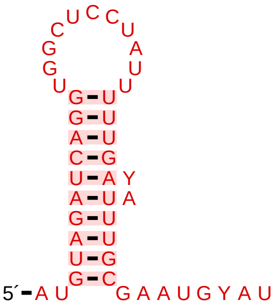

### SampleBMeta_472_1

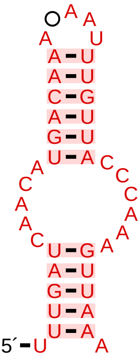

SampleBMeta_477_1

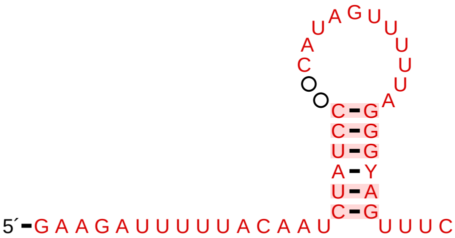

### SampleBMeta_479_1

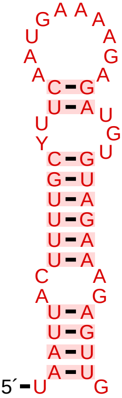

### SampleBMeta_480_1

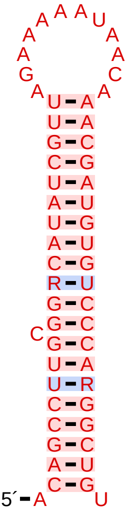

### SampleBMeta_482_1

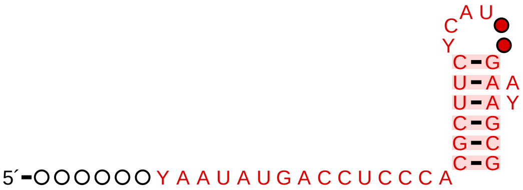

### SampleBMeta_485_1

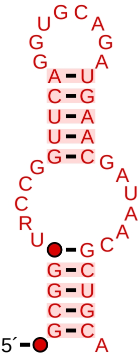

### SampleBMeta_487_1

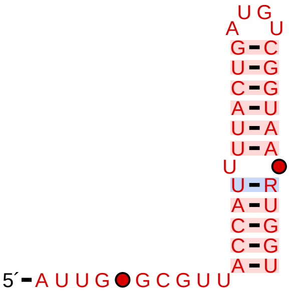

### SampleBMeta_488_1

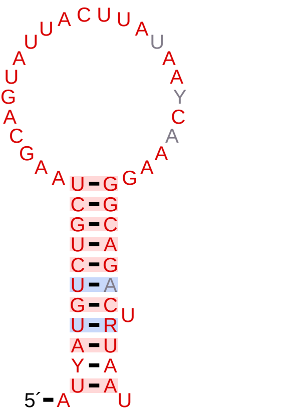

### SampleBMeta_490_1

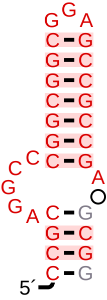

SampleBMeta_492_1

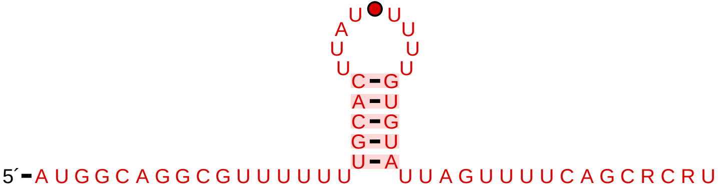

### SampleBMeta_495_1

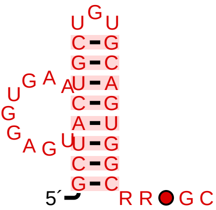

### SampleBMeta_497_1

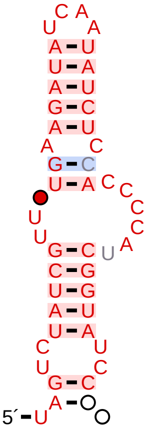

### SampleBMeta_503_1

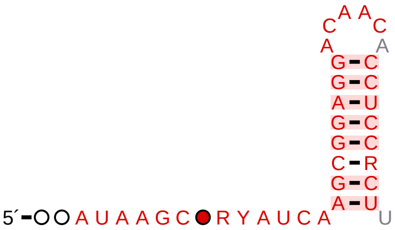

### SampleBMeta_504_1

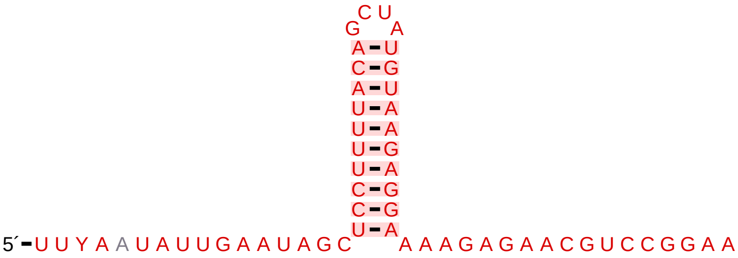

### SampleBMeta_505_1

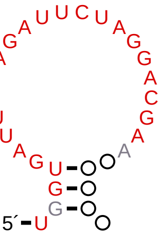

SampleBMeta_506_1

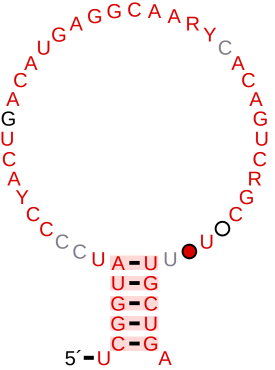

### SampleBMeta_508_1

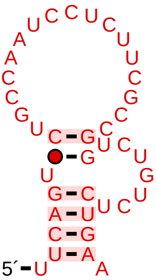

### SampleBMeta_51_1

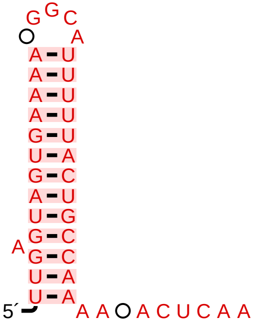

### SampleBMeta_510_1

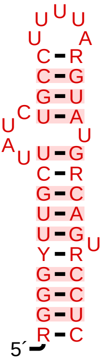

SampleBMeta_516_1

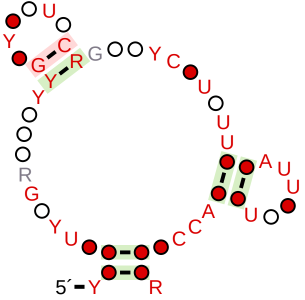

### SampleBMeta_518_1

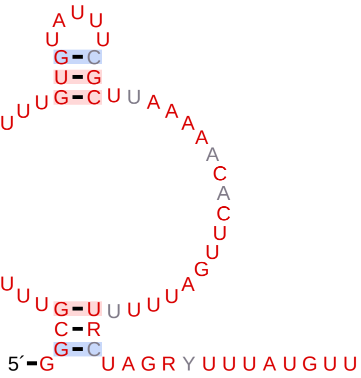

### SampleBMeta_521_1

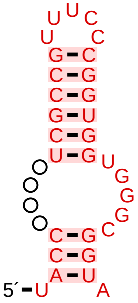

### SampleBMeta_523_1

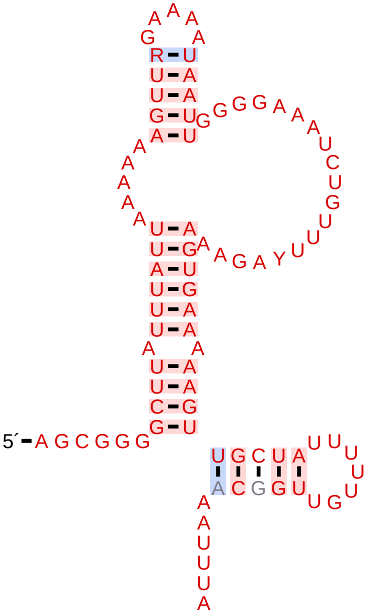

SampleBMeta_524_1

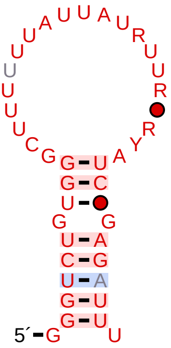

### SampleBMeta_526_1

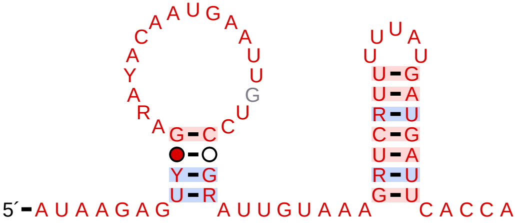

SampleBMeta_528_1

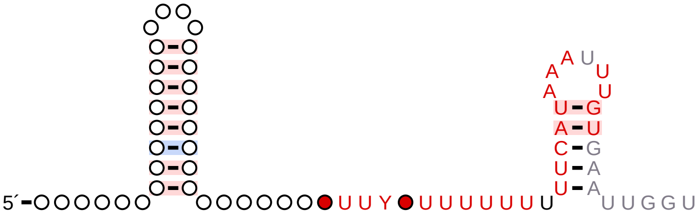

### SampleBMeta_531_1

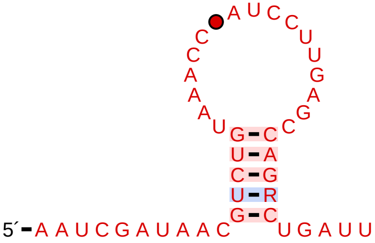

### SampleBMeta_537_1

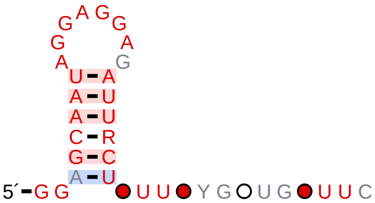

### SampleBMeta_538_1

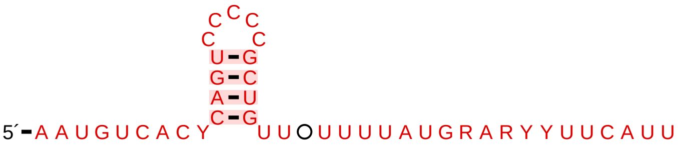

### SampleBMeta_539_1

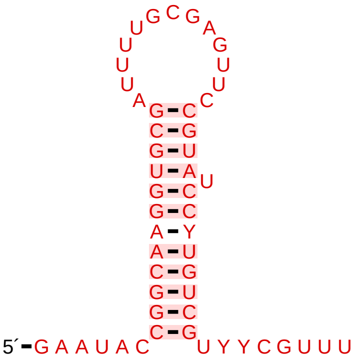

SampleBMeta_54_1

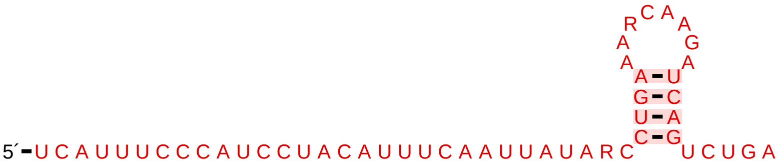

### SampleBMeta_541_1

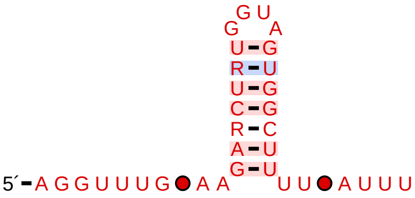

### SampleBMeta_544_1

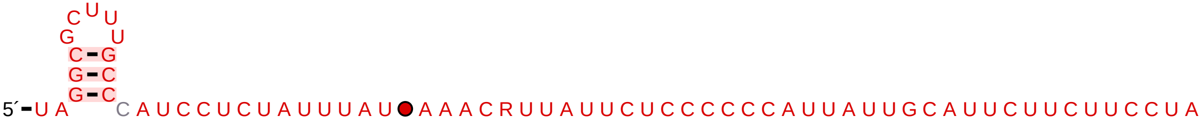

### SampleBMeta_547_1

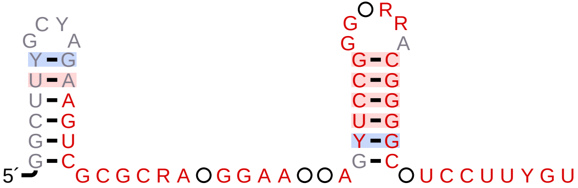

### SampleBMeta_548_1

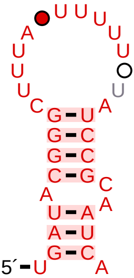

### SampleBMeta_552_1

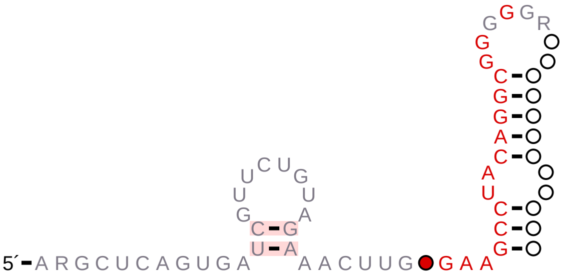

### SampleBMeta_560_1

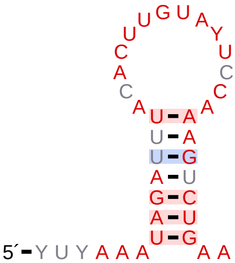

### SampleBMeta_562_1

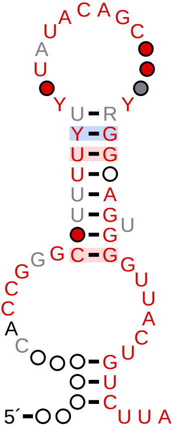

### SampleBMeta_564_1

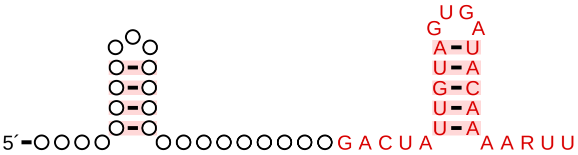

### SampleBMeta_566_1

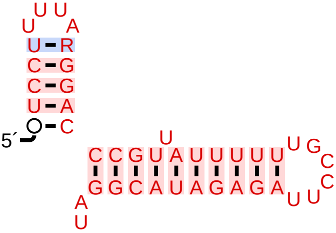

### SampleBMeta_569_1

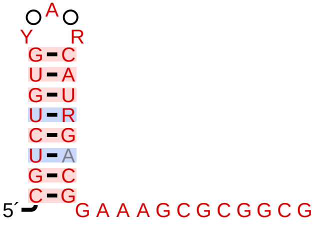

SampleBMeta_575_1

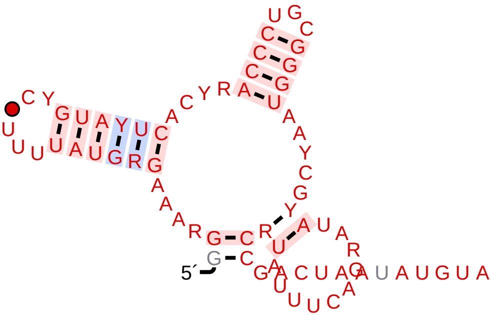

### SampleBMeta_577_1

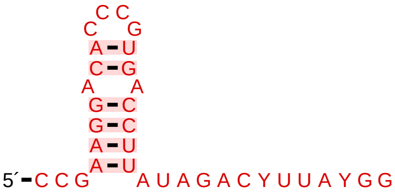

### SampleBMeta_578_1

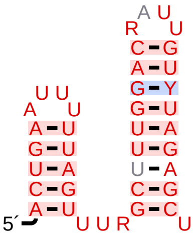

### SampleBMeta_580_1

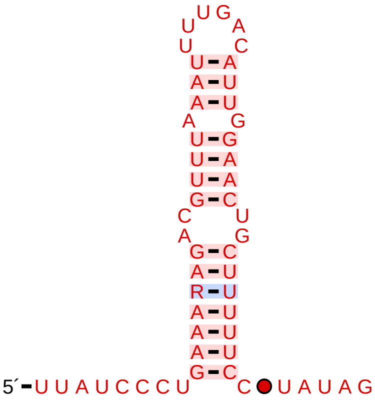

SampleBMeta_581_1

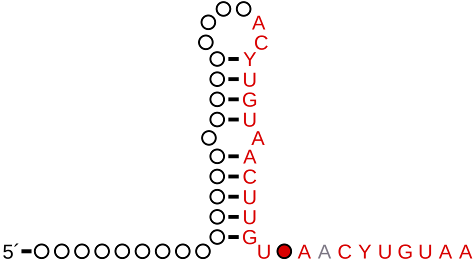

SampleBMeta_582_1

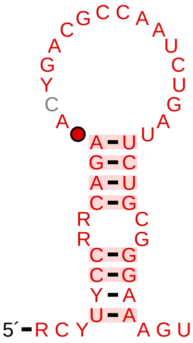

### SampleBMeta_586_1

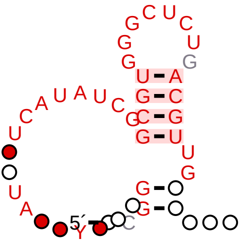

### SampleBMeta_587_1

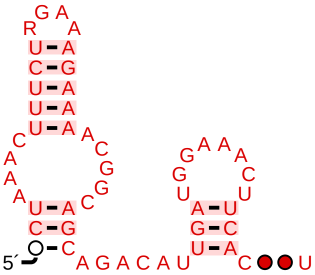

### SampleBMeta_590_1

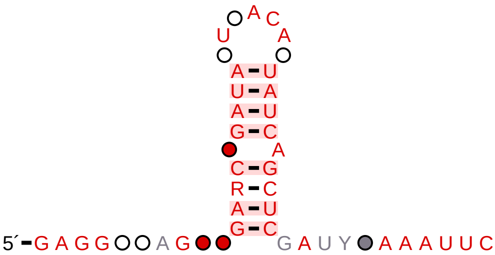

SampleBMeta_591_1

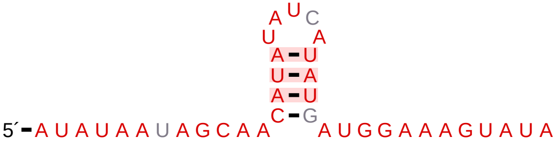

### SampleBMeta_592_1

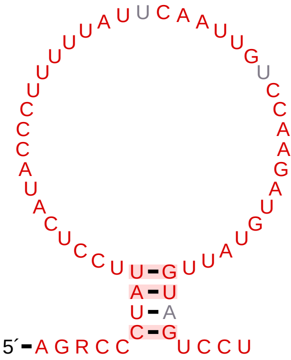

### SampleBMeta_594_1

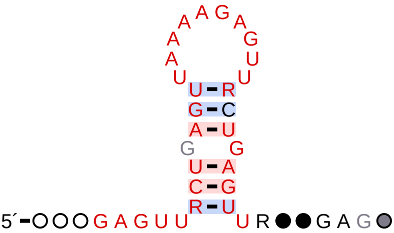

### SampleBMeta_596_1

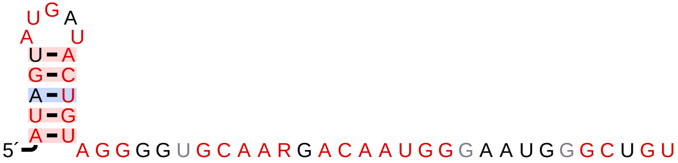

SampleBMeta_597_1

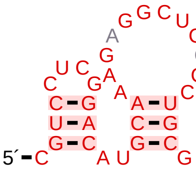

SampleBMeta_60_1

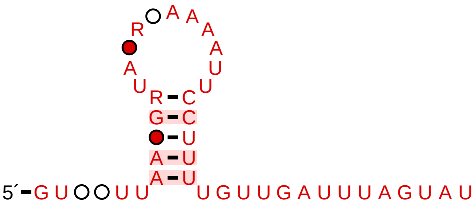

### SampleBMeta_605_1

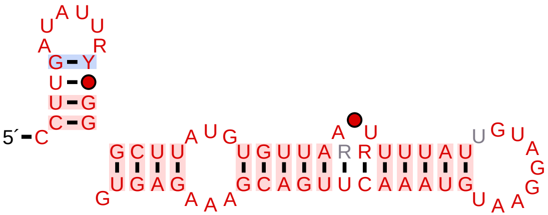

### SampleBMeta_608_1

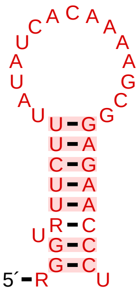

SampleBMeta_61_1

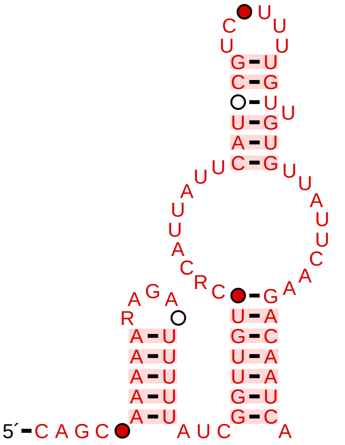

### SampleBMeta_610_1

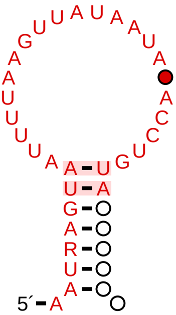

### SampleBMeta_614_1

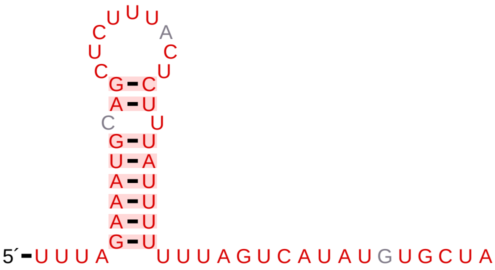

### SampleBMeta_615_1

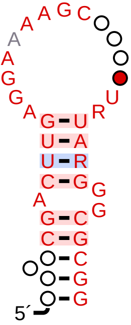

### SampleBMeta_616_1

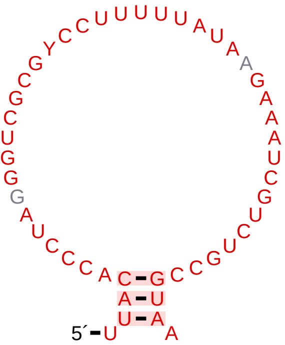

SampleBMeta_618_1

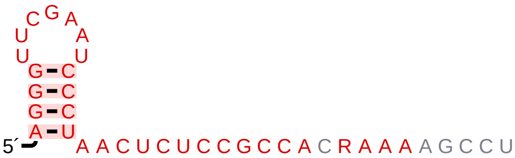

### SampleBMeta_621_1

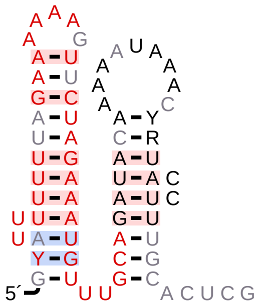

SampleBMeta_622_1

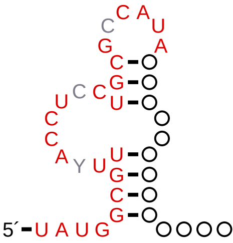

SampleBMeta_623_1

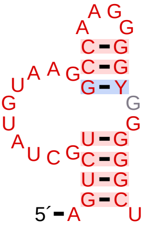

### SampleBMeta_626_1

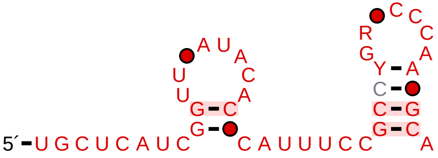

### SampleBMeta_628_1

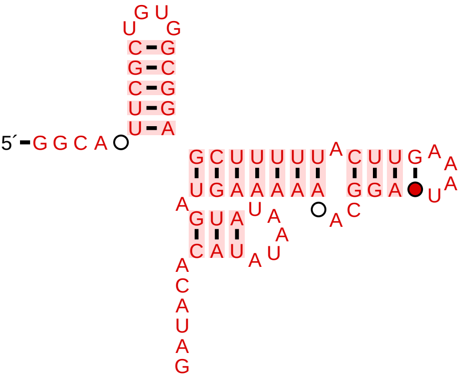

### SampleBMeta_63_1

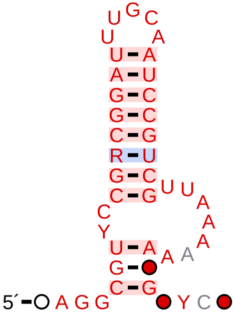

### SampleBMeta_631_1

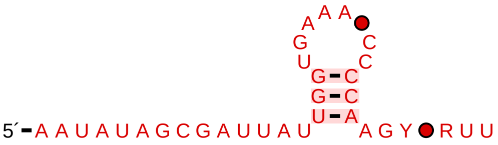

SampleBMeta_632_1

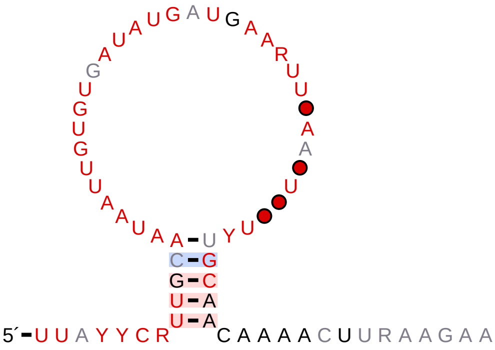

### SampleBMeta_633_1

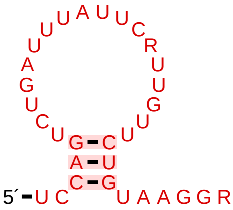

SampleBMeta_634_1

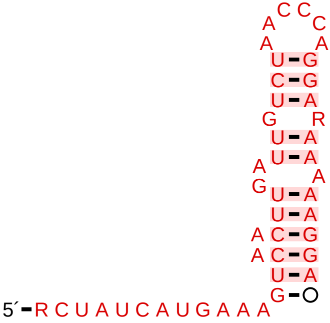

### SampleBMeta_640_1

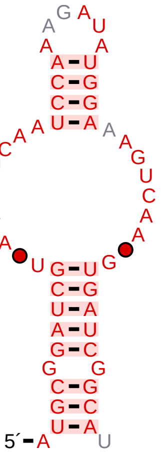

### SampleBMeta_642_1

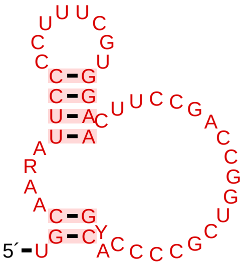

### SampleBMeta_65_1

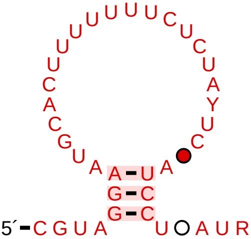

### SampleBMeta_656_1

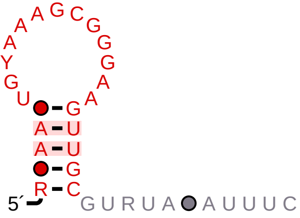

### SampleBMeta_660_1

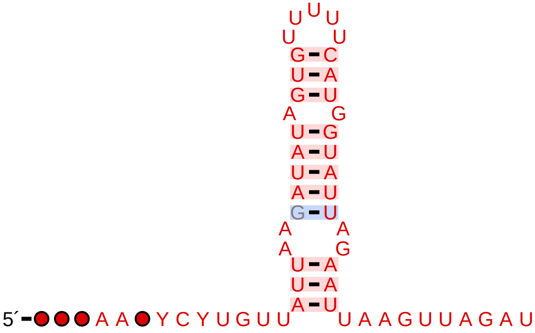

### SampleBMeta_662_1

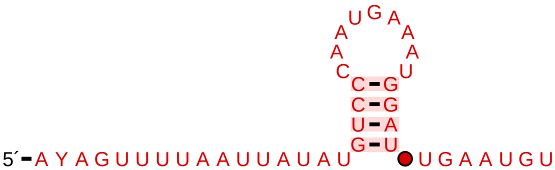

### SampleBMeta_665_1

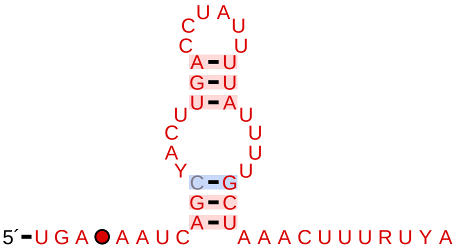

### SampleBMeta_666_1

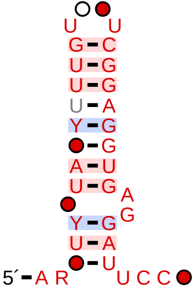

### SampleBMeta_667_1

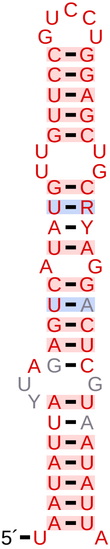

### SampleBMeta_669_1

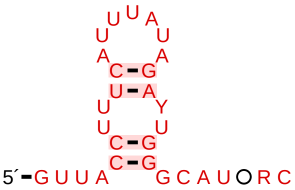

SampleBMeta_674_1

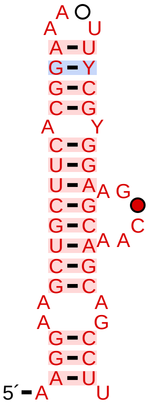

### SampleBMeta_675_1

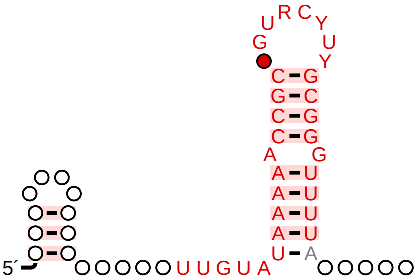

SampleBMeta_677_1

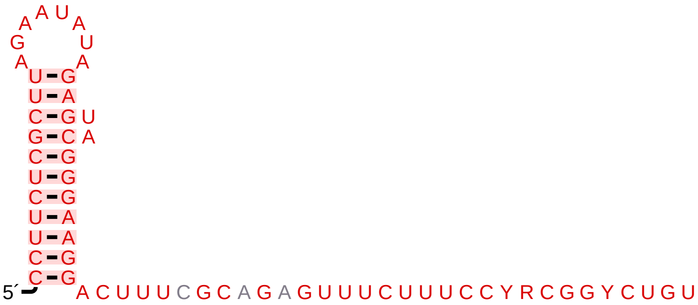

### SampleBMeta_678_1

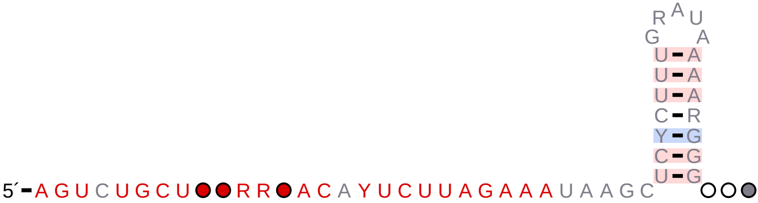

### SampleBMeta_679_1

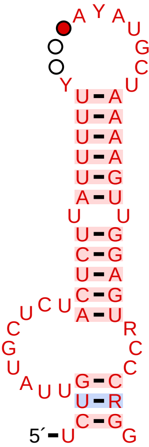

SampleBMeta_68_1

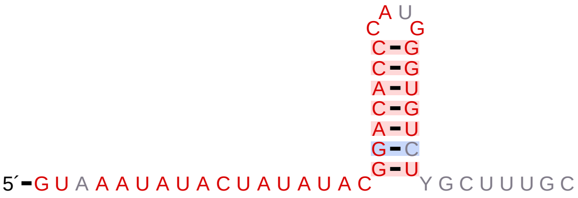

### SampleBMeta_680_1

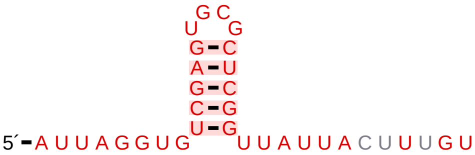

### SampleBMeta_681_1

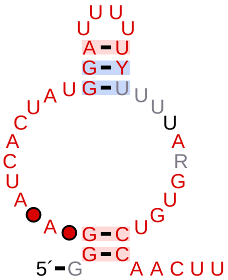

### SampleBMeta_69_1

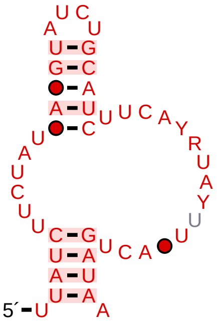

### SampleBMeta_690_1

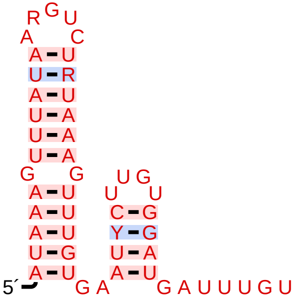

### SampleBMeta_692_1

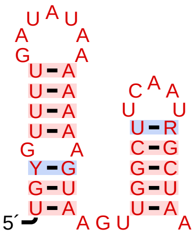

### SampleBMeta_693_1

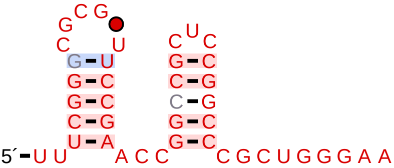

SampleBMeta_695_1

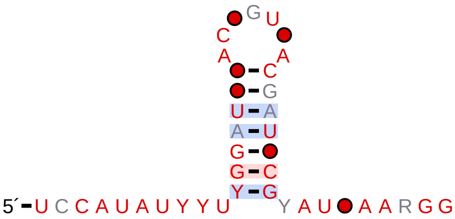

SampleBMeta_696_1

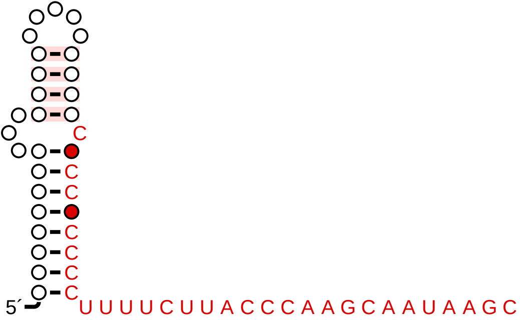

### SampleBMeta_697_1

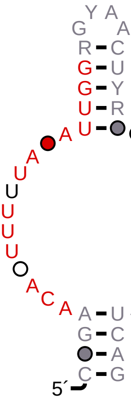

### SampleBMeta_698_1

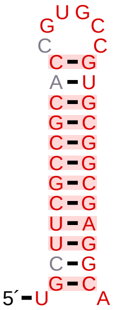

SampleBMeta_699_1

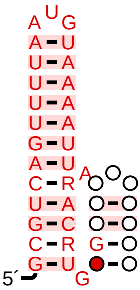

### SampleBMeta_7_1

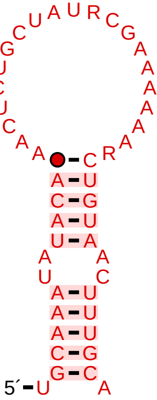

### SampleBMeta_700_1

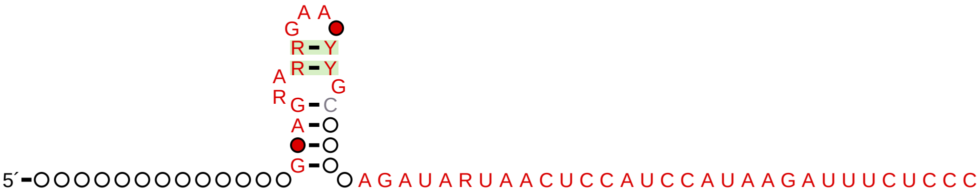

### SampleBMeta_701_1

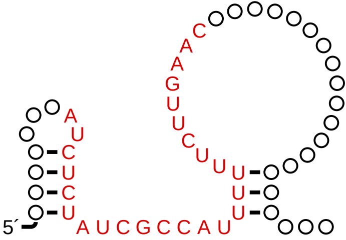

### SampleBMeta_703_1

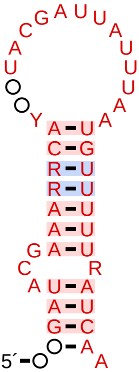

### SampleBMeta_705_1

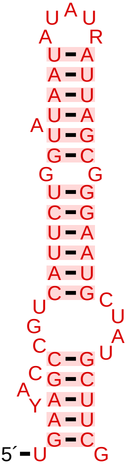

SampleBMeta_706_1

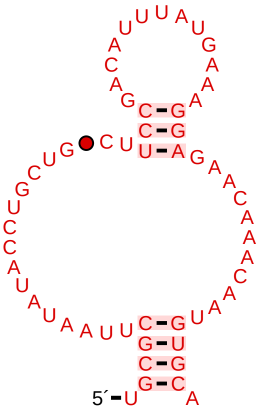

### SampleBMeta_711_1

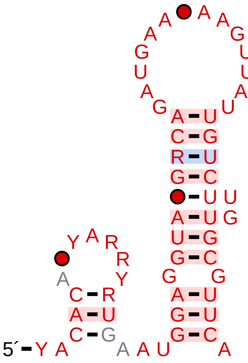

SampleBMeta_714_1

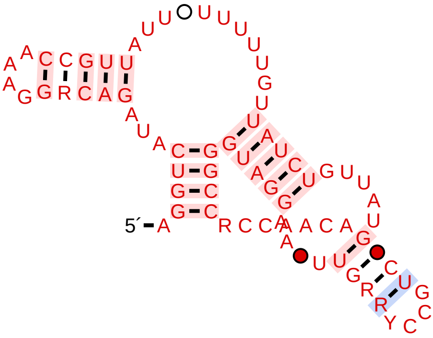

### SampleBMeta_717_1

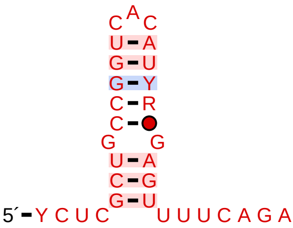

### SampleBMeta_724_1

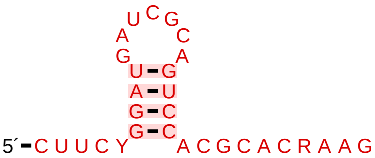

### SampleBMeta_73_1

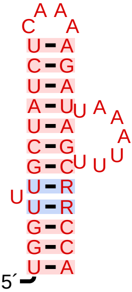

### SampleBMeta_732_1

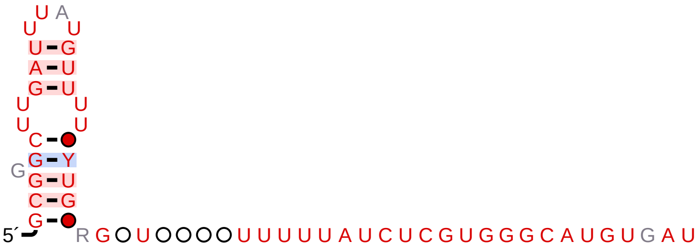

SampleBMeta_734_1

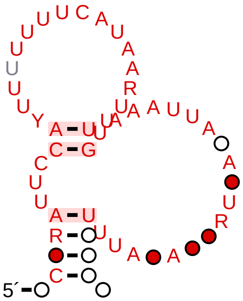

### SampleBMeta_736_1

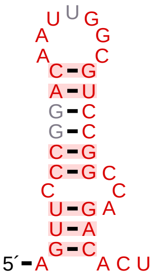

### SampleBMeta_738_1

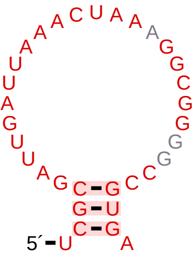

### SampleBMeta_74_1

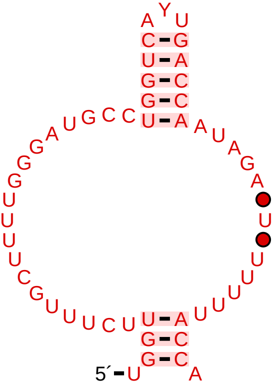

### SampleBMeta_740_1

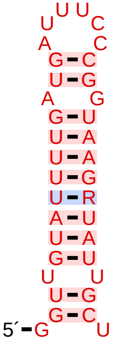

SampleBMeta_741_1

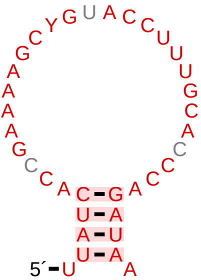

### SampleBMeta_742_1

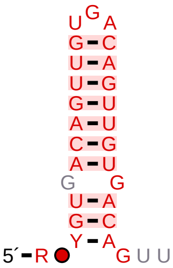

### SampleBMeta_743_1

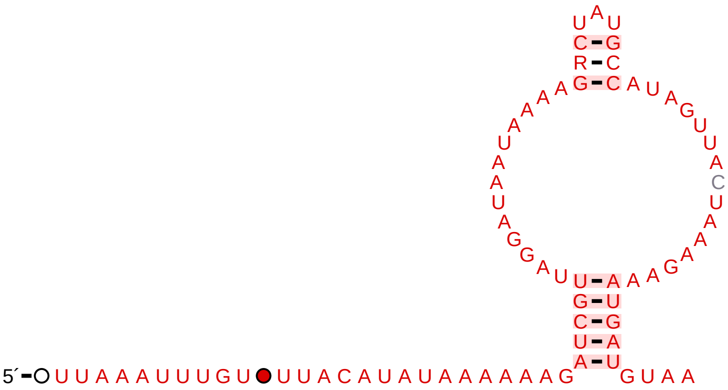

### SampleBMeta_75_1

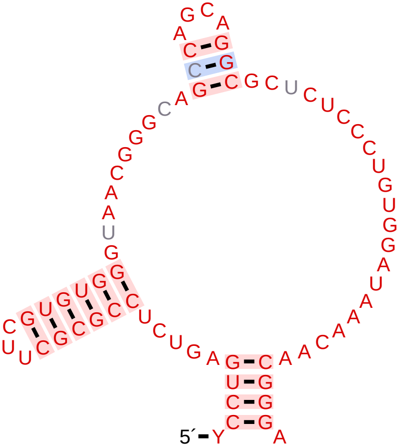

SampleBMeta_754_1

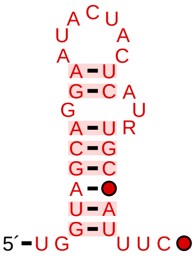

### SampleBMeta_756_1

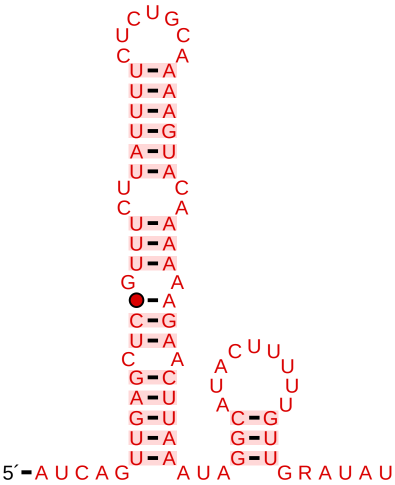

### SampleBMeta_757_1

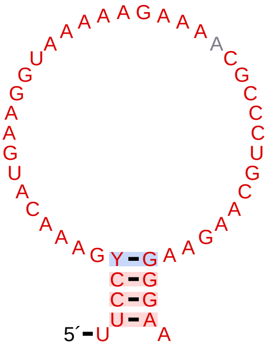

### SampleBMeta_76_1

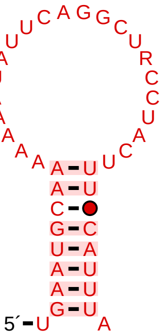

### SampleBMeta_761_1

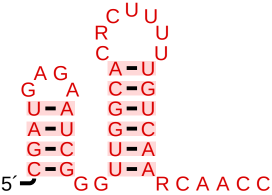

SampleBMeta_764_1

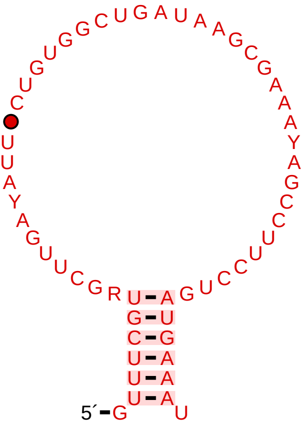

### SampleBMeta_765_1

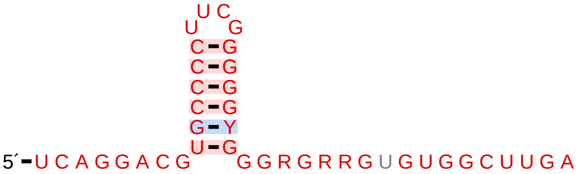

### SampleBMeta_766_1

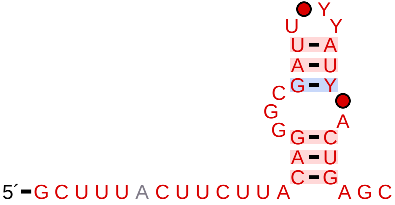

### SampleBMeta_769_1

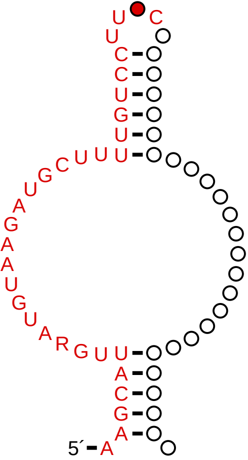

SampleBMeta_77_1

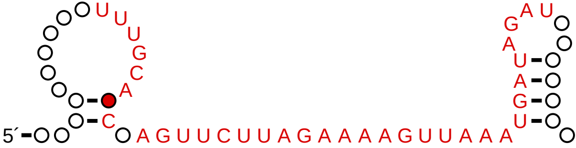

SampleBMeta_770_1

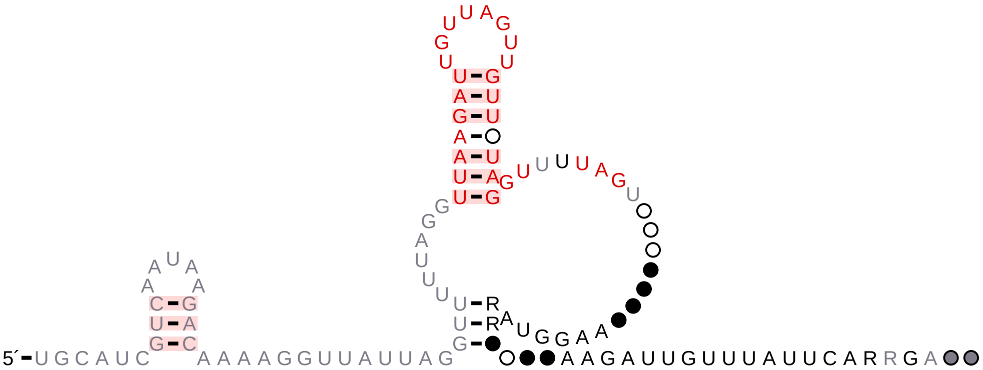

SampleBMeta_771_1

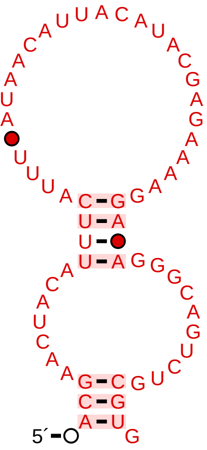

### SampleBMeta_772_1

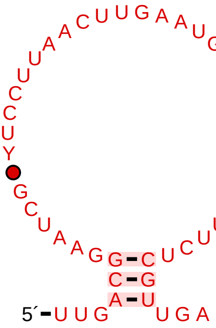

### SampleBMeta_777_1

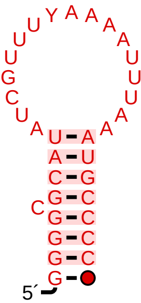

### SampleBMeta_782_1

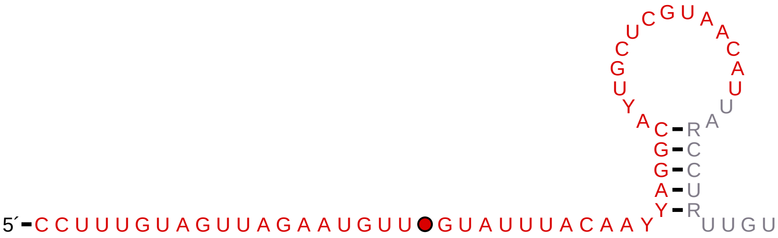

### SampleBMeta_784_1

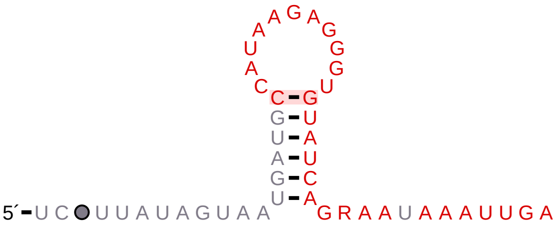

### SampleBMeta_785_1

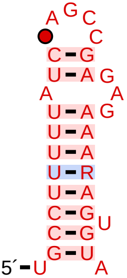

### SampleBMeta_786_1

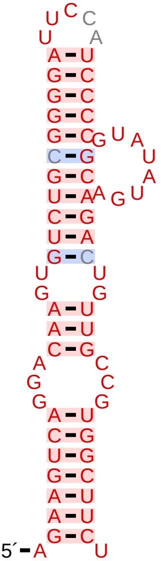

### SampleBMeta_788_1

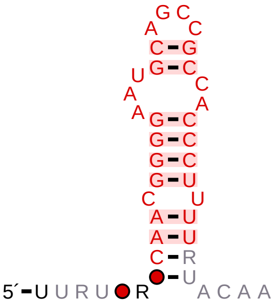

### SampleBMeta_791_1

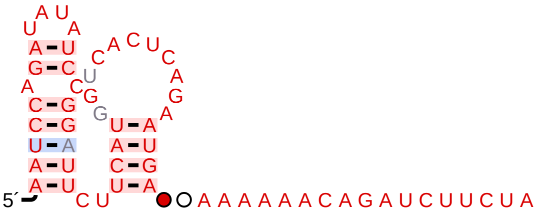

### SampleBMeta_792_1

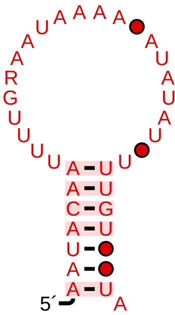

### SampleBMeta_795_1

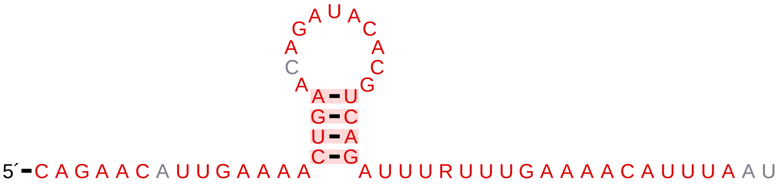

SampleBMeta_797_1

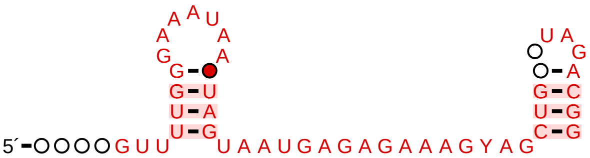

### SampleBMeta_80_1

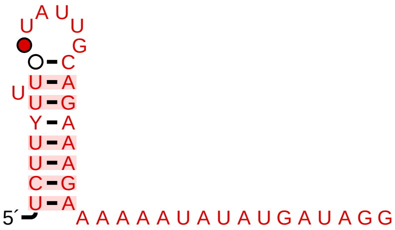

### SampleBMeta_800_1

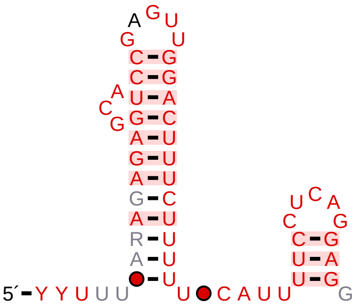

### SampleBMeta_803_1

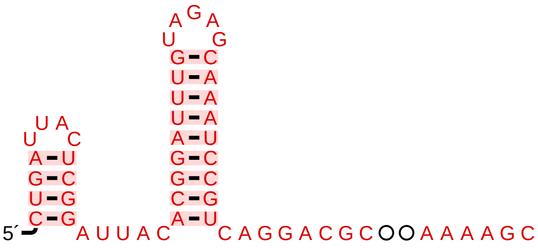

### SampleBMeta_804_1

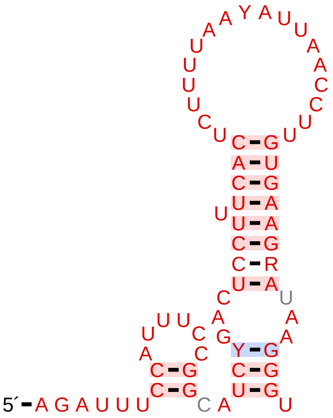

### SampleBMeta_805_1

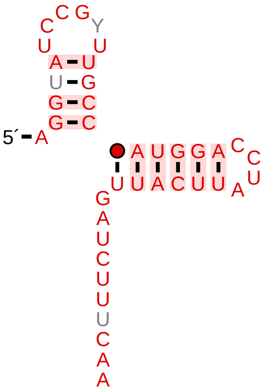

### SampleBMeta_810_1

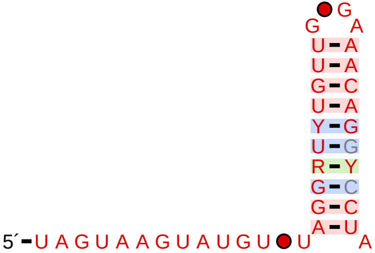

### SampleBMeta_813_1

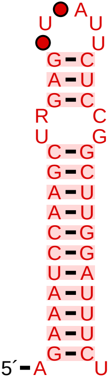

### SampleBMeta_814_1

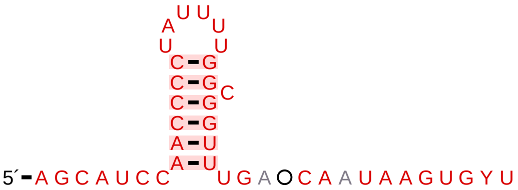

### SampleBMeta_815_1

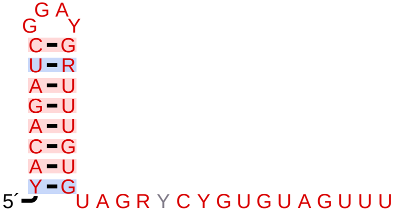

### SampleBMeta_819_1

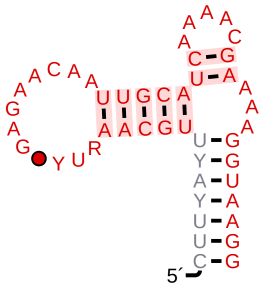

### SampleBMeta_82_1

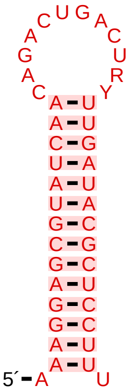

### SampleBMeta_83_1

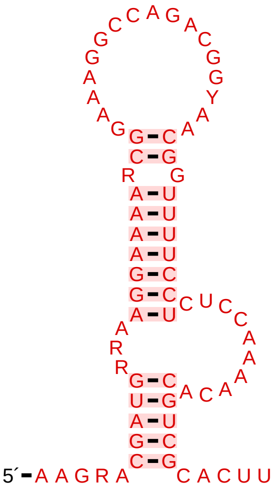

### SampleBMeta_87_1

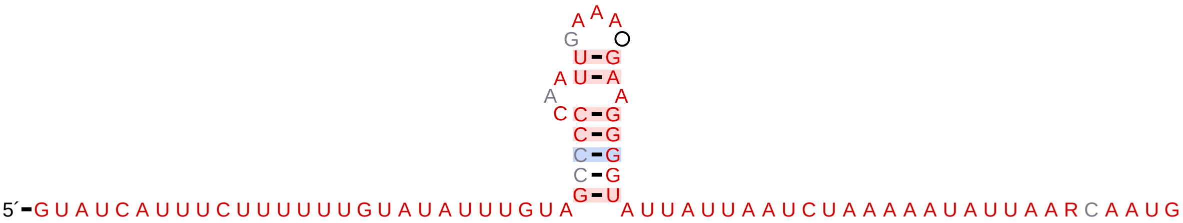

SampleBMeta_90_1

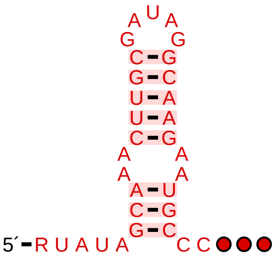

SampleBMeta_91_1

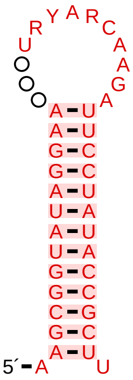

SampleBMeta_97_1

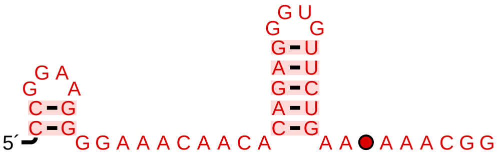

### SampleCMeta_10_1

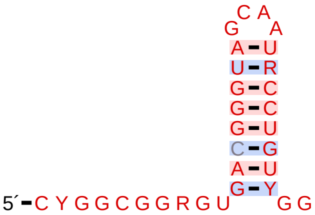

### SampleCMeta_100_1

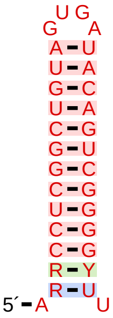

### SampleCMeta_103_1

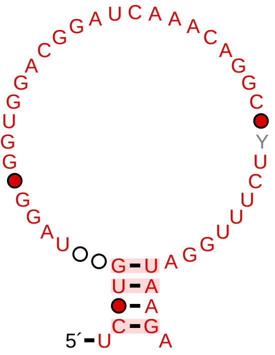

### SampleCMeta_107_1

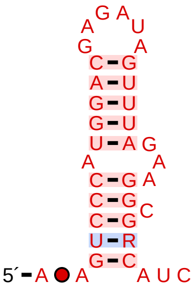

### SampleCMeta_11_1

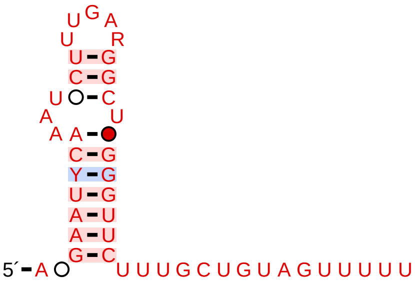

### SampleCMeta_113_1

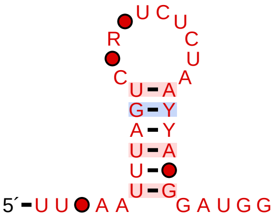

SampleCMeta_117_1

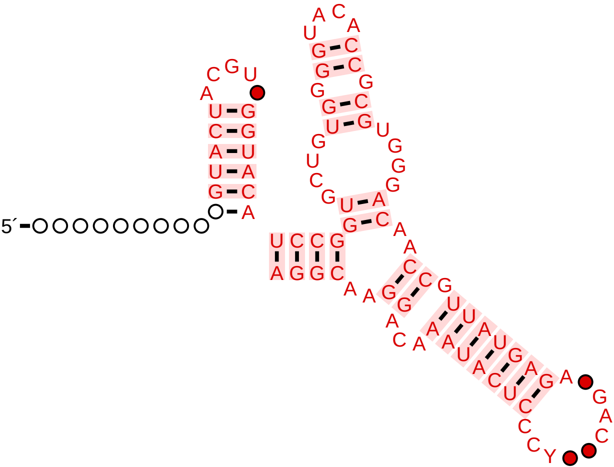

### SampleCMeta_118_1

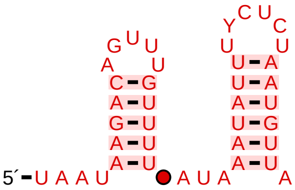

### SampleCMeta_12_1

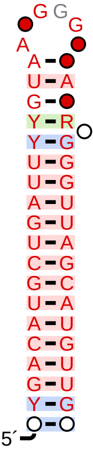

### SampleCMeta_120_1

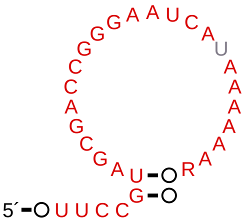

### SampleCMeta_126_1

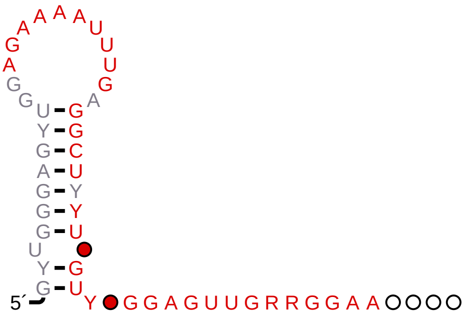

SampleCMeta_127_1

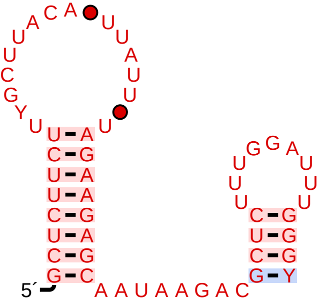

SampleCMeta_129_1

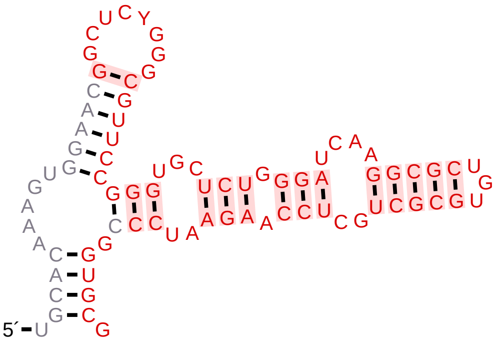

### SampleCMeta_131_1

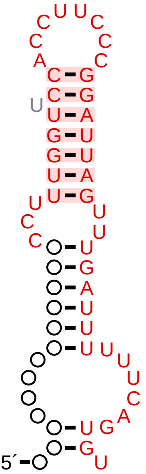

### SampleCMeta_132_1

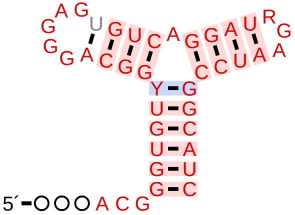

### SampleCMeta_133_1

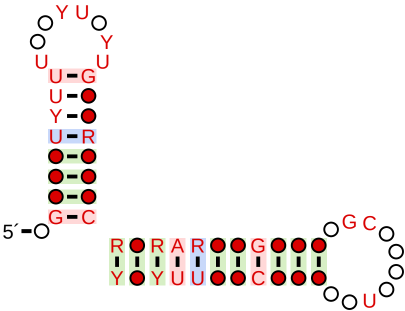

SampleCMeta_140_1

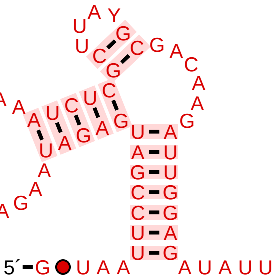

### SampleCMeta_142_1

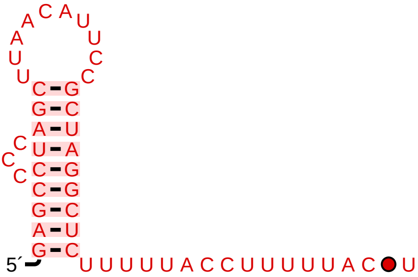

### SampleCMeta_143_1

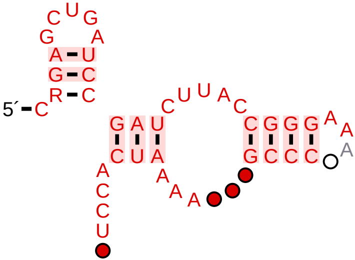

SampleCMeta_148_1

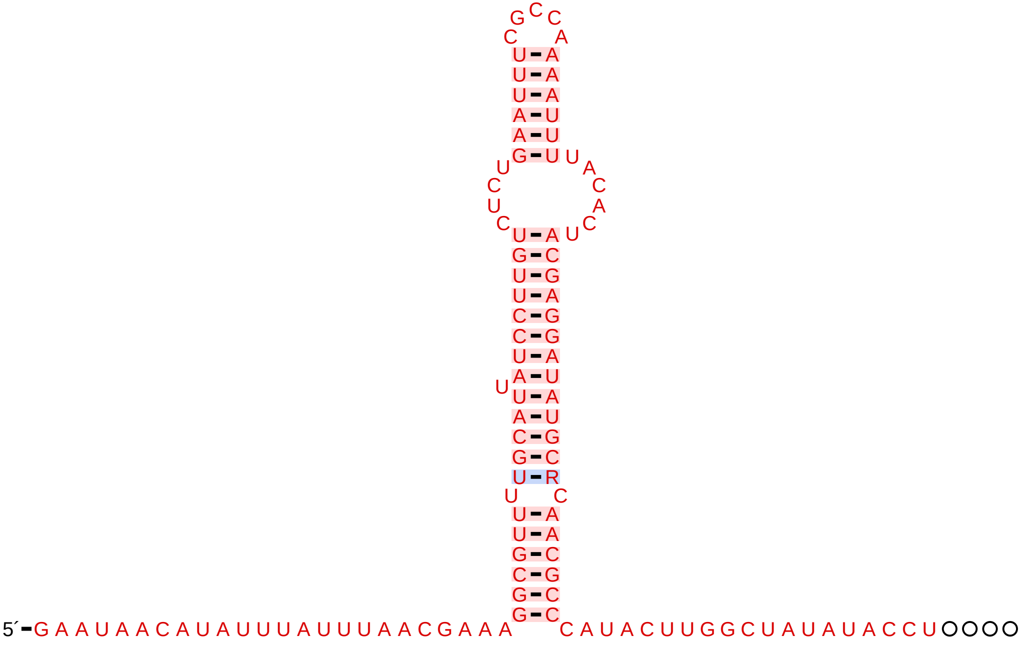

### SampleCMeta_150_1

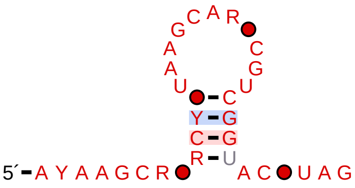

SampleCMeta_151_1

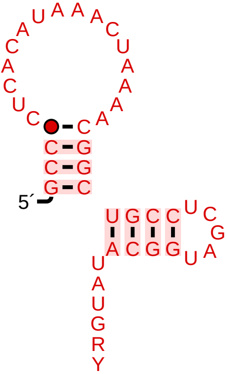

### SampleCMeta_152_1

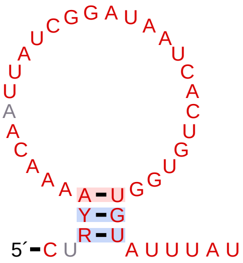

SampleCMeta_154_1

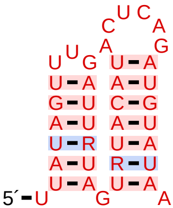

### SampleCMeta_159_1

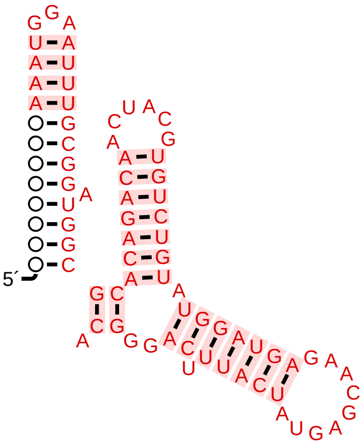

### SampleCMeta_16_1

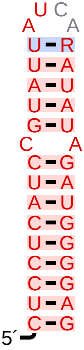

### SampleCMeta_161_1

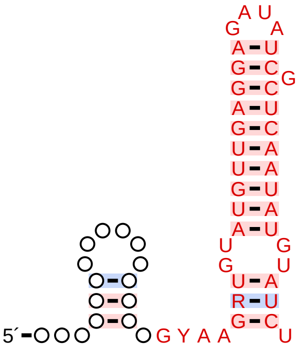

### SampleCMeta_163_1

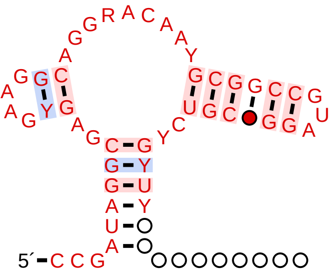

### SampleCMeta_165_1

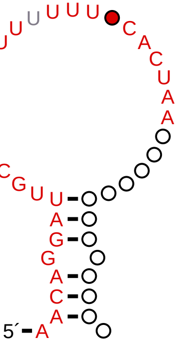

SampleCMeta_17_1

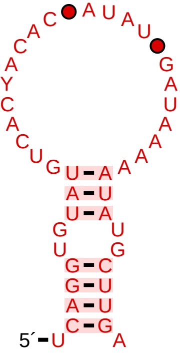

SampleCMeta_170_1

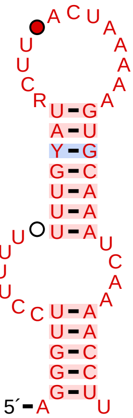

### SampleCMeta_171_1

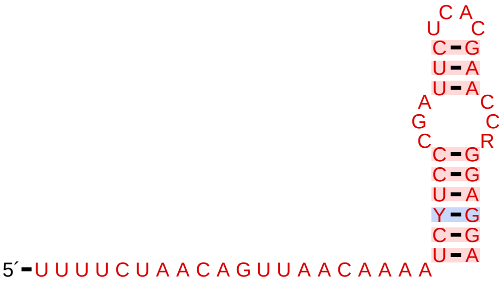

### SampleCMeta_174_1

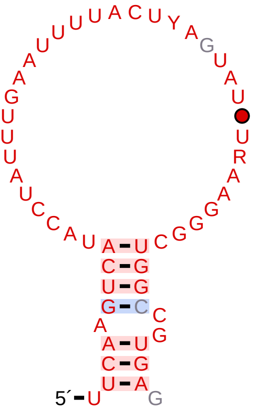

### SampleCMeta_175_1

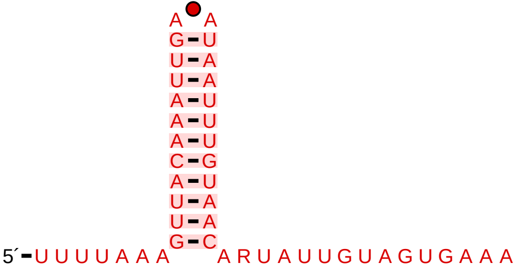

### SampleCMeta_176_1

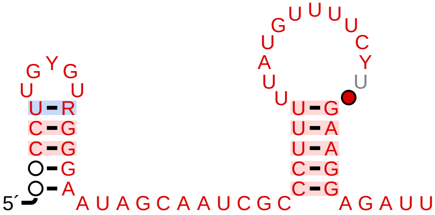

### SampleCMeta_179_1

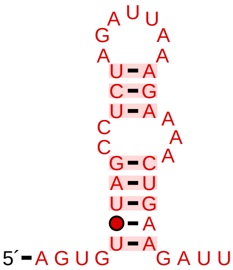

SampleCMeta_180_1

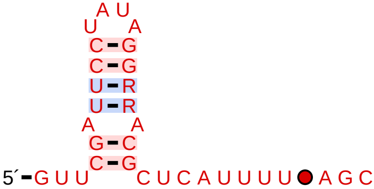

### SampleCMeta_182_1

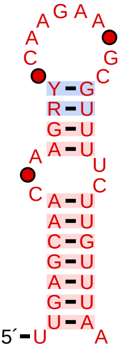

### SampleCMeta_183_1

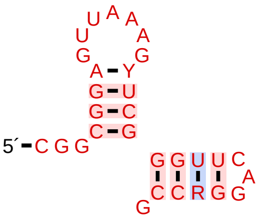

### SampleCMeta_187_1

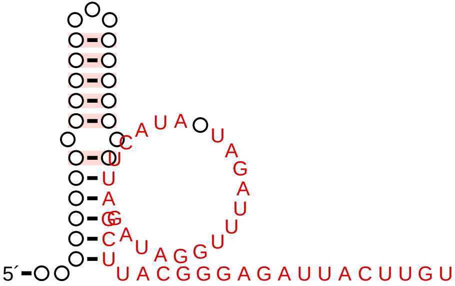

### SampleCMeta_189_1

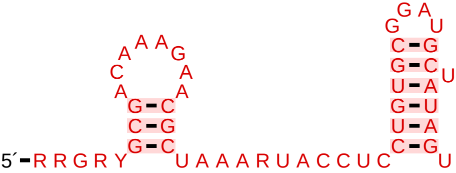

### SampleCMeta_19_1

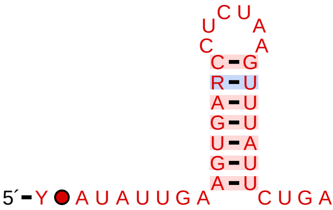

### SampleCMeta_190_1

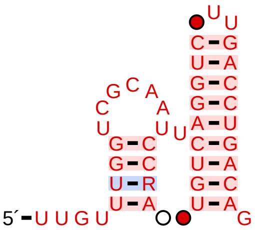

### SampleCMeta_191_1

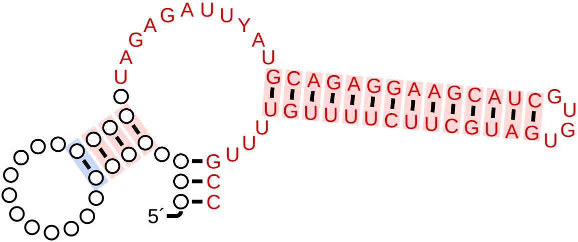

### SampleCMeta_192_1

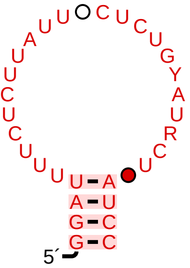

SampleCMeta_193_1

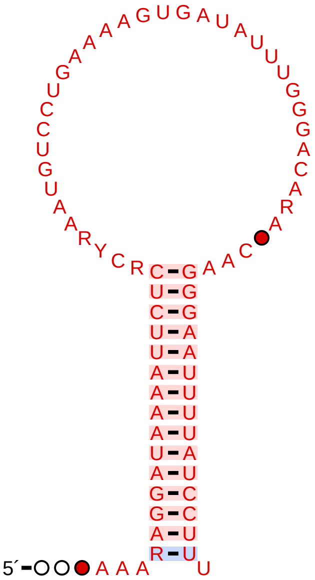

### SampleCMeta_197_1

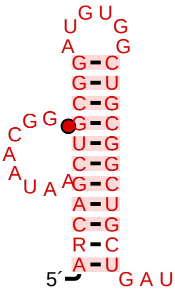

### SampleCMeta_199_1

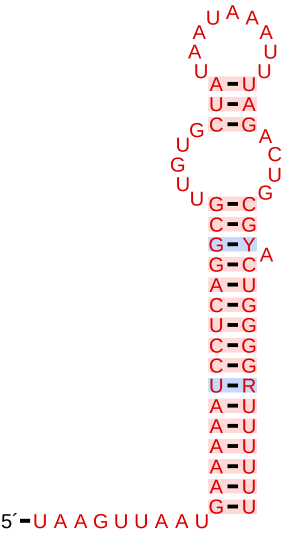

### SampleCMeta_20_1

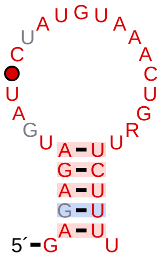

### SampleCMeta_200_1

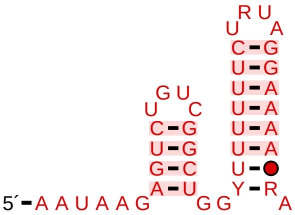

### SampleCMeta_201_1

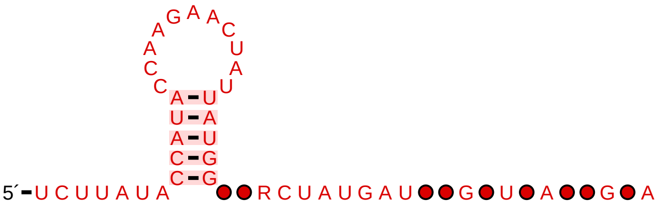

### SampleCMeta_202_1

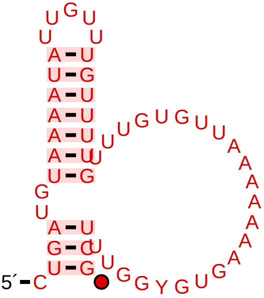

### SampleCMeta_203_1

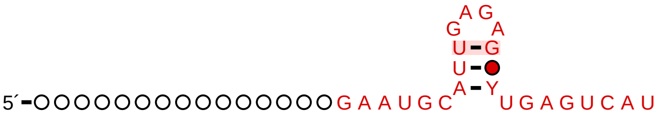

### SampleCMeta_206_1

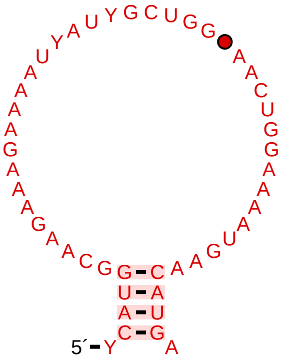

SampleCMeta_208_1

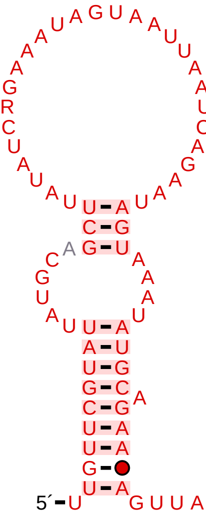

SampleCMeta_211_1

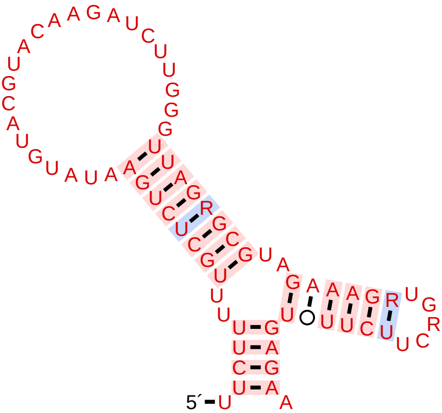

### SampleCMeta_212_1

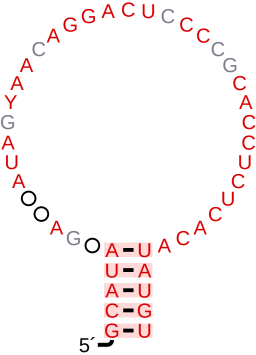

### SampleCMeta_213_1

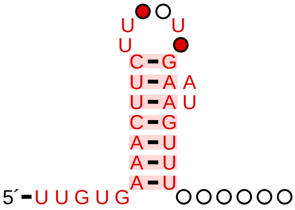

### SampleCMeta_215_1

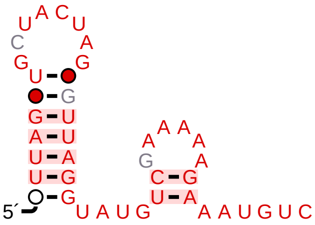

### SampleCMeta_218_1

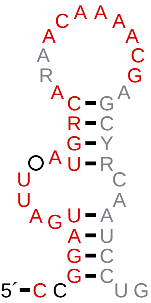

### SampleCMeta_22_1

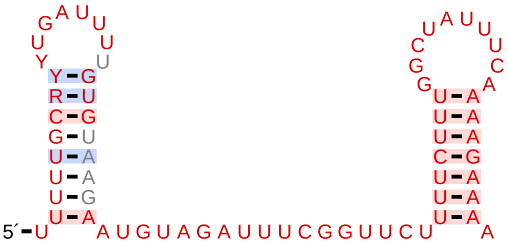

### SampleCMeta_221_1

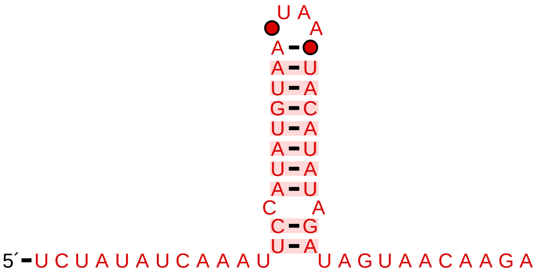

### SampleCMeta_222_1

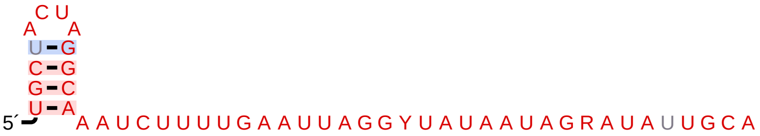

### SampleCMeta_224_1

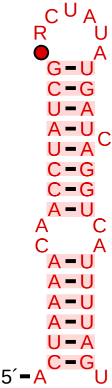

### SampleCMeta_225_1

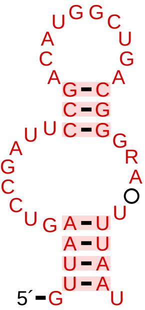

SampleCMeta_229_1

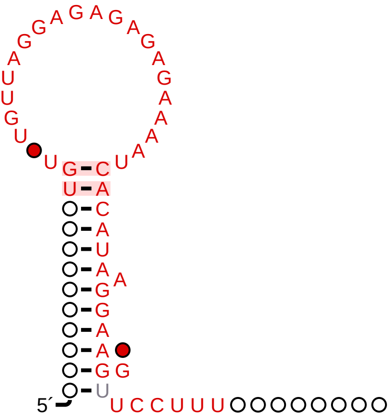

SampleCMeta_23_1

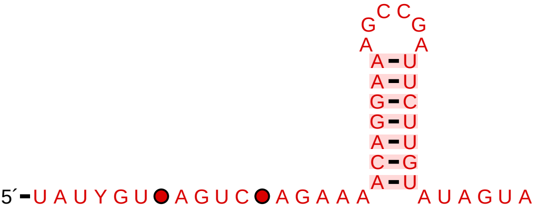

### SampleCMeta_230_1

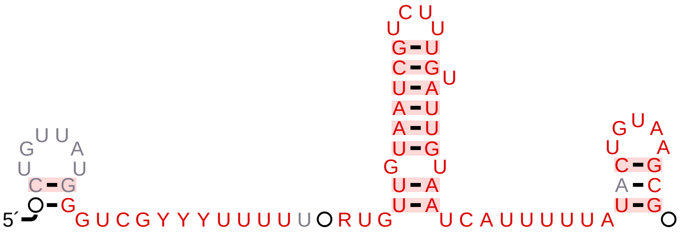

### SampleCMeta_232_1

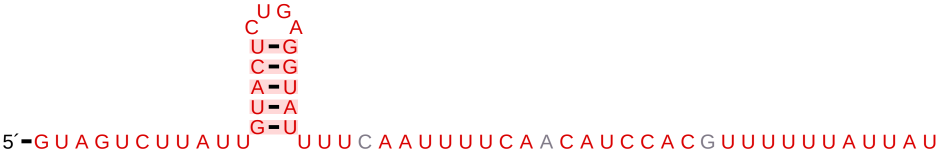

### SampleCMeta_233_1

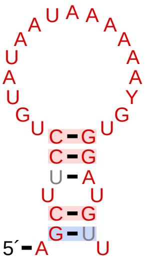

### SampleCMeta_234_1

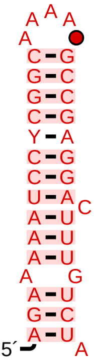

### SampleCMeta_235_1

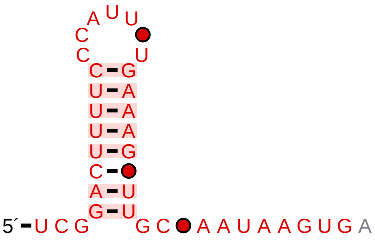

### SampleCMeta_242_1

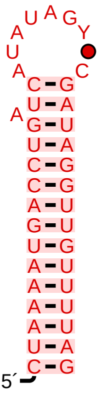

### SampleCMeta_244_1

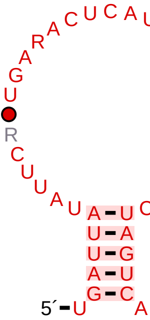

### SampleCMeta_247_1

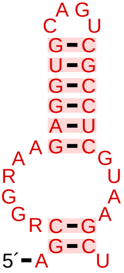

### SampleCMeta_248_1

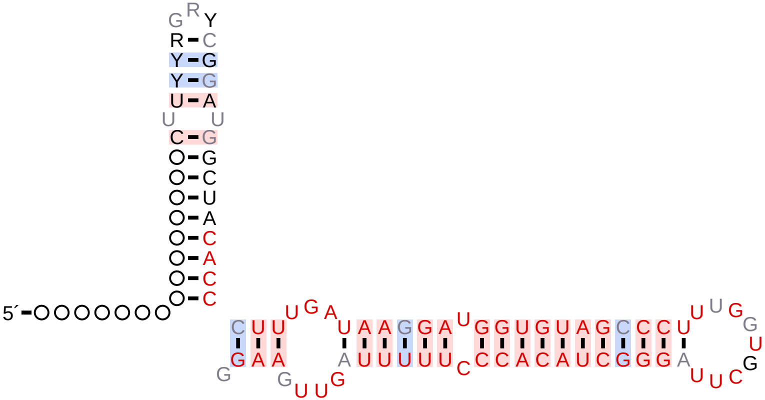

### SampleCMeta_249_1

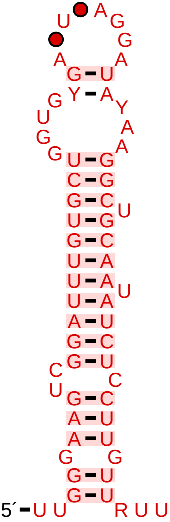

### SampleCMeta_254_1

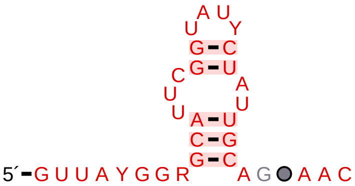

### SampleCMeta_258_1

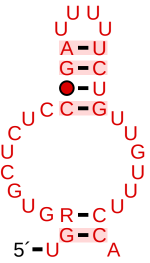

### SampleCMeta_26_1

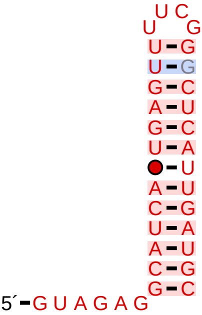

### SampleCMeta_260_1

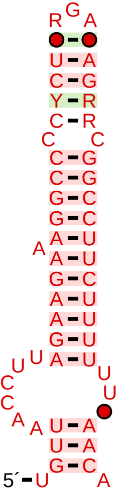

SampleCMeta_262_1

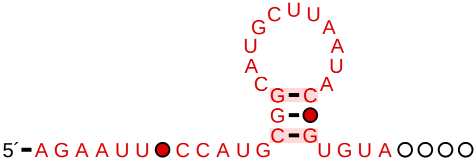

### SampleCMeta_265_1

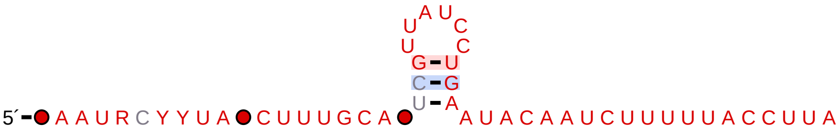

### SampleCMeta_266_1

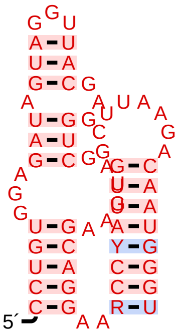

### SampleCMeta_267_1

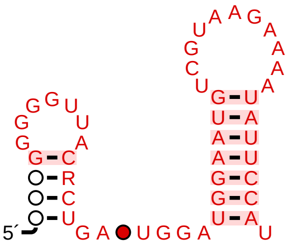

### SampleCMeta_27_1

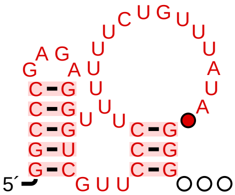

### SampleCMeta_271_1

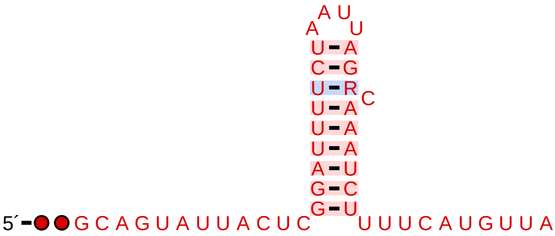

### SampleCMeta_275_1

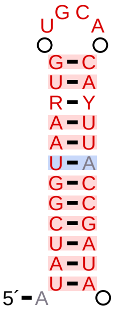

### SampleCMeta_276_1

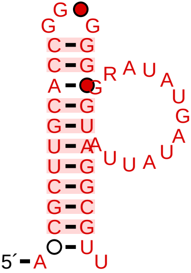

### SampleCMeta_283_1

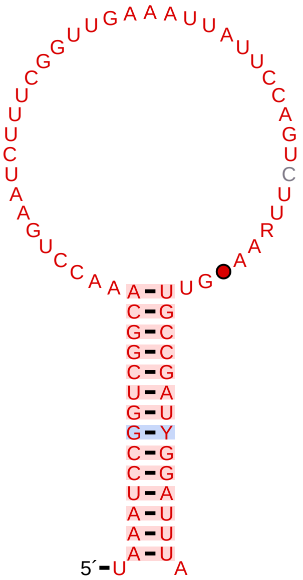

### SampleCMeta_285_1

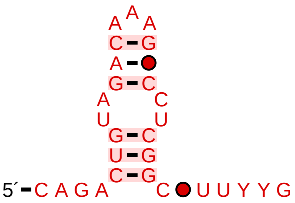

### SampleCMeta_288_1

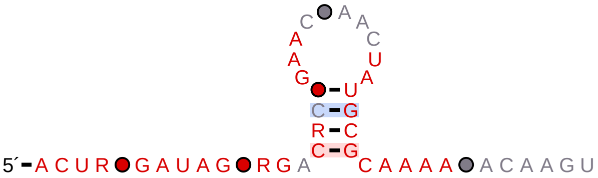

### SampleCMeta_289_1

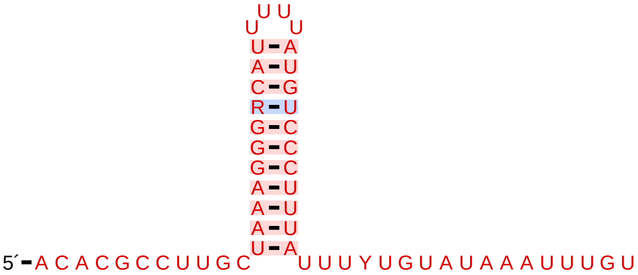

SampleCMeta_29_1

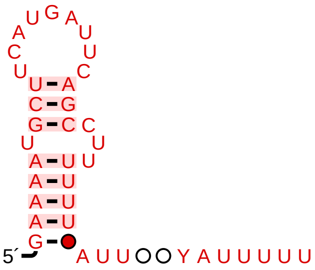

### SampleCMeta_291_1

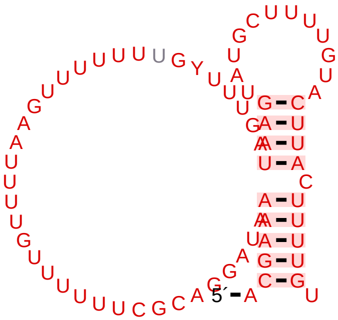

### SampleCMeta_293_1

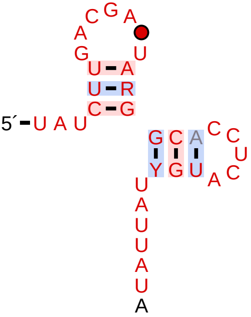

### SampleCMeta_294_1

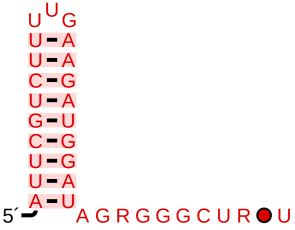

### SampleCMeta_301_1

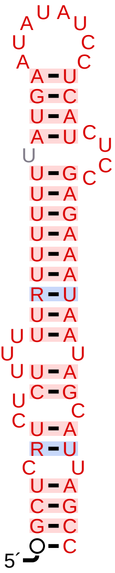

### SampleCMeta_302_1

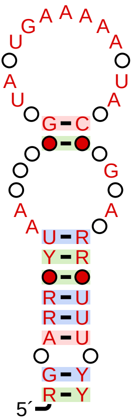

### SampleCMeta_305_1

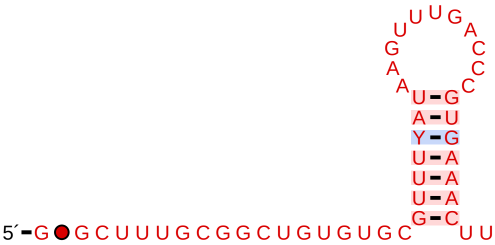

### SampleCMeta_306_1

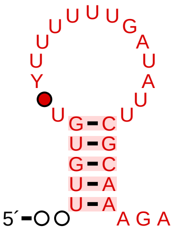

### SampleCMeta_310_1

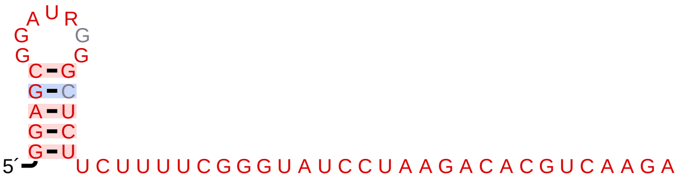

### SampleCMeta_313_1

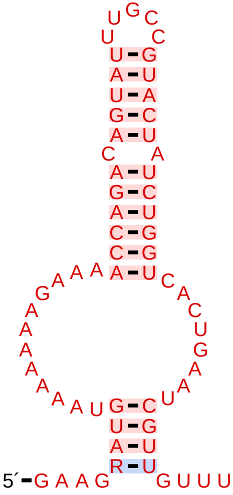

### SampleCMeta_319_1

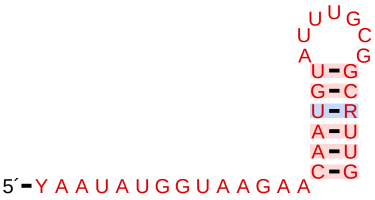

### SampleCMeta_32_1

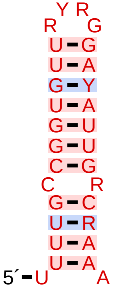

SampleCMeta_321_1

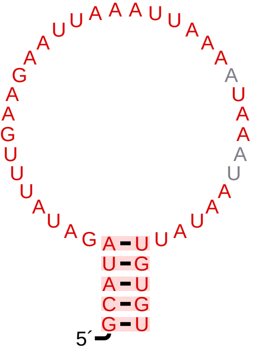

### SampleCMeta_323_1

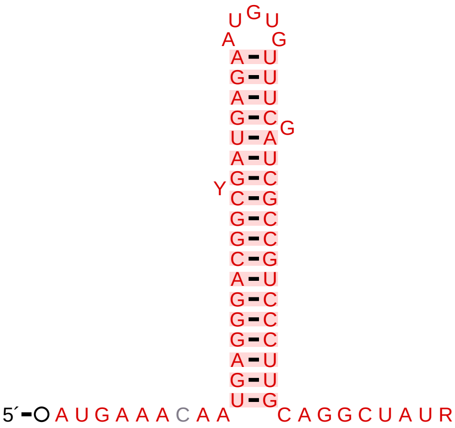

### SampleCMeta_325_1

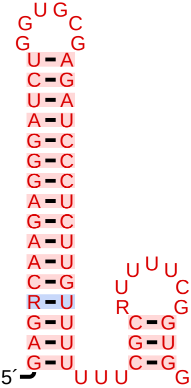

### SampleCMeta_326_1

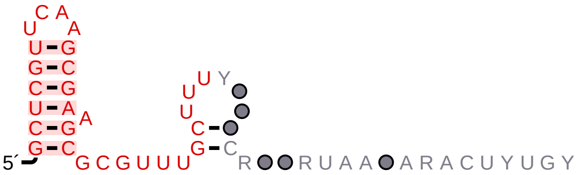

### SampleCMeta_329_1

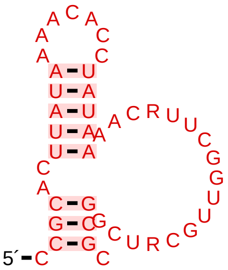

### SampleCMeta_333_1

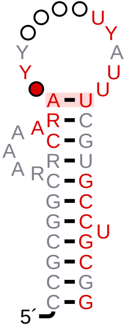

### SampleCMeta_338_1

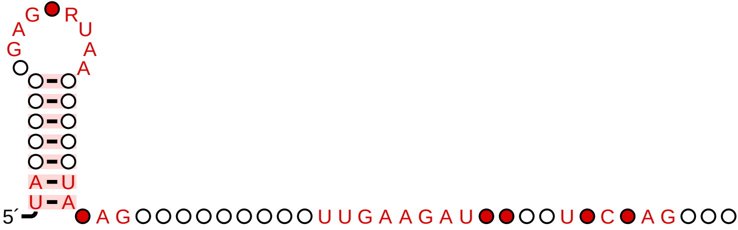

SampleCMeta_340_1

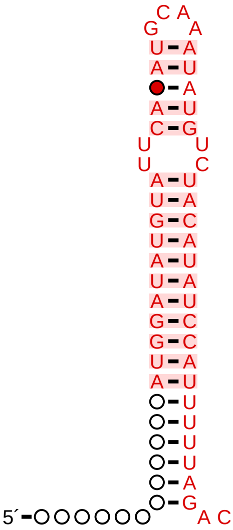

### SampleCMeta_344_1

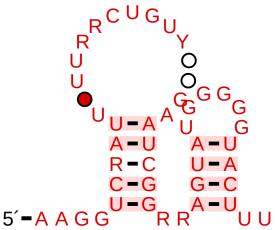

### SampleCMeta_346_1

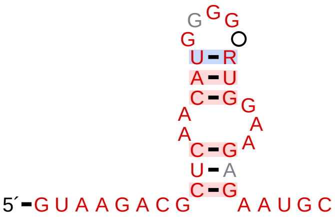

### SampleCMeta_347_1

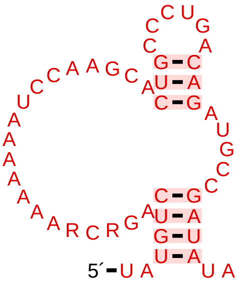

SampleCMeta_35_1

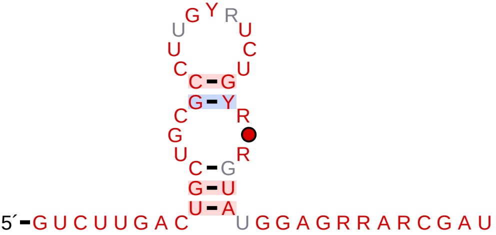

### SampleCMeta_353_1

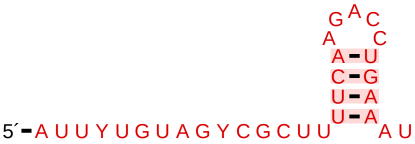

### SampleCMeta_354_1

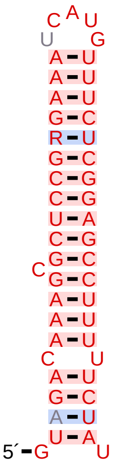

### SampleCMeta_356_1

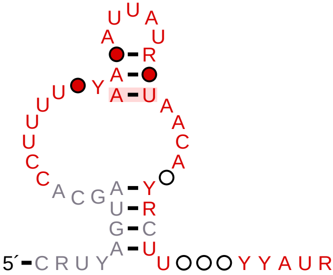

### SampleCMeta_357_1

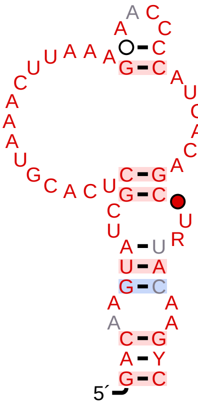

### SampleCMeta_358_1

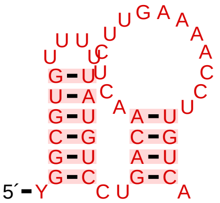

### SampleCMeta_360_1

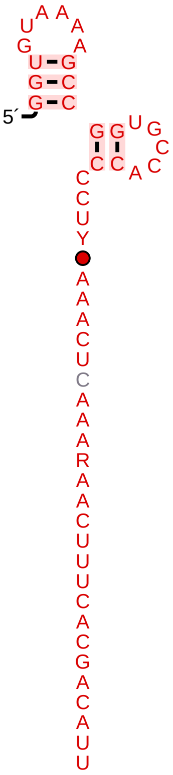

### SampleCMeta_361_1

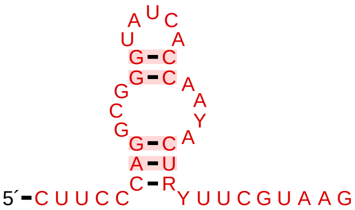

### SampleCMeta_366_1

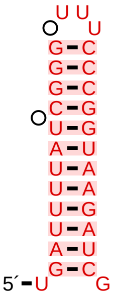

### SampleCMeta_367_1

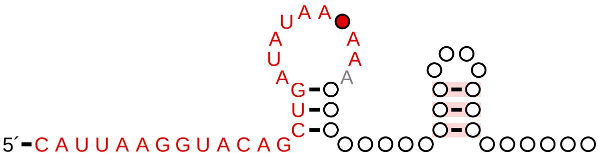

SampleCMeta_368_1

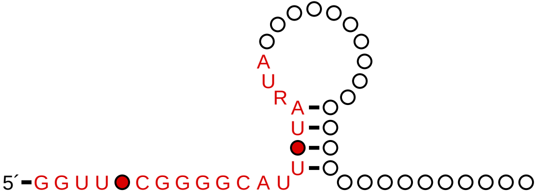

### SampleCMeta_370_1

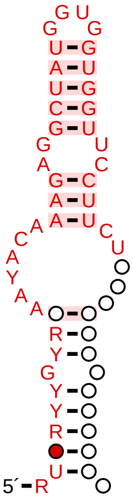

### SampleCMeta_374_1

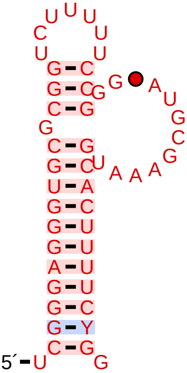

### SampleCMeta_375_1

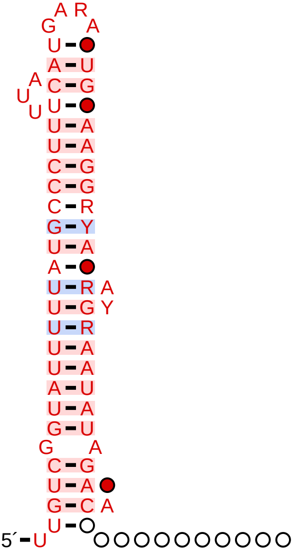

### SampleCMeta_377_1

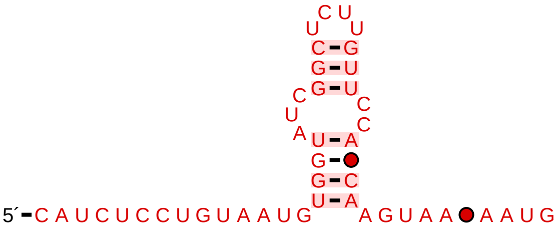

### SampleCMeta_378_1

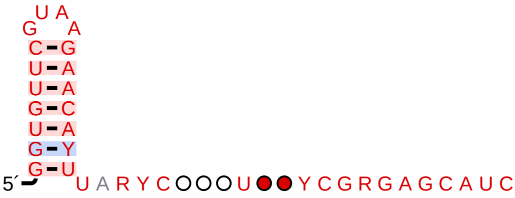

### SampleCMeta_380_1

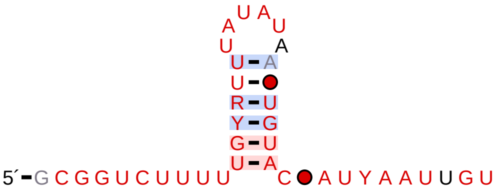

### SampleCMeta_381_1

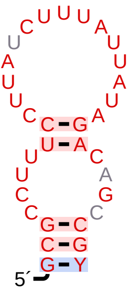

### SampleCMeta_384_1

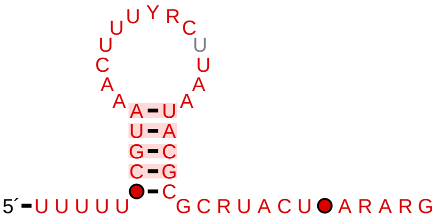

SampleCMeta_385_1

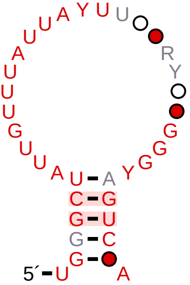

### SampleCMeta_386_1

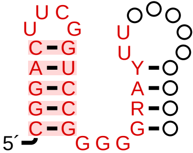

### SampleCMeta_387_1

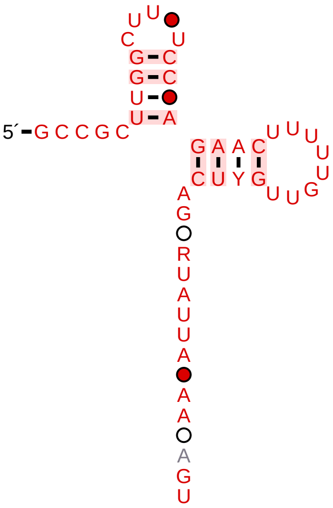

### SampleCMeta_391_1

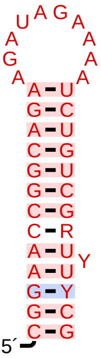

### SampleCMeta_393_1

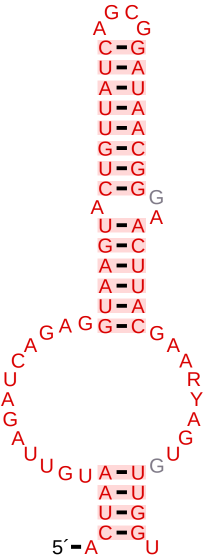

### SampleCMeta_394_1

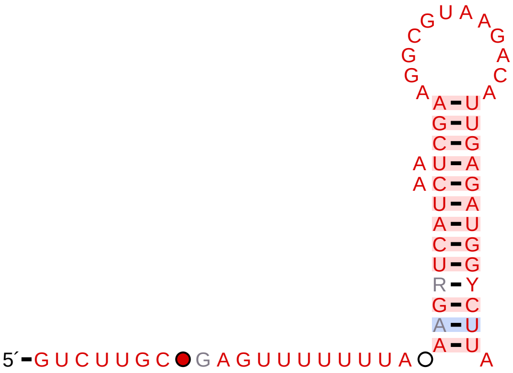

### SampleCMeta_396_1

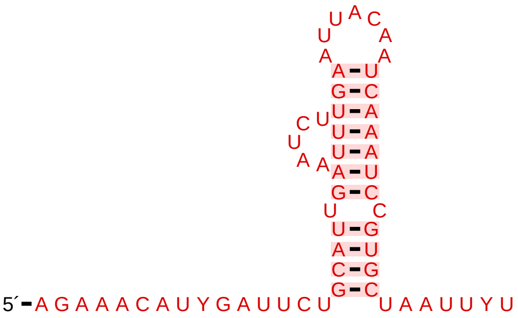

### SampleCMeta_398_1

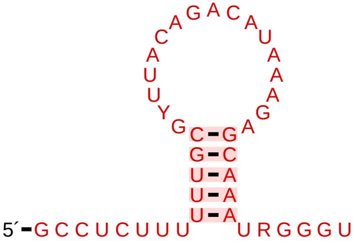

### SampleCMeta_399_1

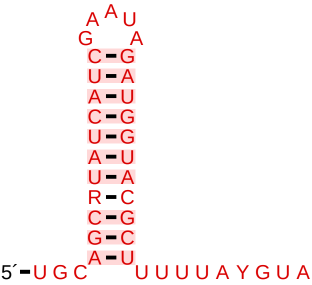

SampleCMeta_40_1

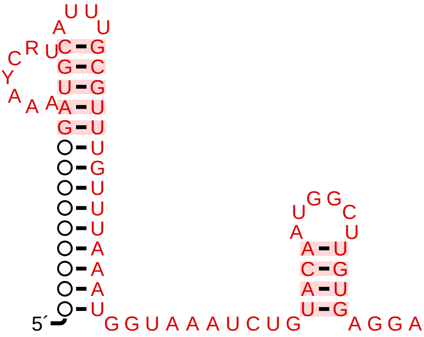

### SampleCMeta_400_1

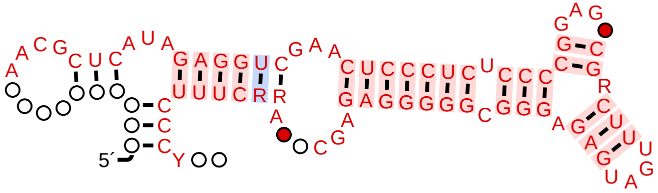

### SampleCMeta_403_1

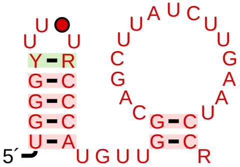

SampleCMeta_404_1

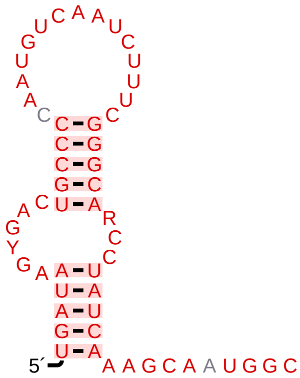

### SampleCMeta_406_1

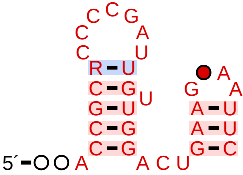

### SampleCMeta_407_1

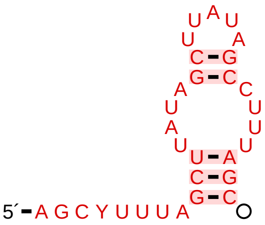

### SampleCMeta_413_1

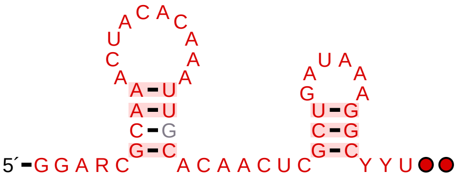

### SampleCMeta_414_1

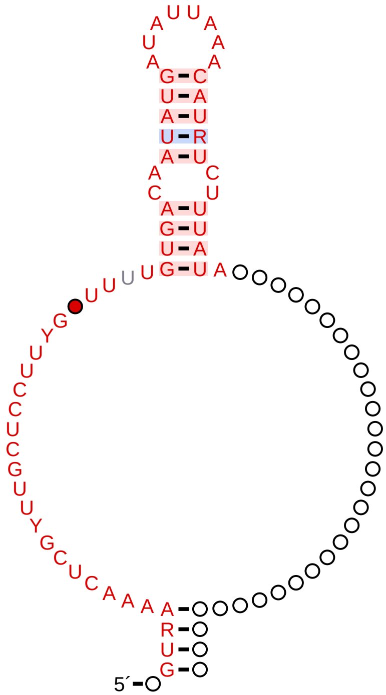

### SampleCMeta_415_1

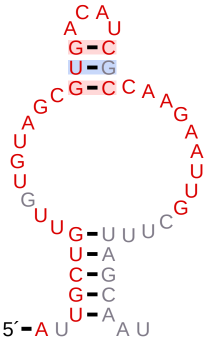

### SampleCMeta_418_1

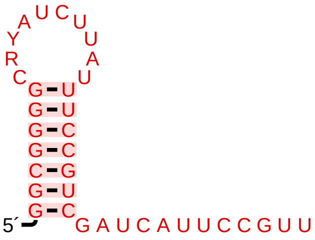

### SampleCMeta_42_1

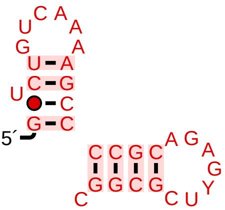

### SampleCMeta_420_1

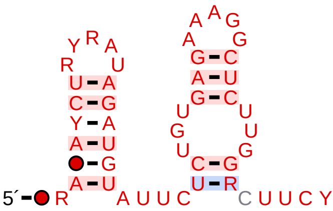

### SampleCMeta_421_1

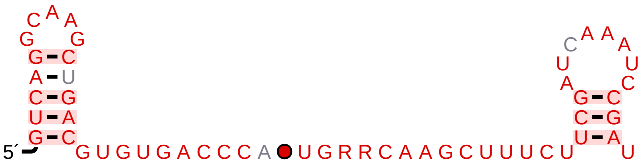

### SampleCMeta_424_1

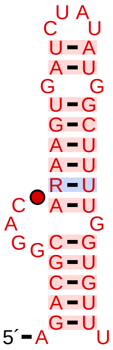

SampleCMeta_427_1

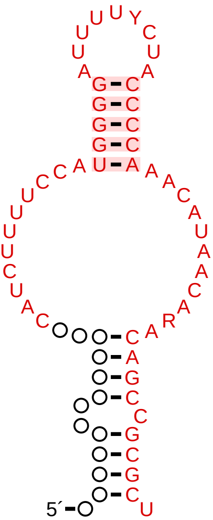

SampleCMeta_43_1

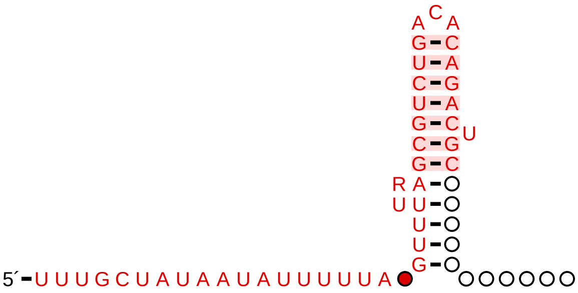

### SampleCMeta_431_1

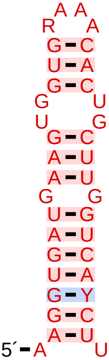

### SampleCMeta_432_1

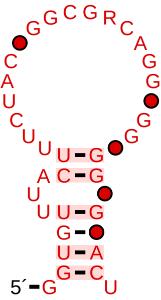

### SampleCMeta_436_1

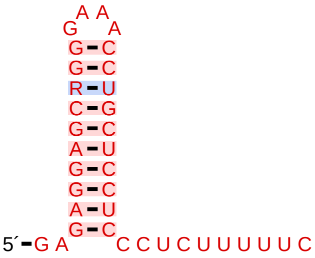

SampleCMeta_438_1

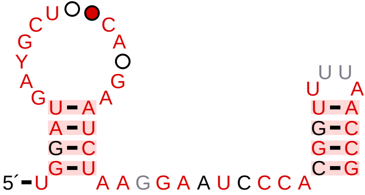

### SampleCMeta_440_1

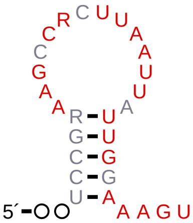

### SampleCMeta_441_1

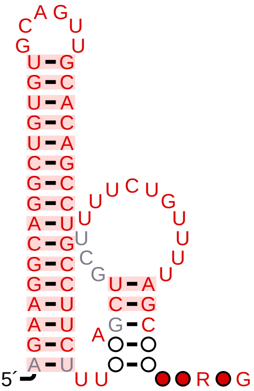

### SampleCMeta_444_1

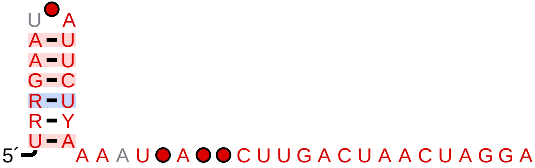

### SampleCMeta_445_1

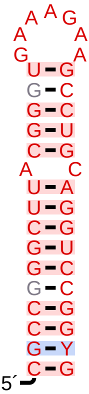

### SampleCMeta_449_1

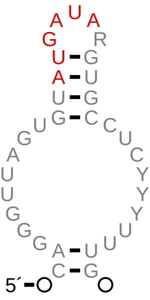

### SampleCMeta_45_1

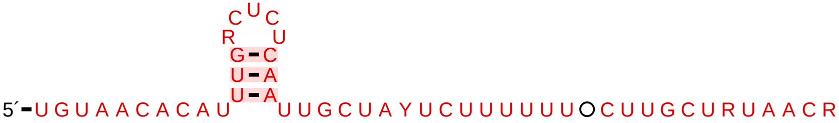

### SampleCMeta_450_1

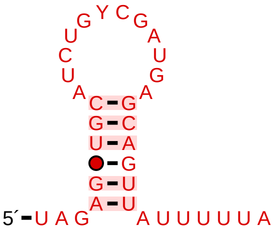

SampleCMeta_453_1

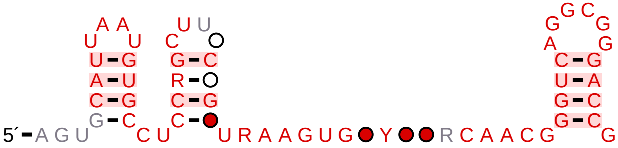

SampleCMeta_457_1

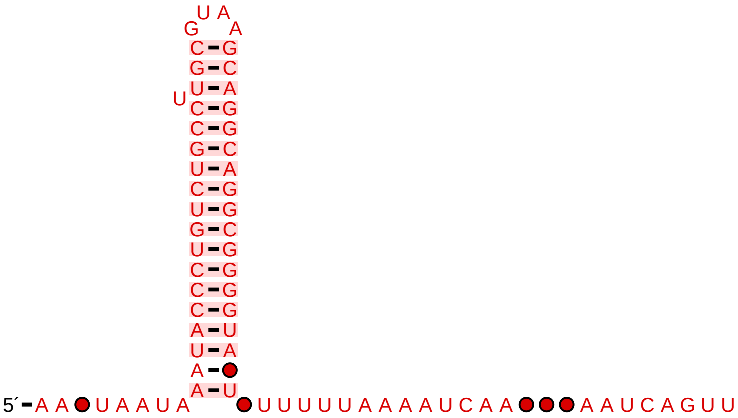

### SampleCMeta_459_1

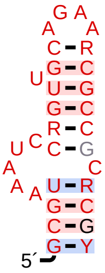

### SampleCMeta_460_1

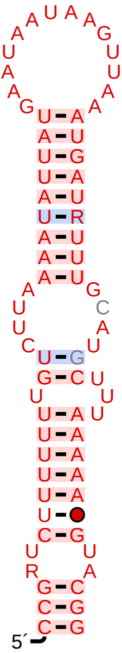

### SampleCMeta_461_1

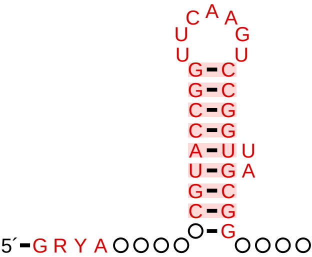

### SampleCMeta_464_1

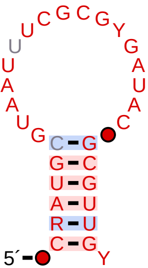

### SampleCMeta_467_1

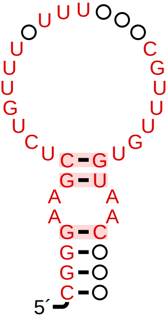

### SampleCMeta_469_1

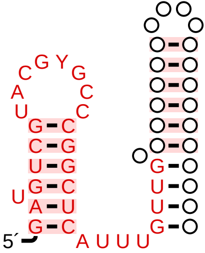

### SampleCMeta_47_1

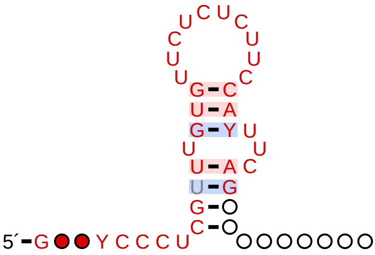

### SampleCMeta_470_1

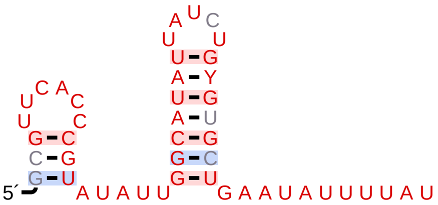

### SampleCMeta_471_1

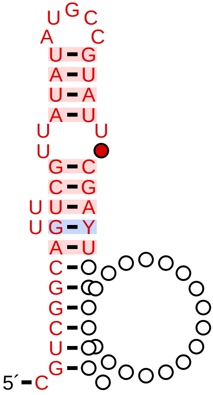

### SampleCMeta_472_1

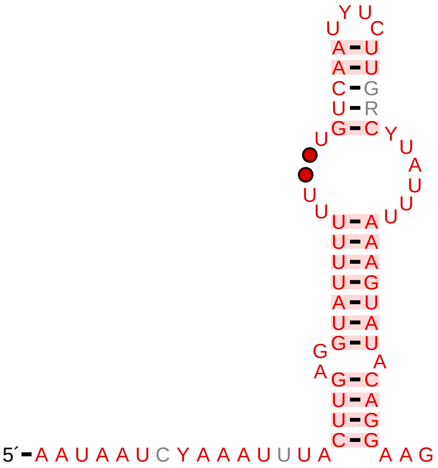

### SampleCMeta_473_1

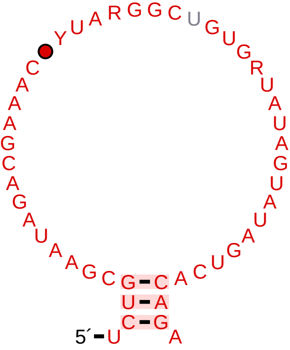

SampleCMeta_478_1

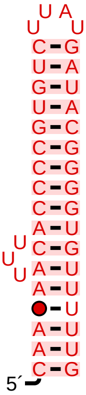

### SampleCMeta_480_1

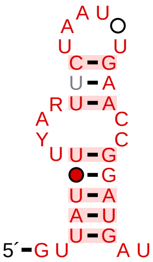

SampleCMeta_481_1

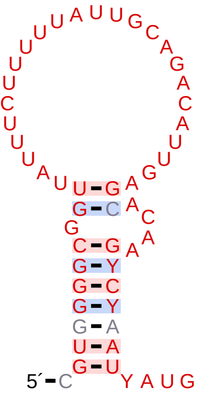

### SampleCMeta_482_1

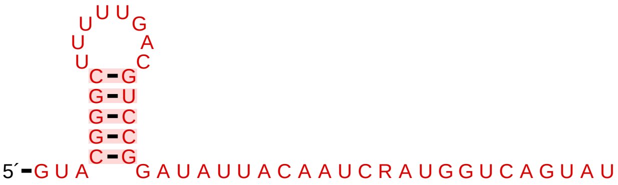

### SampleCMeta_483_1

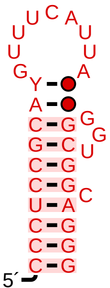

### SampleCMeta_484_1

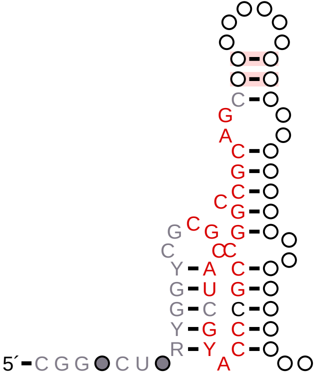

### SampleCMeta_485_1

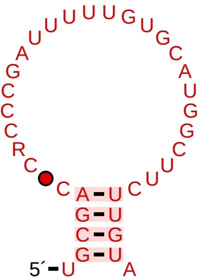

### SampleCMeta_486_1

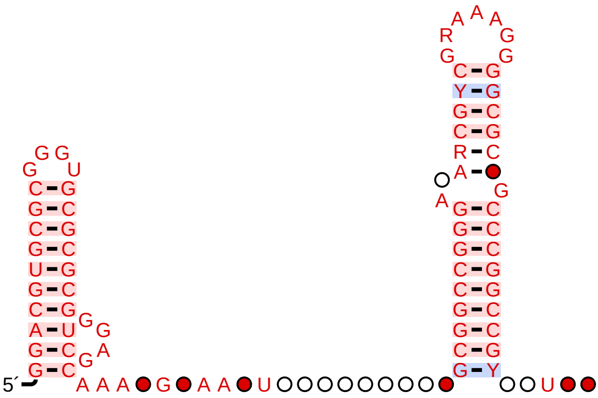

### SampleCMeta_488_1

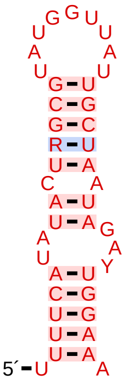

SampleCMeta_489_1

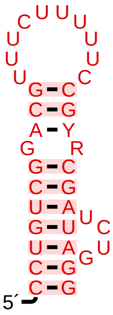

### SampleCMeta_490_1

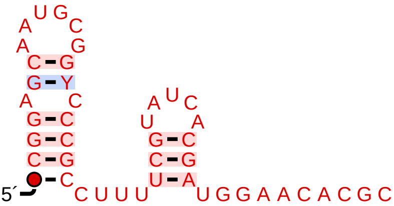

### SampleCMeta_493_1

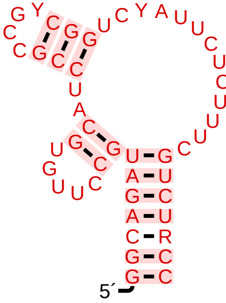

### SampleCMeta_495_1

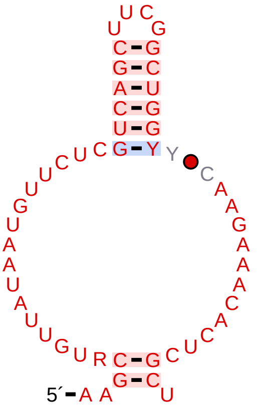

### SampleCMeta_496_1

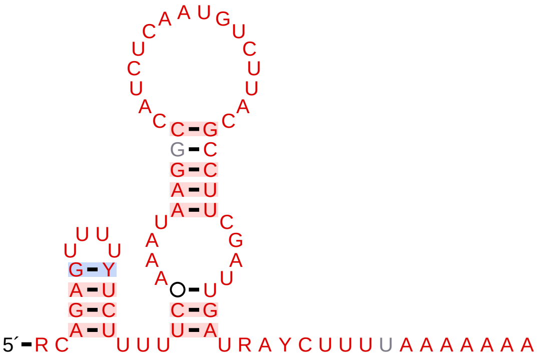

### SampleCMeta_497_1

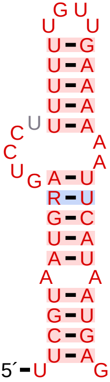

### SampleCMeta_499_1

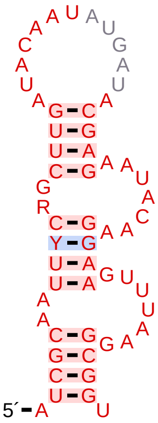

### SampleCMeta_509_1

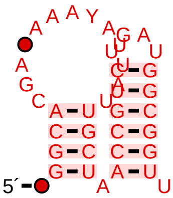

### SampleCMeta_511_1

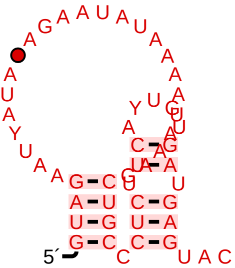

### SampleCMeta_512_1

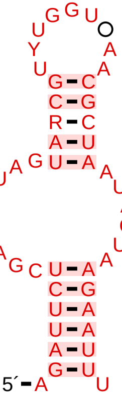

### SampleCMeta_513_1

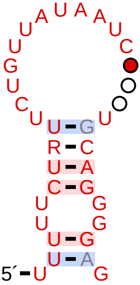

### SampleCMeta_515_1

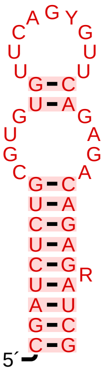

### SampleCMeta_517_1

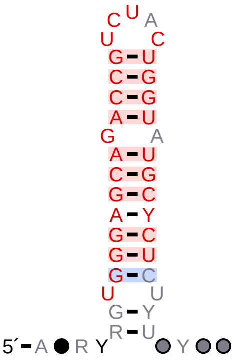

### SampleCMeta_521_1

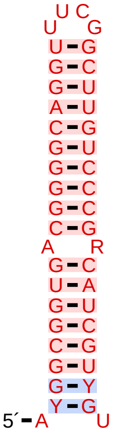

### SampleCMeta_524_1

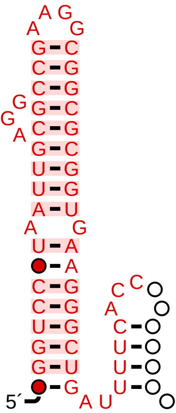

### SampleCMeta_528_1

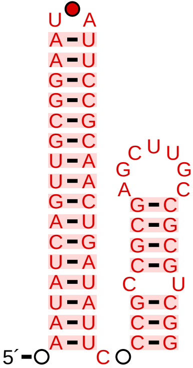

SampleCMeta_529_1

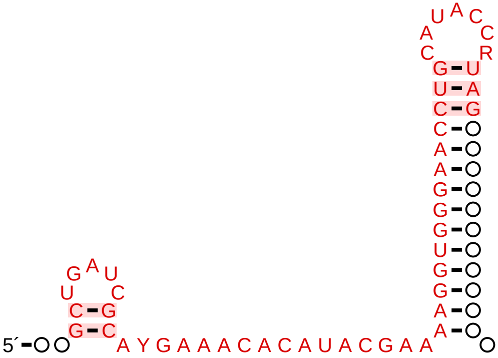

### SampleCMeta_53_1

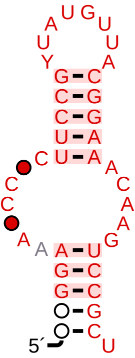

### SampleCMeta_532_1

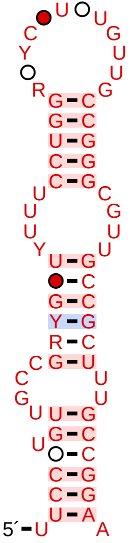

SampleCMeta_535_1

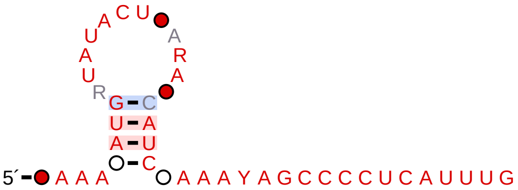

SampleCMeta_537_1

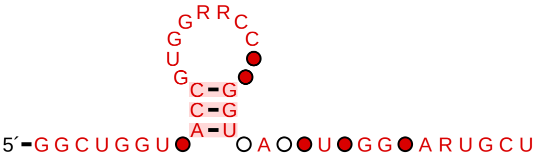

### SampleCMeta_539_1

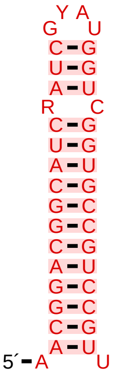

SampleCMeta_542_1

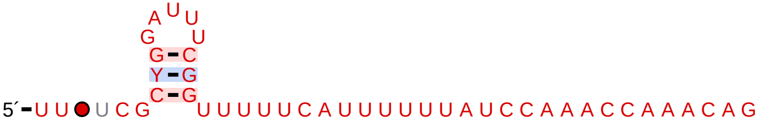

### SampleCMeta_544_1

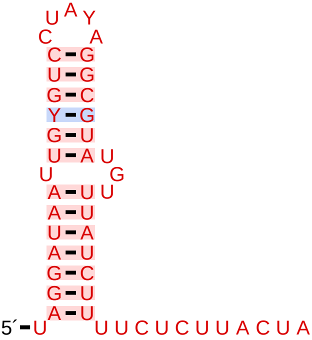

### SampleCMeta_547_1

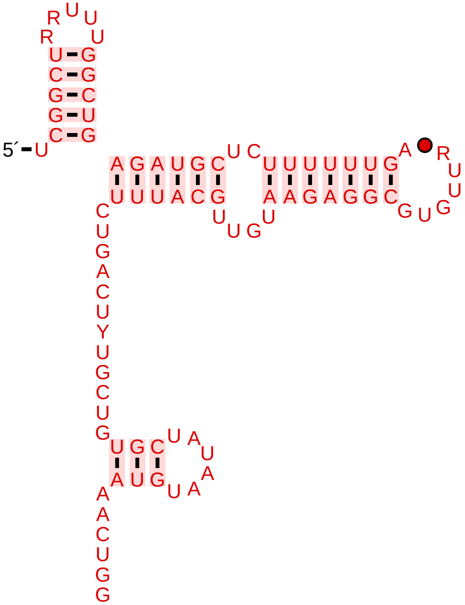

### SampleCMeta_549_1

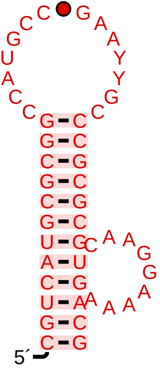

### SampleCMeta_55_1

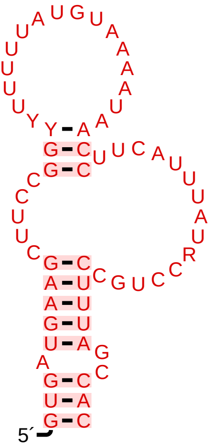

### SampleCMeta_550_1

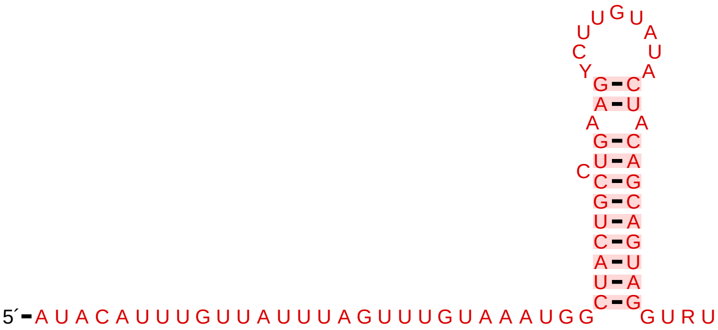

### SampleCMeta_551_1

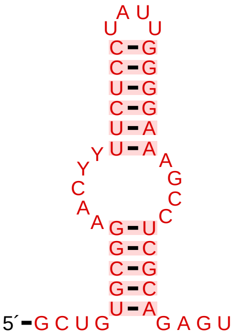

### SampleCMeta_554_1

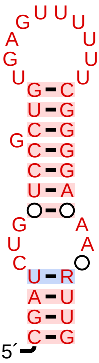

### SampleCMeta_555_1

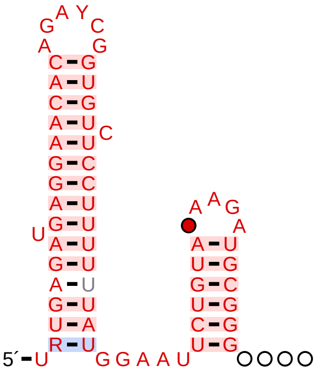

### SampleCMeta_556_1

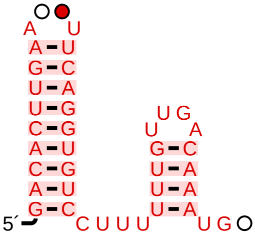

### SampleCMeta_558_1

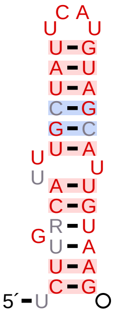

### SampleCMeta_56_1

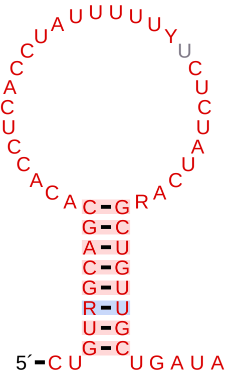

### SampleCMeta_566_1

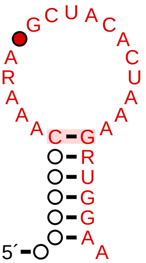

### SampleCMeta_57_1

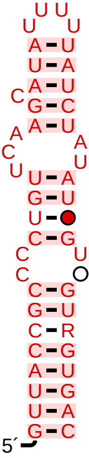

### SampleCMeta_573_1

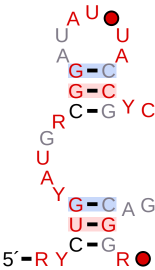

### SampleCMeta_575_1

5´-○○○AUACCC○CYUUACUGUGUUCCUGAGAGCA

### SampleCMeta_576_1

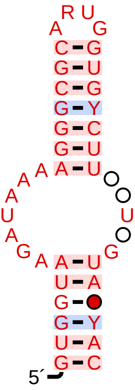

### SampleCMeta_578_1

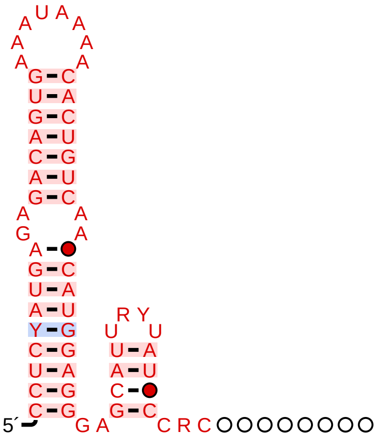

### SampleCMeta_580_1

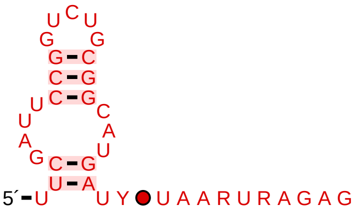

### SampleCMeta_582_1

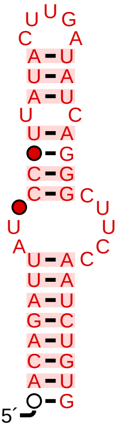

### SampleCMeta_585_1

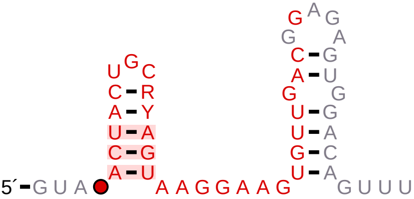

### SampleCMeta_587_1

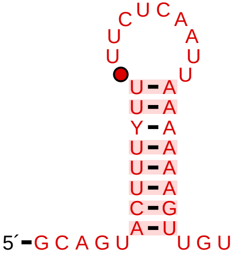

### SampleCMeta_59_1

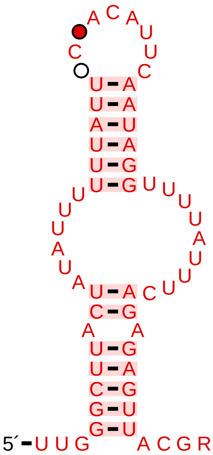

### SampleCMeta_590_1

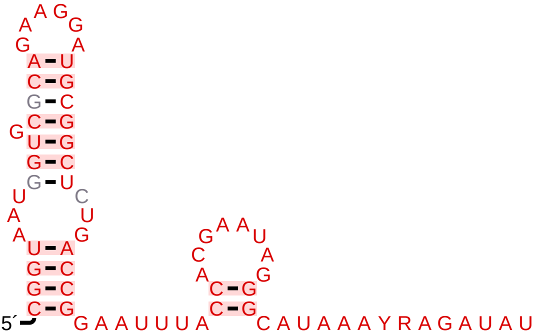

SampleCMeta_592_1

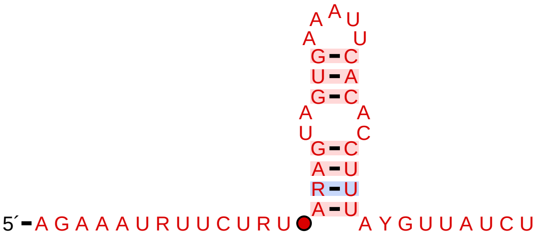

### SampleCMeta_593_1

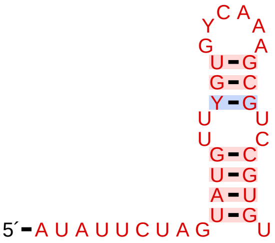

SampleCMeta_594_1

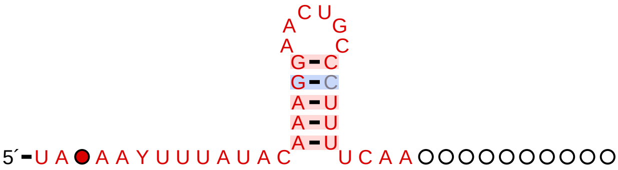

SampleCMeta_595_1

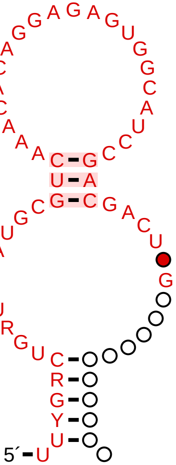

### SampleCMeta_599_1

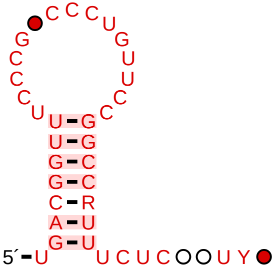

### SampleCMeta_6_1

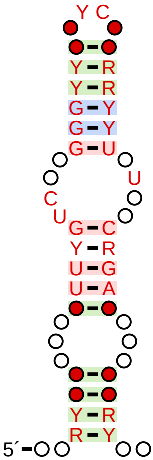

### SampleCMeta_602_1

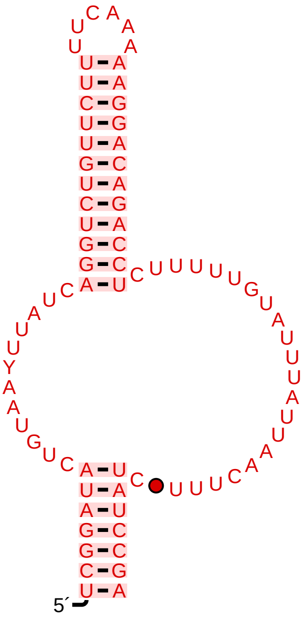

### SampleCMeta_604_1

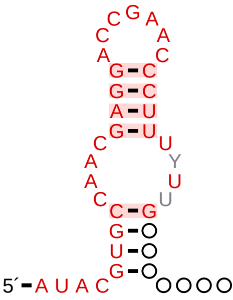

SampleCMeta_606_1

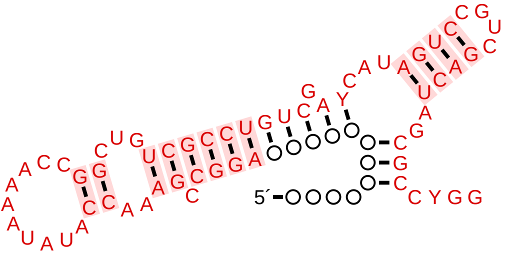

SampleCMeta_607_1

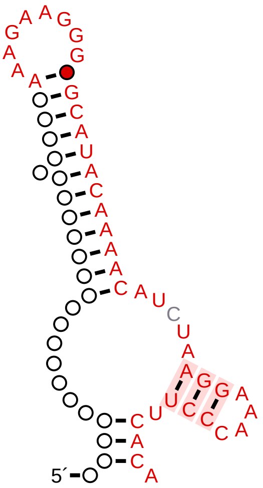

### SampleCMeta_609_1

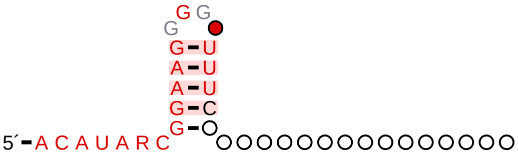

### SampleCMeta_611_1

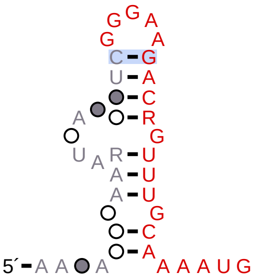

### SampleCMeta_613_1

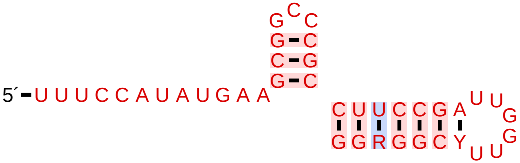

### SampleCMeta_616_1

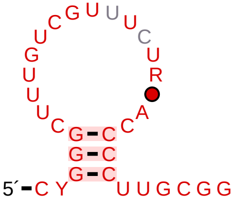

### SampleCMeta_617_1

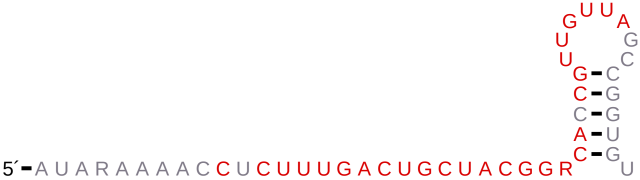

### SampleCMeta_619_1

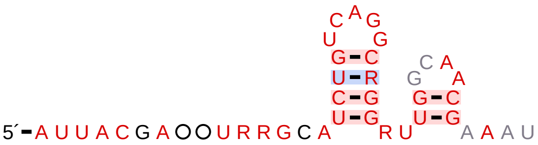

### SampleCMeta_62_1

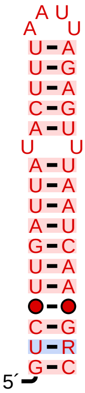

SampleCMeta_620_1

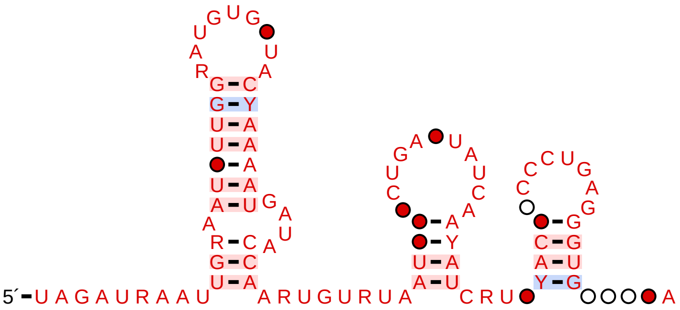

### SampleCMeta_621_1

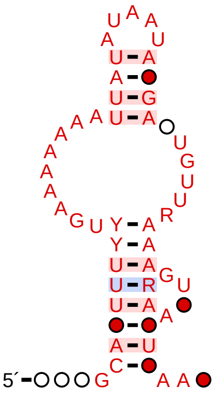

### SampleCMeta_622_1

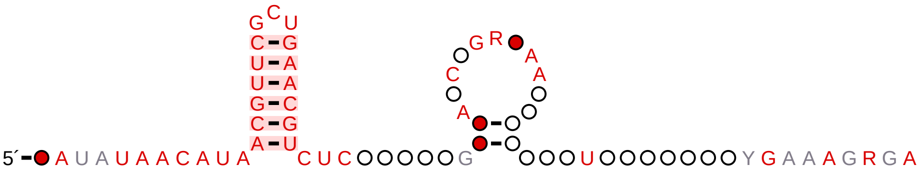

### SampleCMeta_64_1

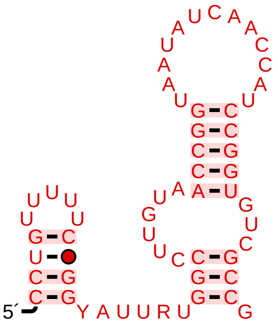

### SampleCMeta_65_1

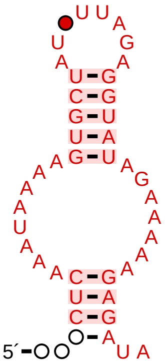

### SampleCMeta_7_1

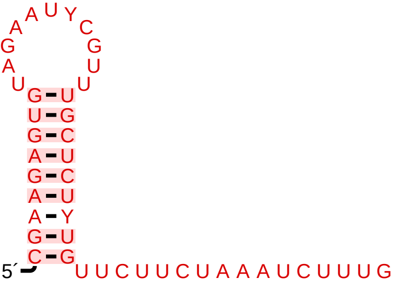

### SampleCMeta_71_1

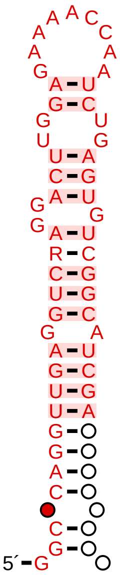

### SampleCMeta_74_1

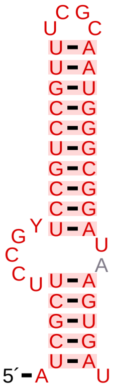

### SampleCMeta_79_1

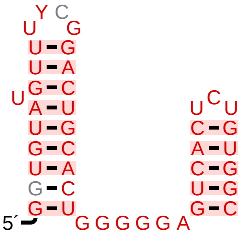

SampleCMeta_8_1

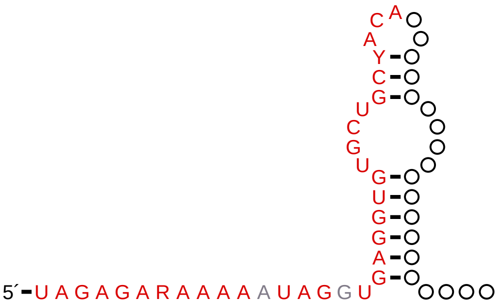

### SampleCMeta_81_1

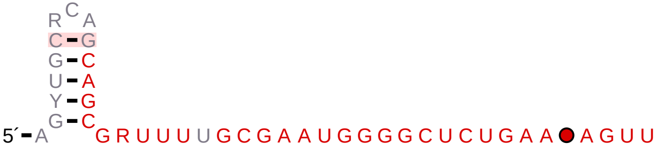

### SampleCMeta_91_1

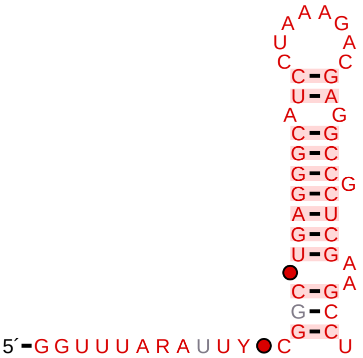

### SampleCMeta_95_1

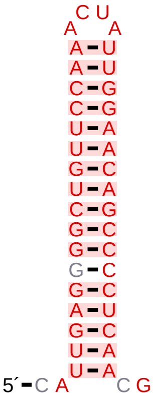

### SampleDMeta_1_1

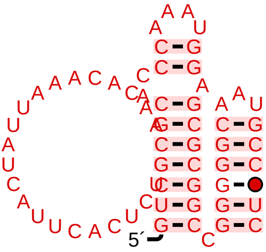

### SampleDMeta_105_1

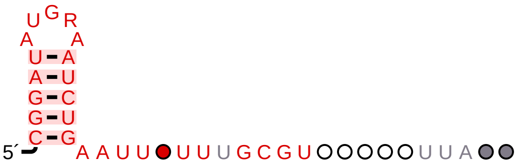

### SampleDMeta_107_1

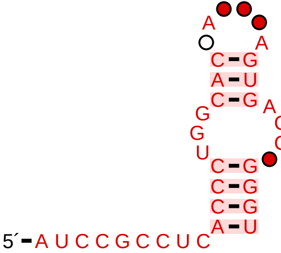

SampleDMeta_108_1

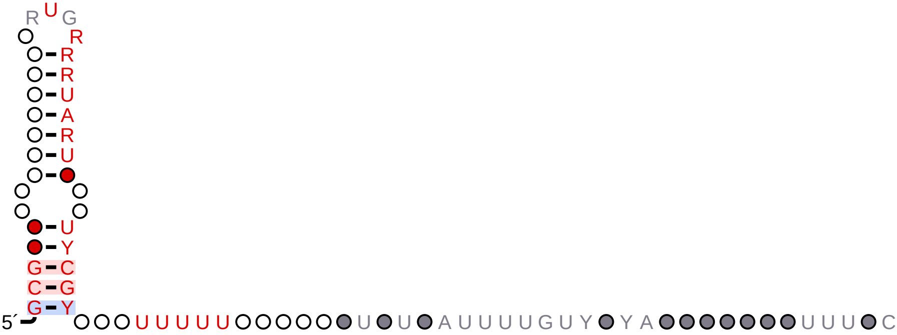

### SampleDMeta_112_1

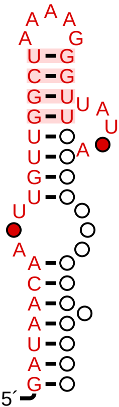

SampleDMeta_114_1

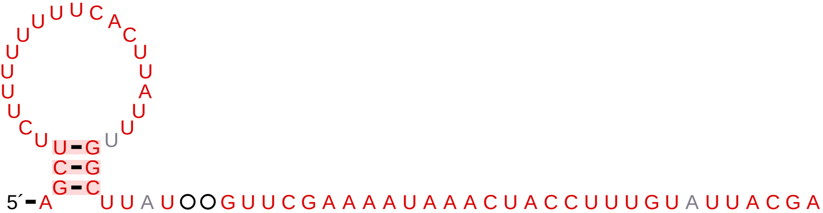

### SampleDMeta_124_1

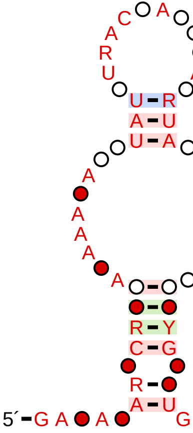

### SampleDMeta_126_1

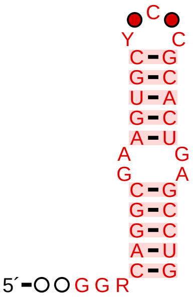

### SampleDMeta_127_1

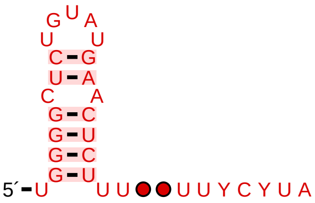

### SampleDMeta_128_1

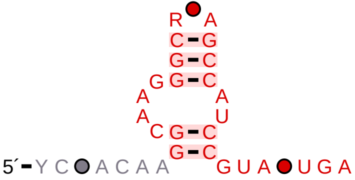

### SampleDMeta_129_1

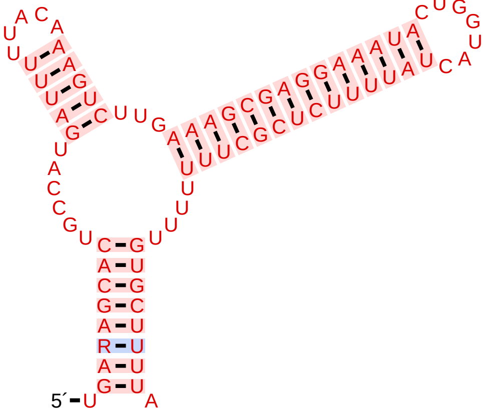

### SampleDMeta_134_1

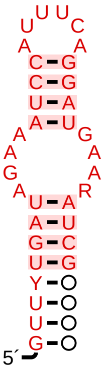

SampleDMeta_135_1

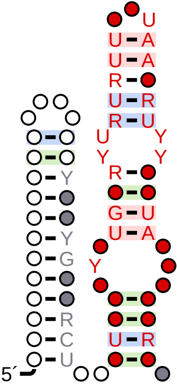

SampleDMeta_138_1

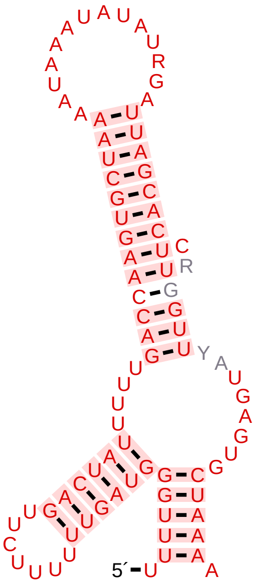

SampleDMeta_14_1

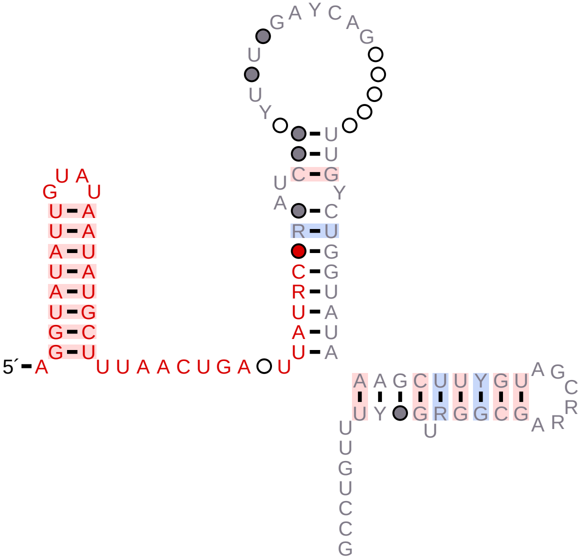

### SampleDMeta_143_1

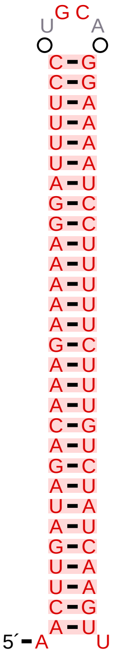

### SampleDMeta_151_1

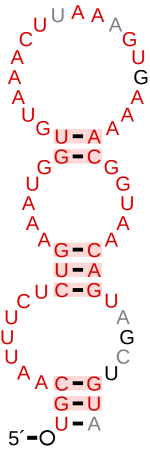

### SampleDMeta_153_1

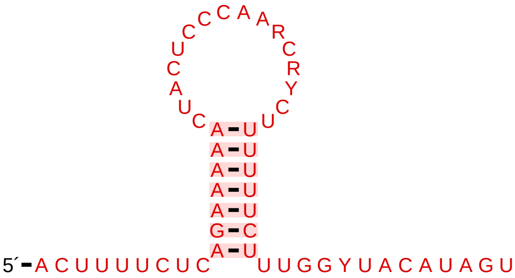

### SampleDMeta_157_1

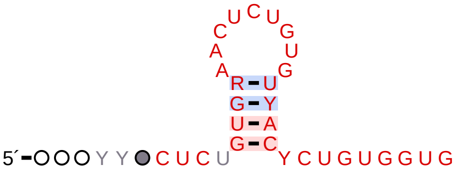

### SampleDMeta_158_1

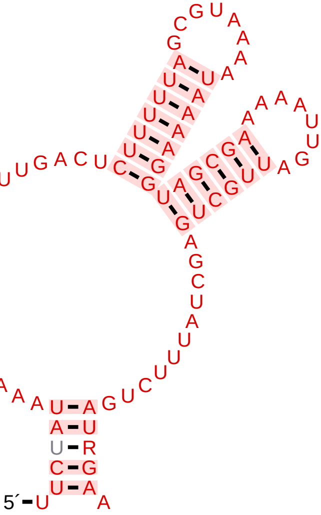

### SampleDMeta_159_1

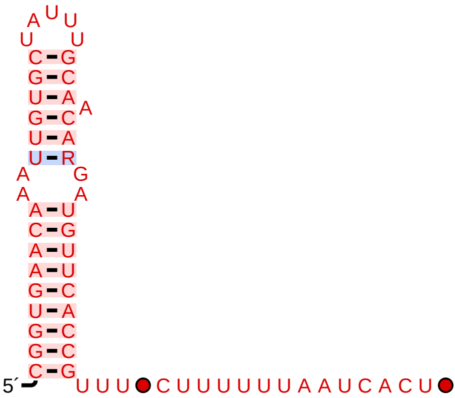

### SampleDMeta_160_1

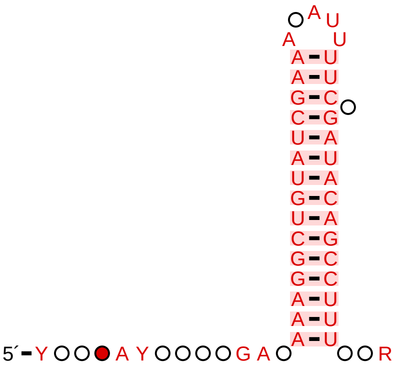

SampleDMeta_163_1

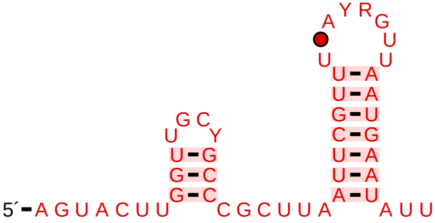

### SampleDMeta_164_1

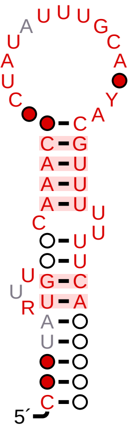

SampleDMeta_167_1

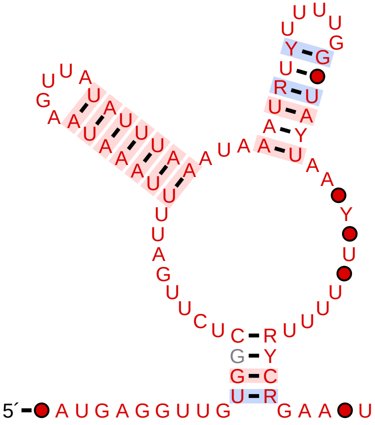

### SampleDMeta_168_1

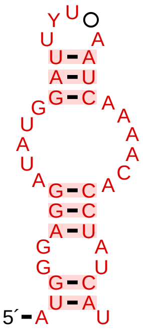

### SampleDMeta_169_1

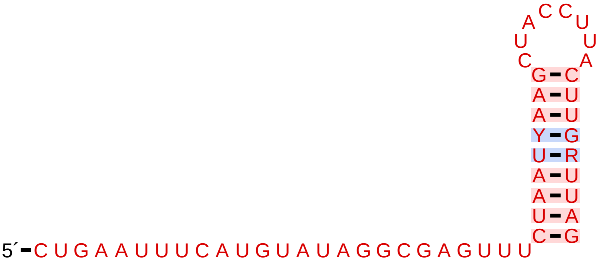

### SampleDMeta_17_1

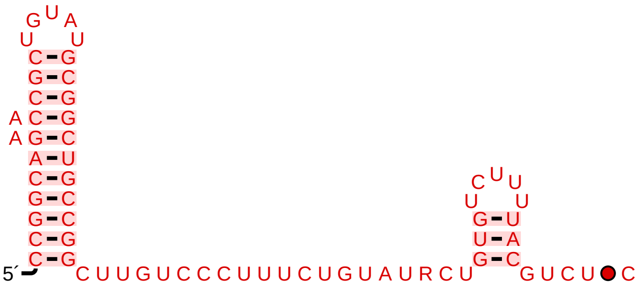

### SampleDMeta_171_1

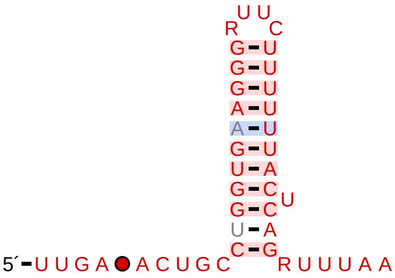

### SampleDMeta_174_1

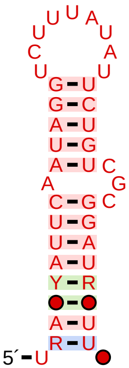

SampleDMeta_177_1

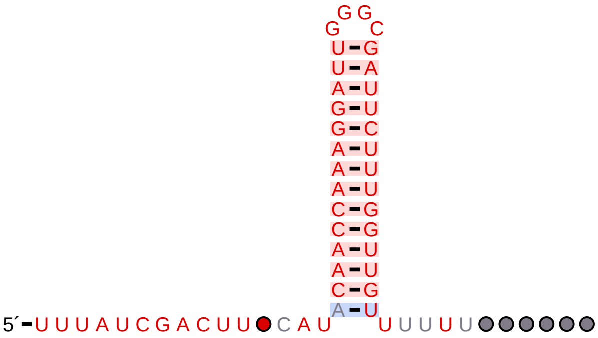

### SampleDMeta_178_1

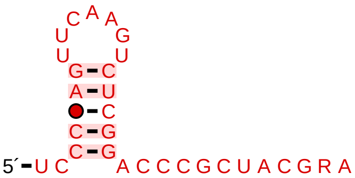

### SampleDMeta_182_1

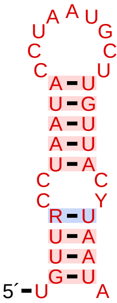

### SampleDMeta_184_1

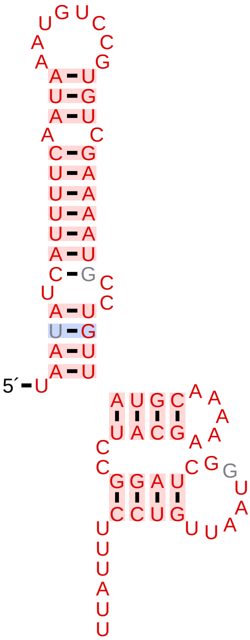

### SampleDMeta_185_1

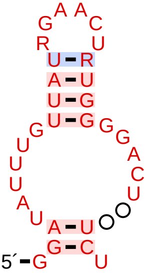

### SampleDMeta_188_1

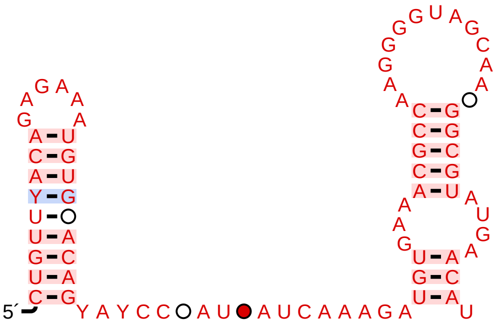

### SampleDMeta_189_1

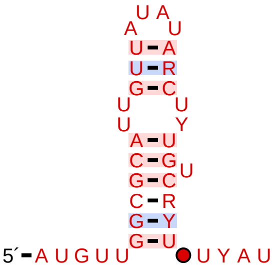

### SampleDMeta_190_1

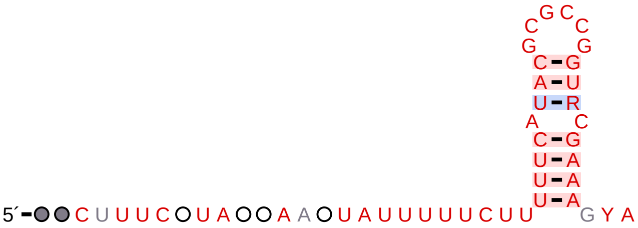

### SampleDMeta_191_1

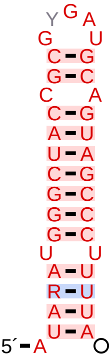

### SampleDMeta_194_1

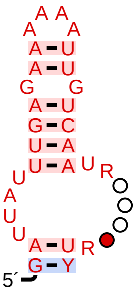

### SampleDMeta_196_1

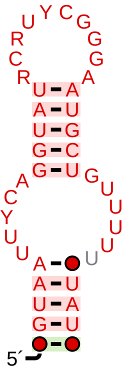

### SampleDMeta_200_1

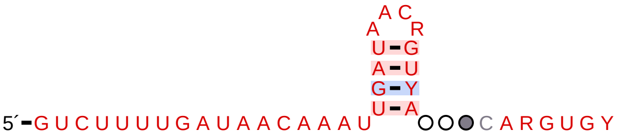

SampleDMeta_204_1

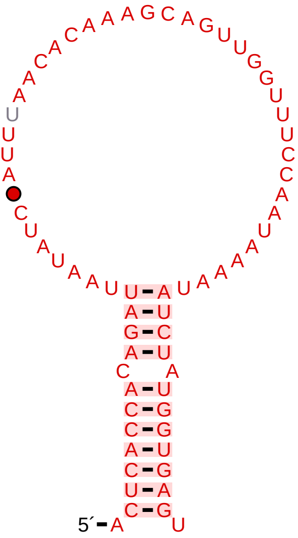

### SampleDMeta_205_1

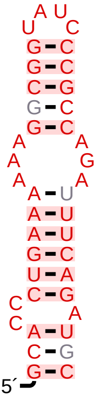

### SampleDMeta_206_1

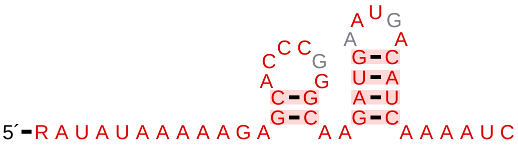

### SampleDMeta_207_1

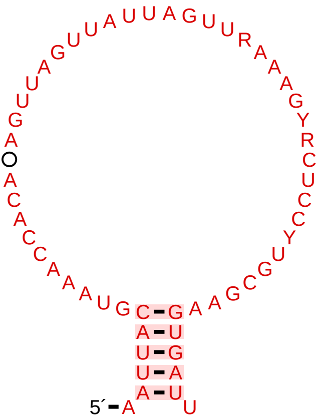

### SampleDMeta_208_1

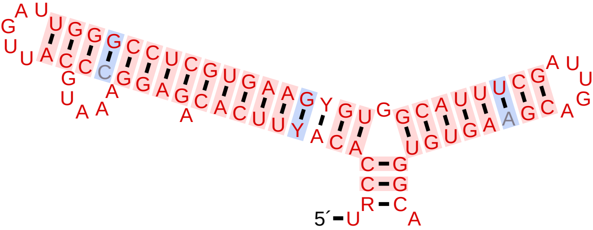

### SampleDMeta_209_1

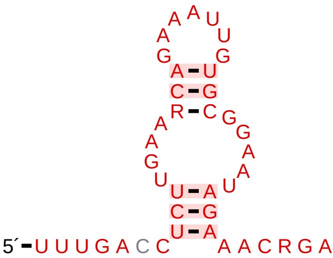

### SampleDMeta_21_1

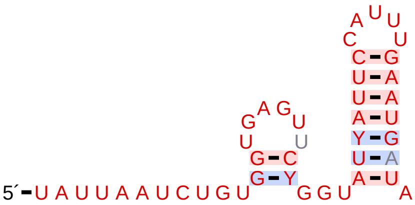

### SampleDMeta_213_1

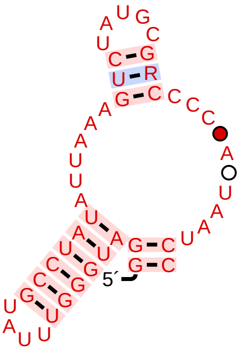

### SampleDMeta_214_1

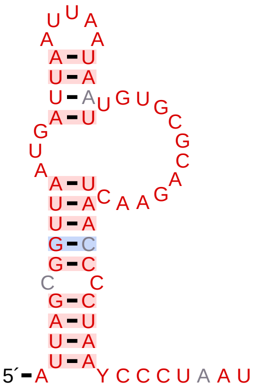

### SampleDMeta_217_1

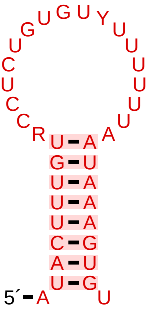

### SampleDMeta_219_1

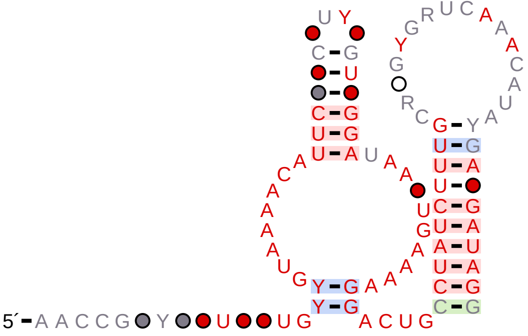

### SampleDMeta_22_1

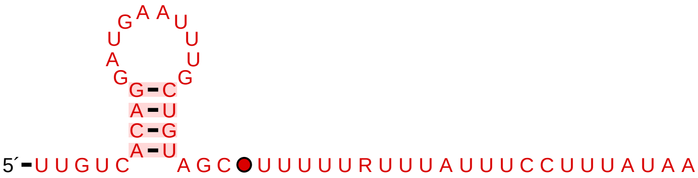

SampleDMeta_220_1

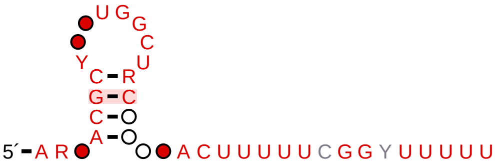

### SampleDMeta_223_1

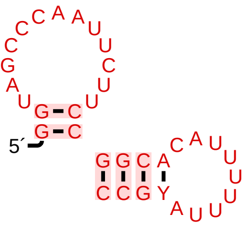

SampleDMeta_224_1

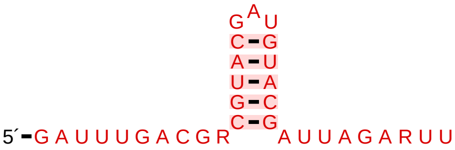

### SampleDMeta_225_1

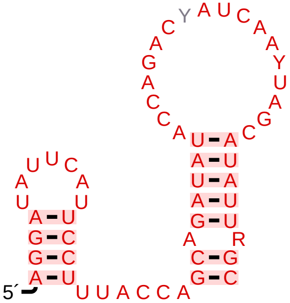

### SampleDMeta_227_1

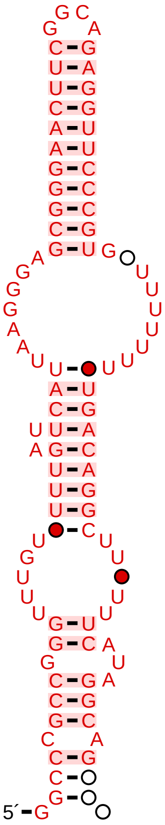

### SampleDMeta_229_1

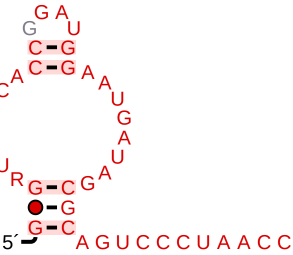

### SampleDMeta_230_1

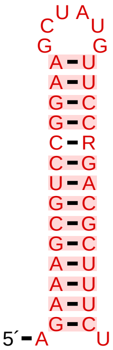

### SampleDMeta_231_1

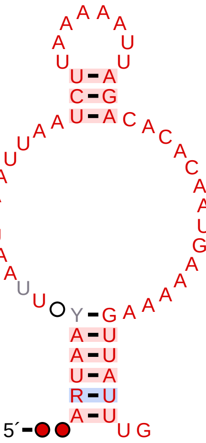

### SampleDMeta_236_1

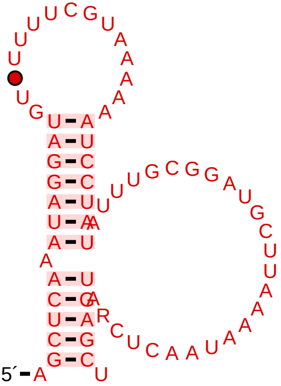

### SampleDMeta_240_1

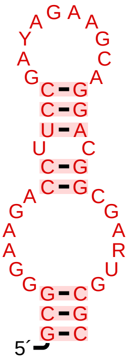

### SampleDMeta_241_1

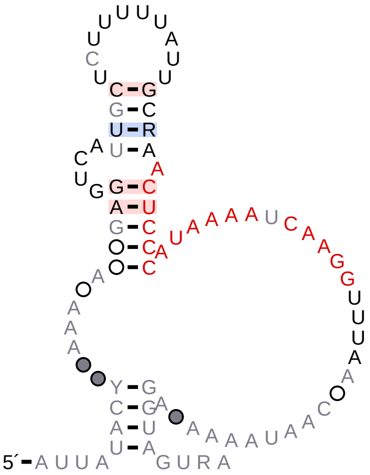

### SampleDMeta_243_1

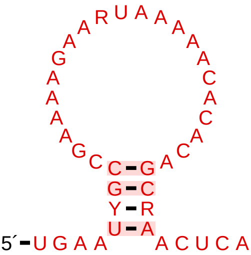

### SampleDMeta_245_1

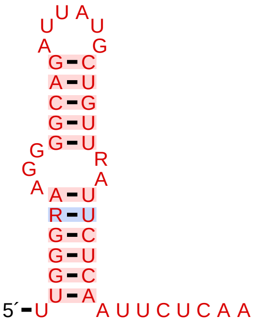

### SampleDMeta_248_1

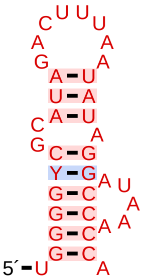

SampleDMeta_249_1

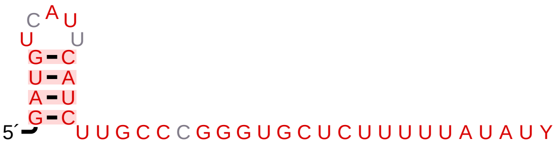

### SampleDMeta_25_1

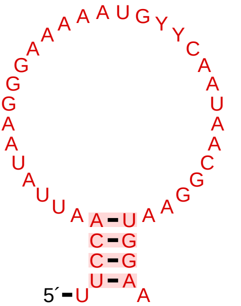

### SampleDMeta_251_1

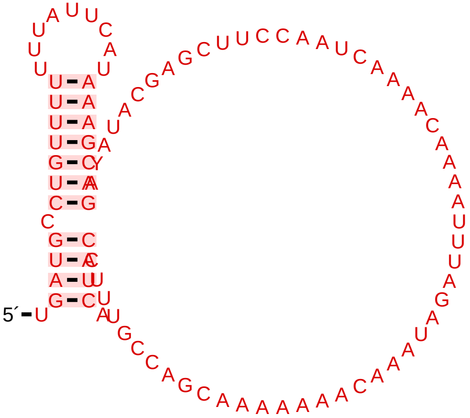

SampleDMeta_253_1

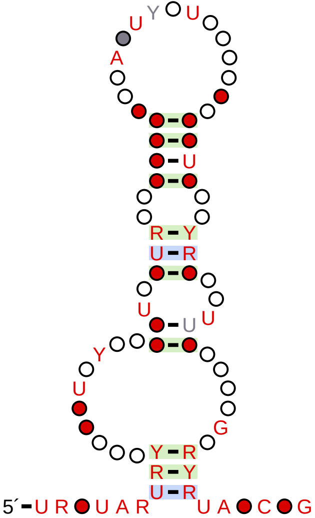

SampleDMeta_257_1

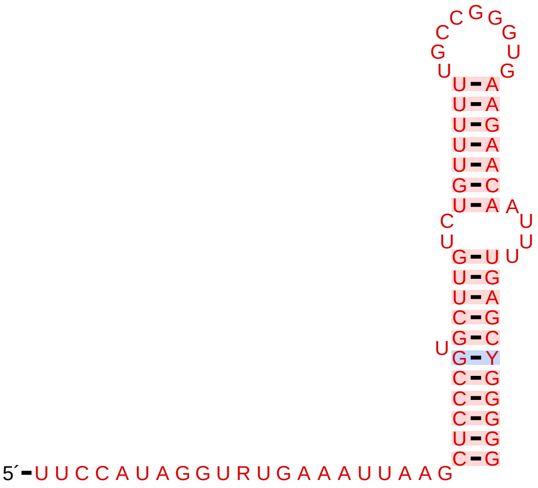

### SampleDMeta_258_1

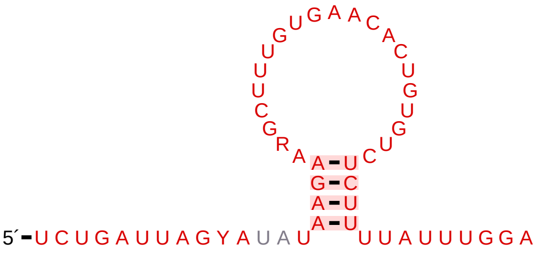

### SampleDMeta_260_1

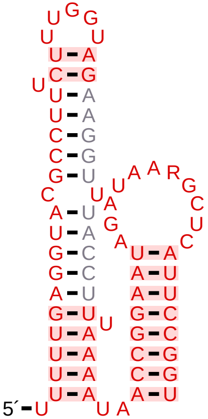

### SampleDMeta_262_1

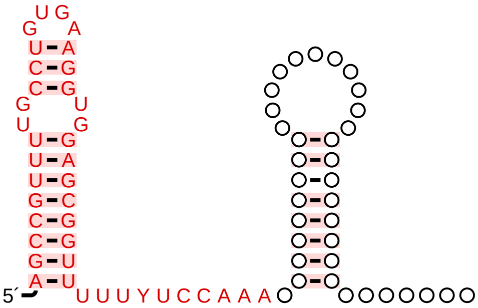

### SampleDMeta_267_1

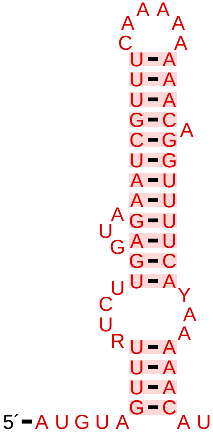

SampleDMeta_27_1

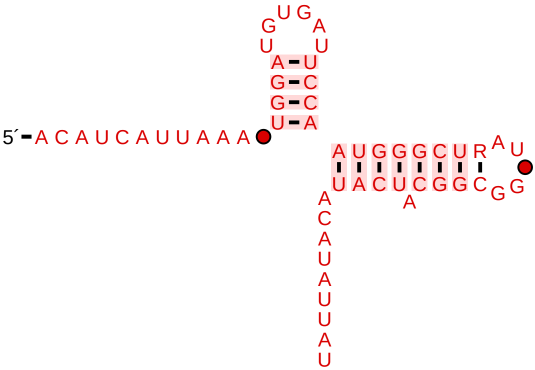

### SampleDMeta_272_1

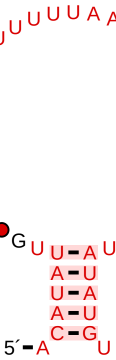

### SampleDMeta_277_1

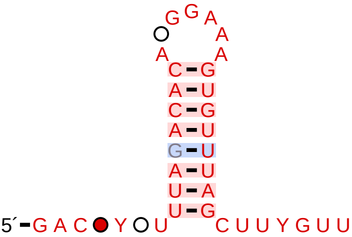

### SampleDMeta_282_1

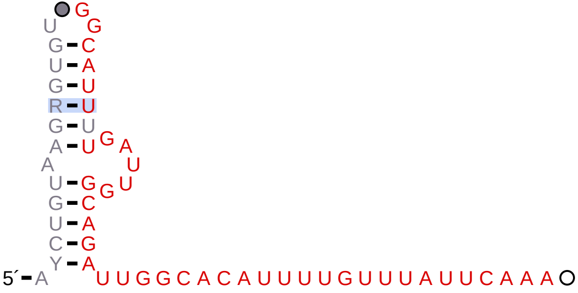

### SampleDMeta_284_1

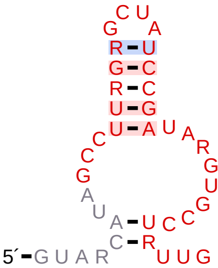

### SampleDMeta_285_1

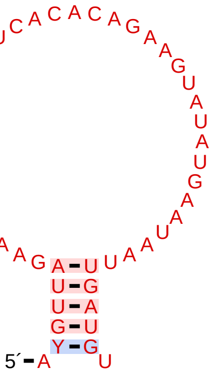

### SampleDMeta_286_1

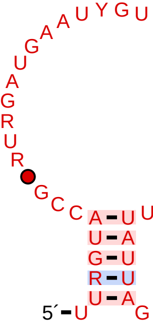

SampleDMeta_288_1

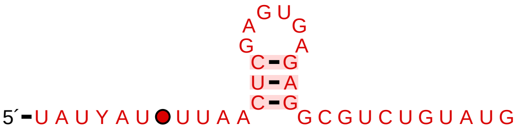

SampleDMeta_29_1

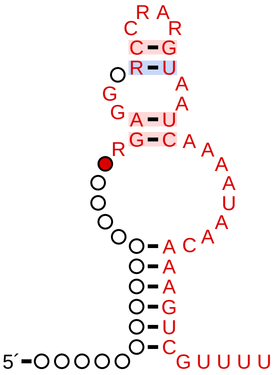

### SampleDMeta_290_1

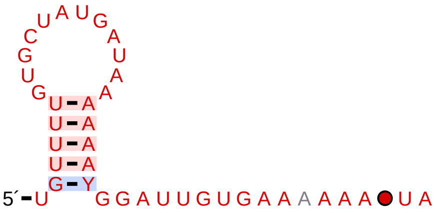

### SampleDMeta_291_1

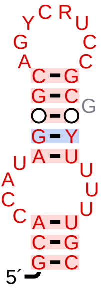

### SampleDMeta_293_1

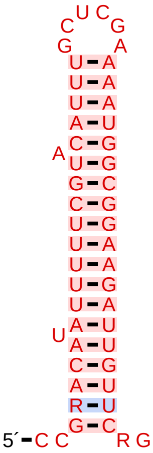

### SampleDMeta_296_1

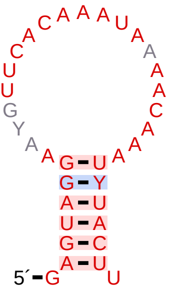

### SampleDMeta_299_1

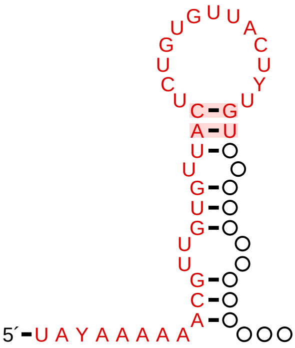

### SampleDMeta_30_1

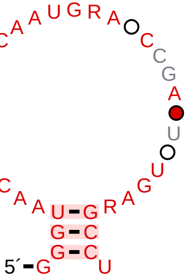

### SampleDMeta_301_1

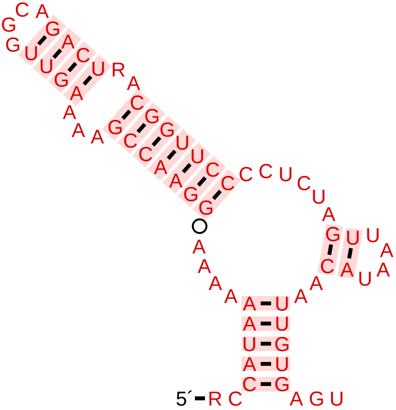

### SampleDMeta_303_1

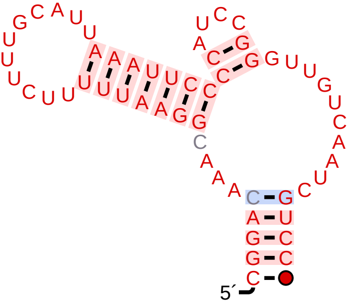

SampleDMeta_304_1

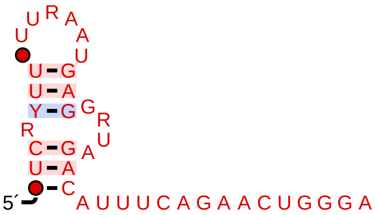

### SampleDMeta_308_1

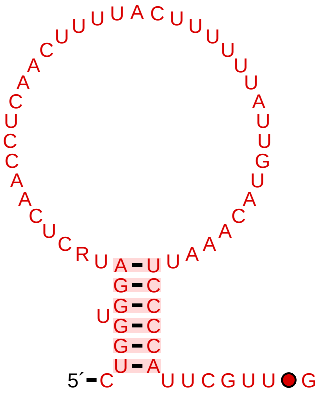

SampleDMeta_31_1

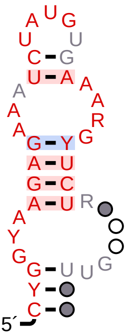

### SampleDMeta_311_1

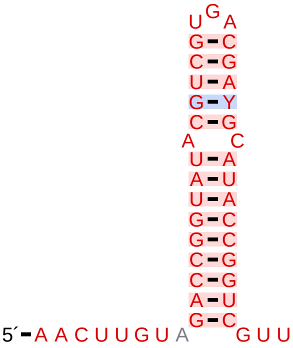

### SampleDMeta_313_1

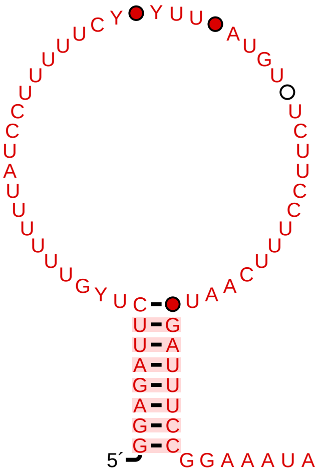

SampleDMeta_315_1

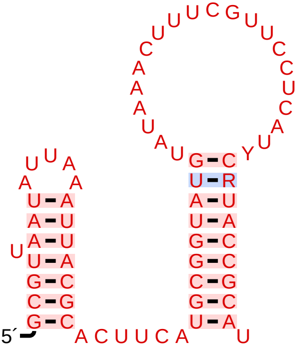

### SampleDMeta_316_1

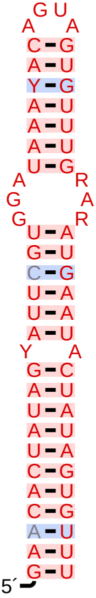

### SampleDMeta_317_1

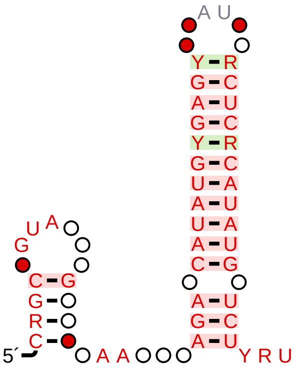

### SampleDMeta_318_1

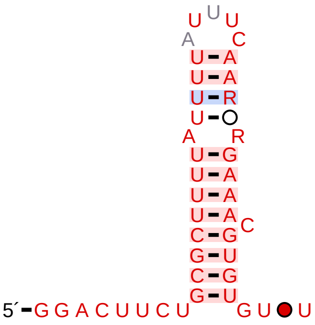

### SampleDMeta_32_1

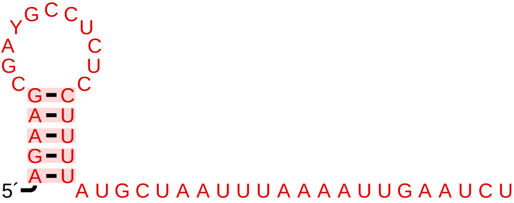

### SampleDMeta_320_1

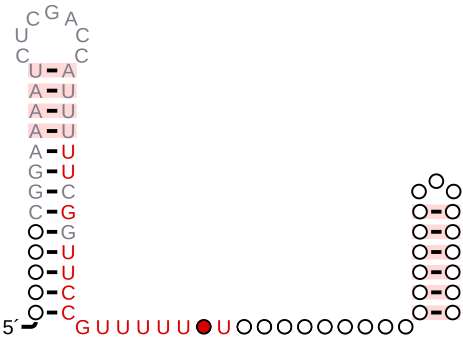

### SampleDMeta_321_1

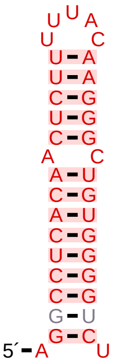

SampleDMeta_323_1

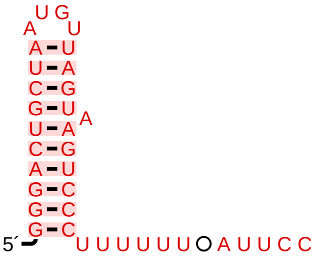

### SampleDMeta_326_1

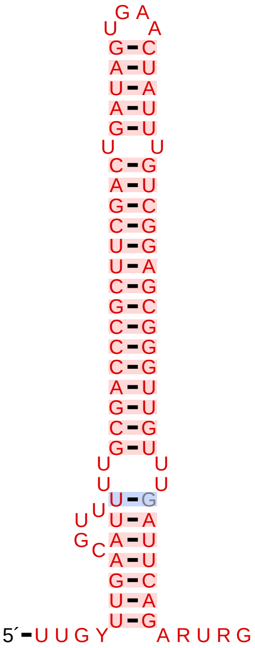

### SampleDMeta_329_1

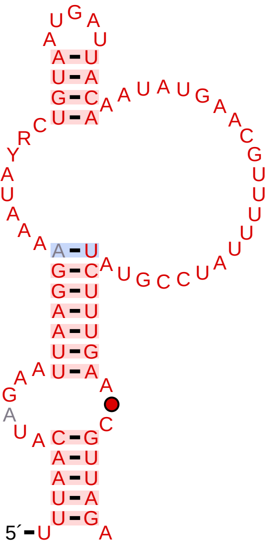

### SampleDMeta_33_1

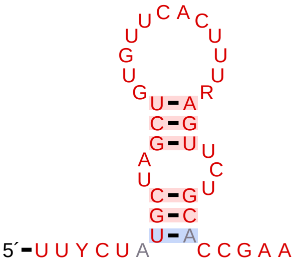

### SampleDMeta_330_1

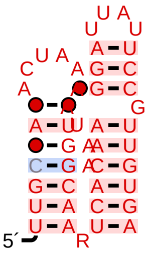

### SampleDMeta_333_1

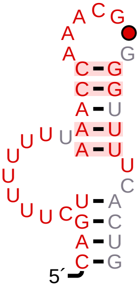

SampleDMeta_336_1

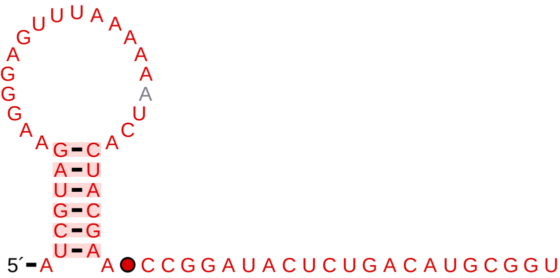

SampleDMeta_337_1

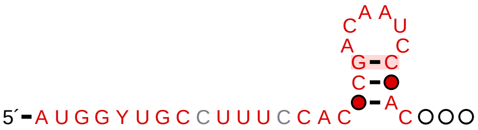

### SampleDMeta_338_1

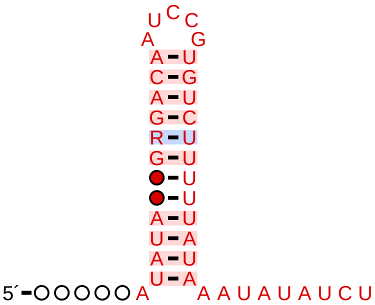

### SampleDMeta_339_1

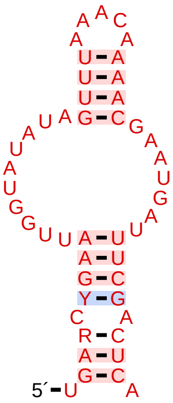

SampleDMeta_341_1

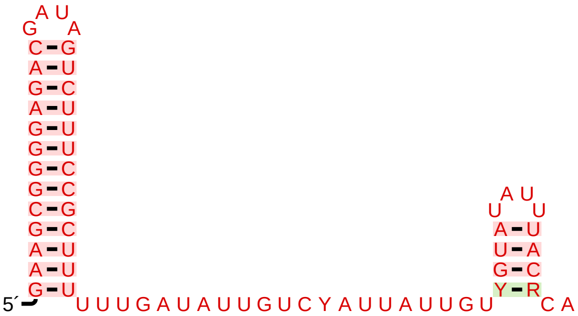

SampleDMeta_344_1

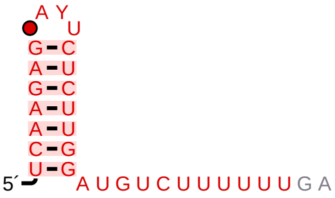

SampleDMeta_345_1

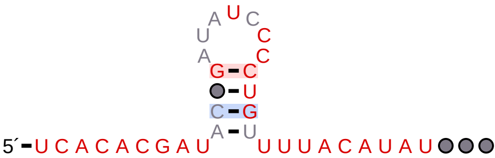

### SampleDMeta_348_1

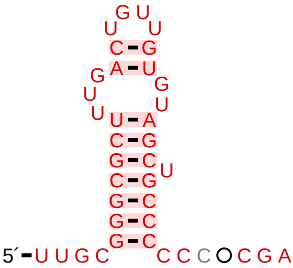

SampleDMeta_349_1

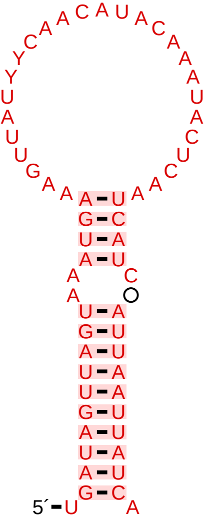

### SampleDMeta_352_1

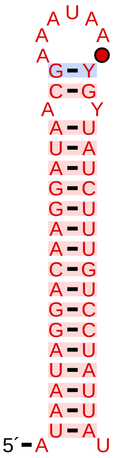

SampleDMeta_353_1

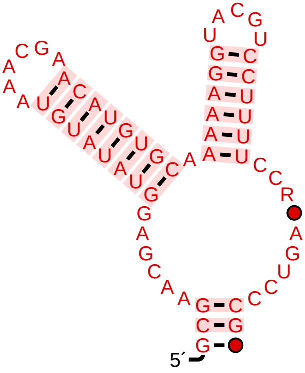

### SampleDMeta_356_1

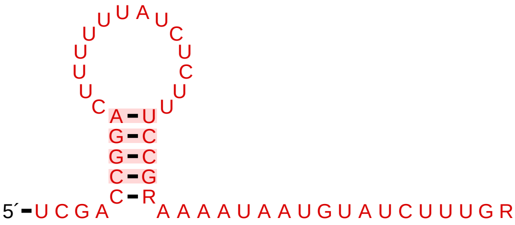

### SampleDMeta_357_1

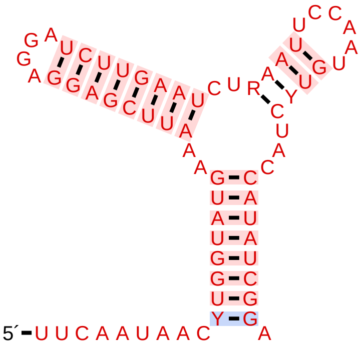

### SampleDMeta_358_1

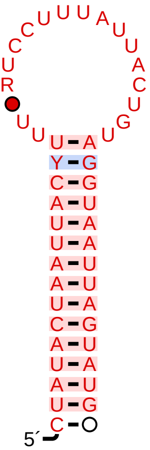

SampleDMeta_359_1

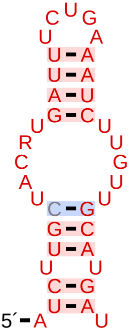

### SampleDMeta_36_1

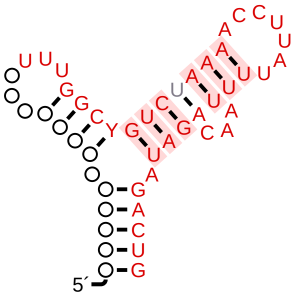

### SampleDMeta_360_1

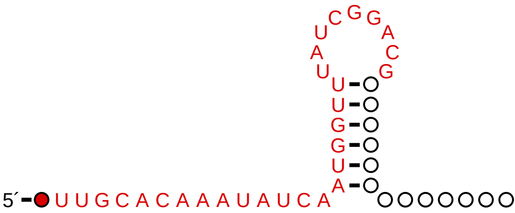

SampleDMeta_361_1

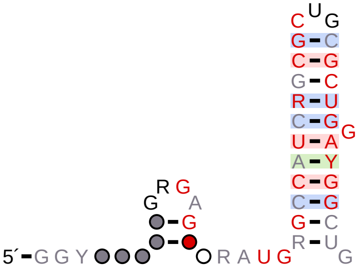

### SampleDMeta_366_1

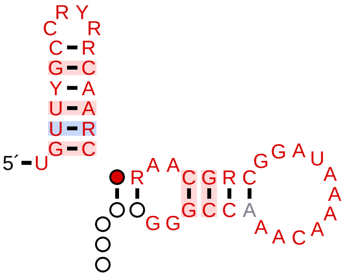

### SampleDMeta_370_1

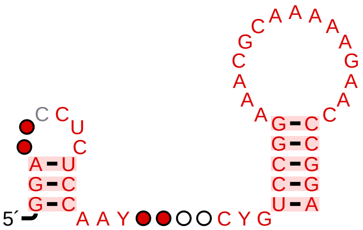

SampleDMeta_372_1

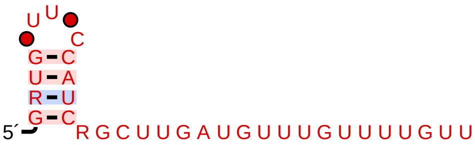

### SampleDMeta_373_1

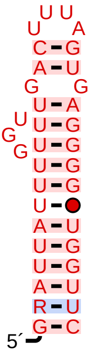

### SampleDMeta_374_1

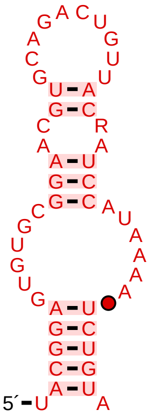

SampleDMeta_375_1

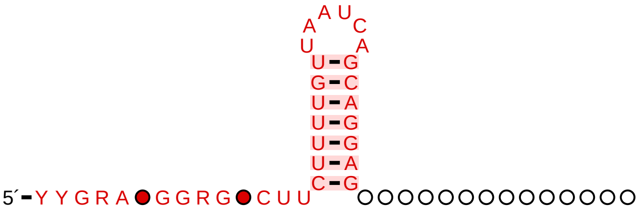

### SampleDMeta_378_1

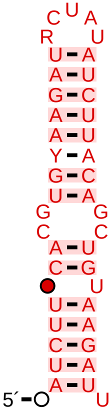

### SampleDMeta_379_1

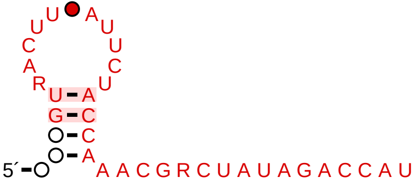

### SampleDMeta_38_1

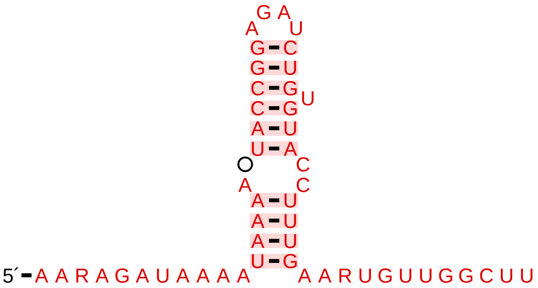

### SampleDMeta_382_1

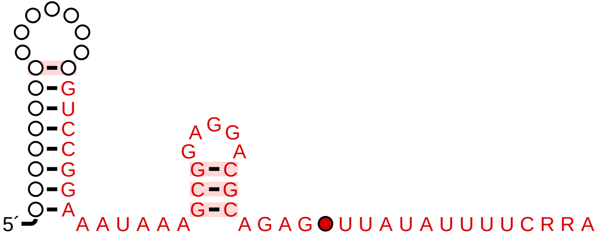

### SampleDMeta_383_1

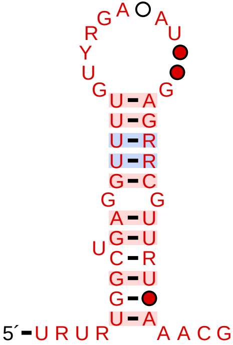

### SampleDMeta_384_1

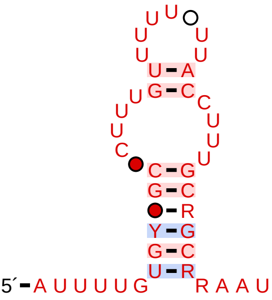

### SampleDMeta_386_1

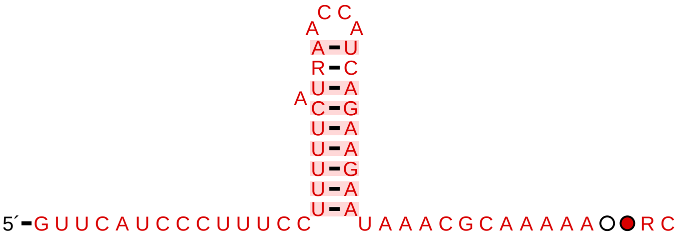

### SampleDMeta_387_1

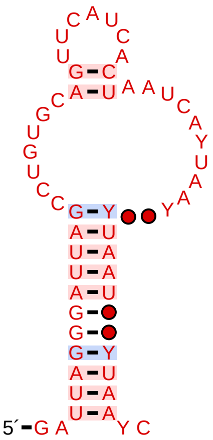

### SampleDMeta_389_1

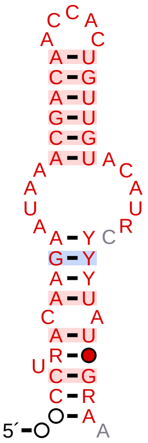

### SampleDMeta_39_1

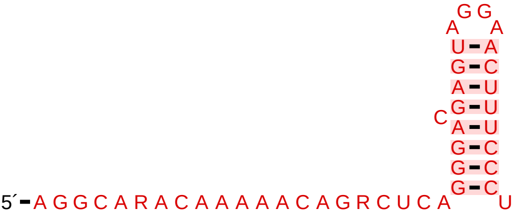

SampleDMeta_393_1

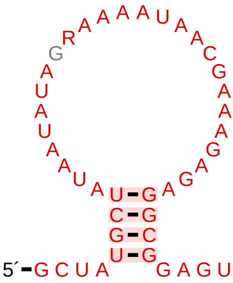

### SampleDMeta_397_1

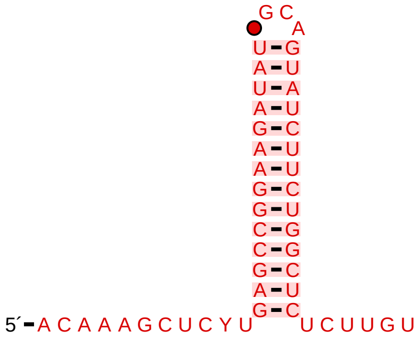

### SampleDMeta_402_1

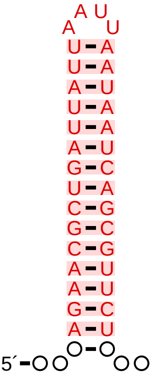

SampleDMeta_405_1

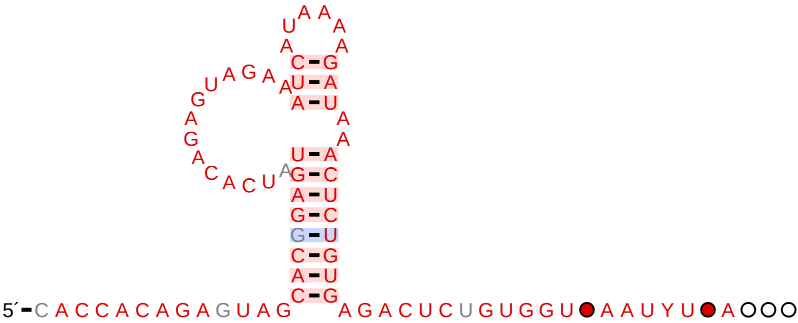

### SampleDMeta_406_1

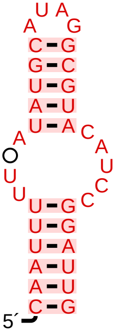

### SampleDMeta_407_1

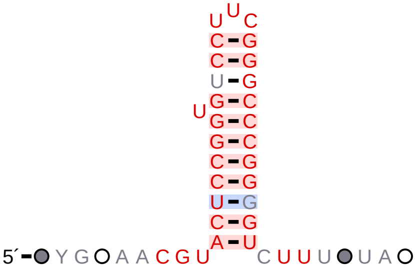

### SampleDMeta_409_1

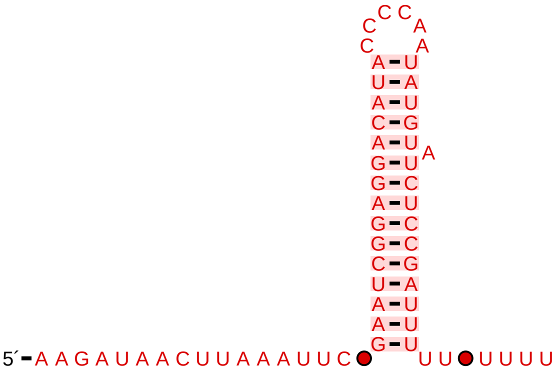

### SampleDMeta_411_1

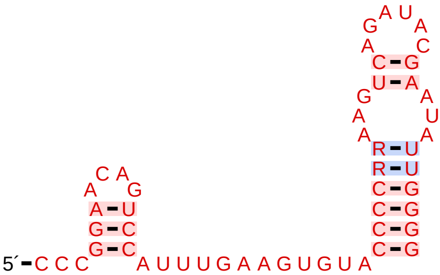

SampleDMeta_414_1

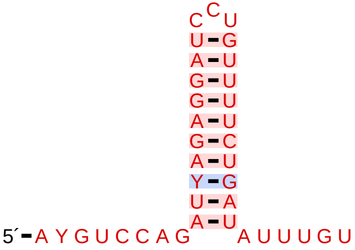

### SampleDMeta_415_1

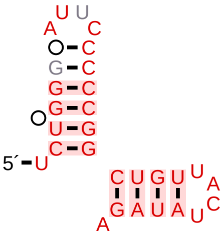

### SampleDMeta_417_1

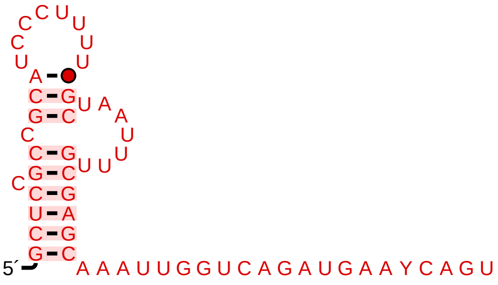

### SampleDMeta_419_1

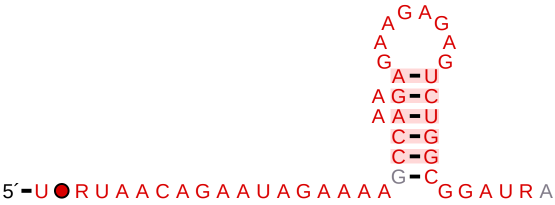

### SampleDMeta_424_1

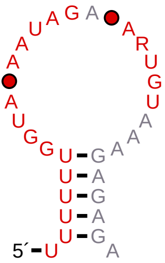

SampleDMeta_427_1

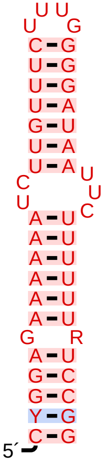

### SampleDMeta_433_1

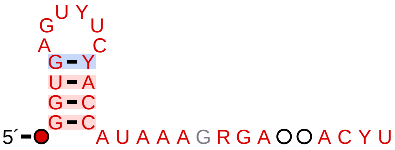

SampleDMeta_437_1

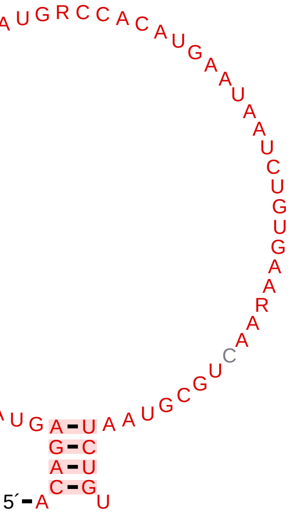

### SampleDMeta_439_1

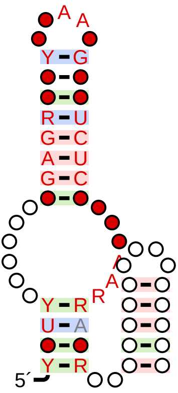

### SampleDMeta_44_1

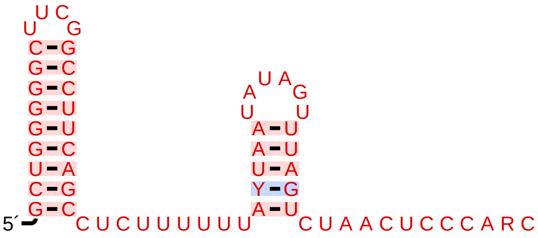

### SampleDMeta_440_1

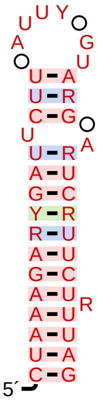

SampleDMeta_441_1

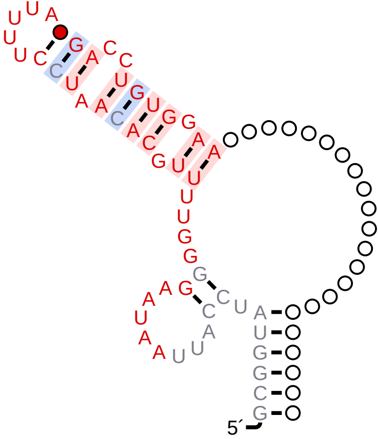

SampleDMeta_443_1

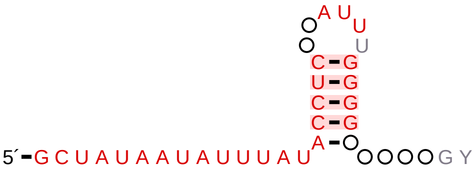

### SampleDMeta_444_1

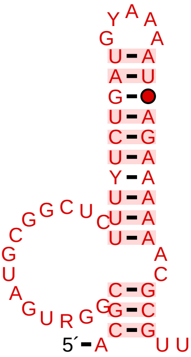

SampleDMeta_445_1

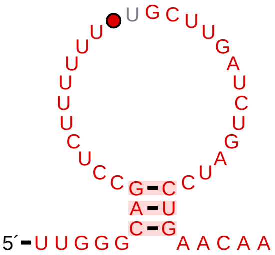

SampleDMeta_448_1

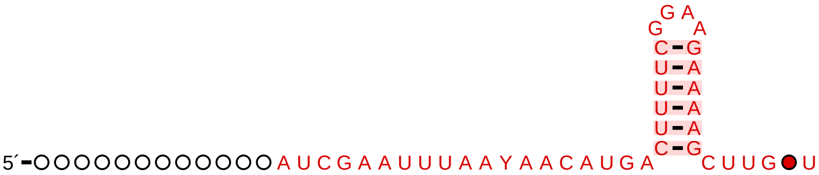

SampleDMeta_45_1

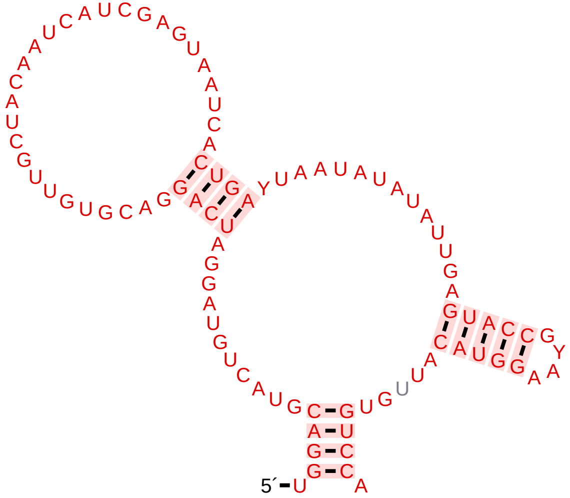

SampleDMeta_451_1

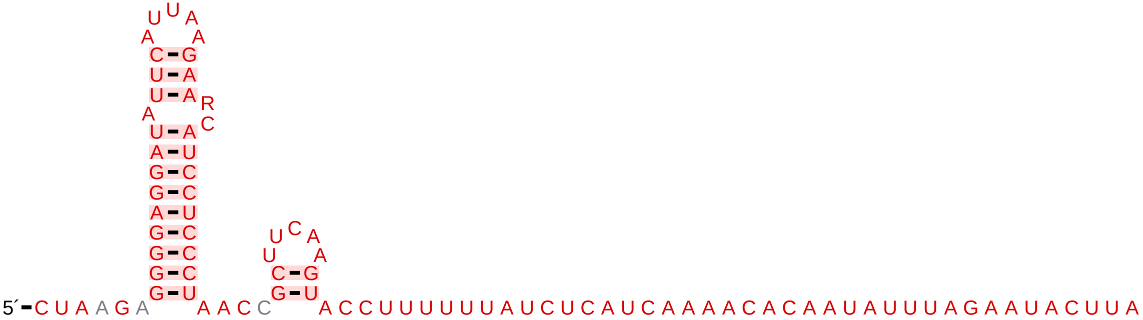

### SampleDMeta_454_1

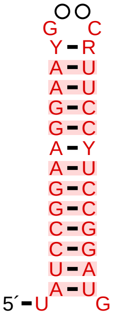

### SampleDMeta_455_1

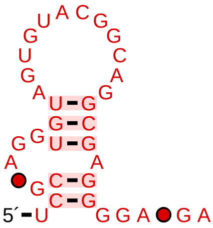

### SampleDMeta_456_1

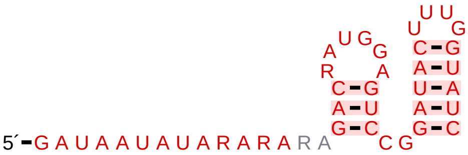

SampleDMeta_458_1

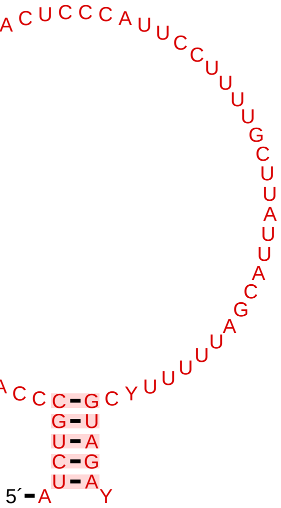

### SampleDMeta_459_1

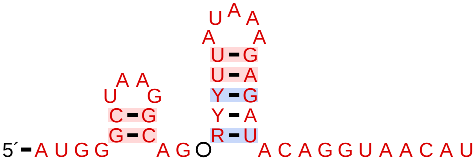

### SampleDMeta_460_1

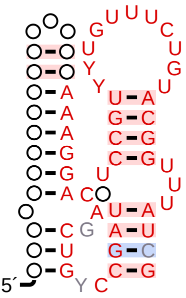

### SampleDMeta_462_1

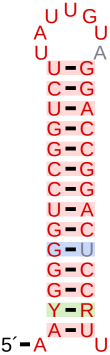

### SampleDMeta_463_1

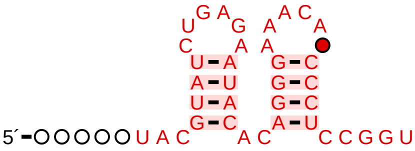

### SampleDMeta_464_1

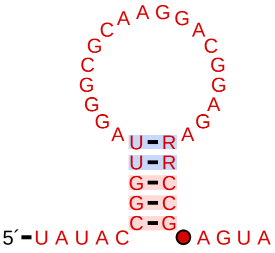

### SampleDMeta_468_1

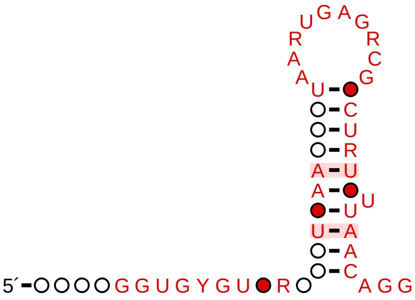

### SampleDMeta_471_1

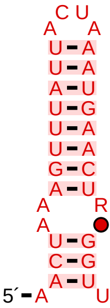

### SampleDMeta_474_1

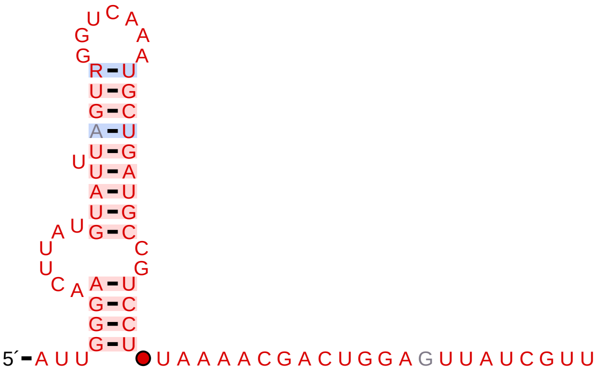

### SampleDMeta_475_1

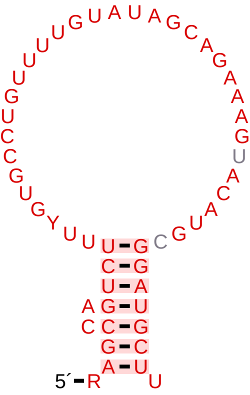

### SampleDMeta_479_1

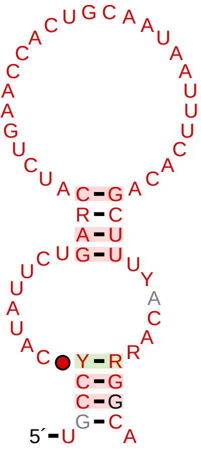

### SampleDMeta_482_1

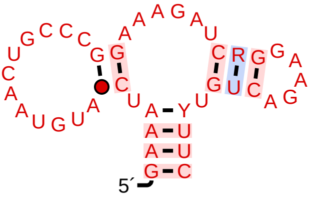

### SampleDMeta_483_1

5´- R U C G U U A U C C C U U U Y U U U

### SampleDMeta_484_1

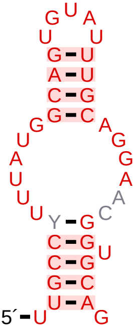

### SampleDMeta_485_1

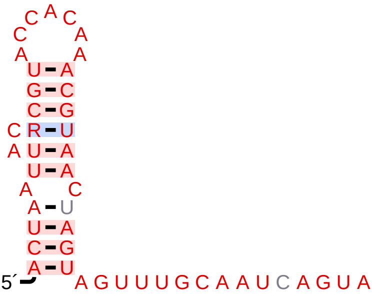

### SampleDMeta_489_1

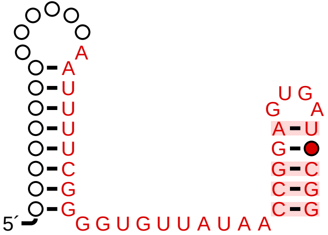

### SampleDMeta_49_1

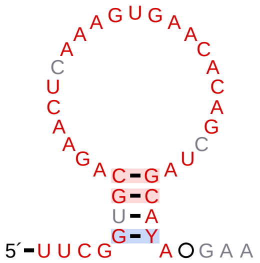

### SampleDMeta_491_1

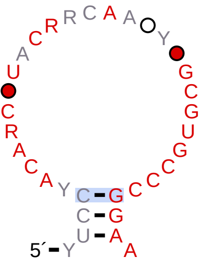

### SampleDMeta_492_1

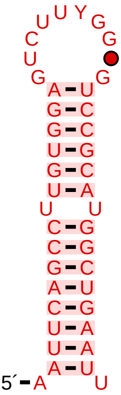

### SampleDMeta_494_1

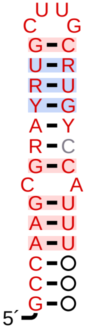

### SampleDMeta_497_1

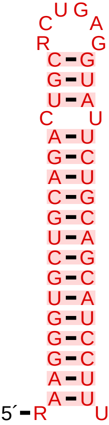

### SampleDMeta_500_1

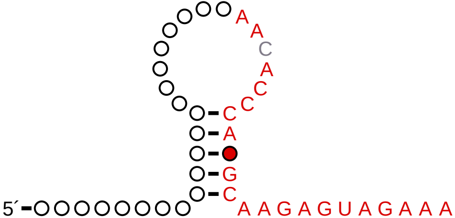

SampleDMeta_501_1

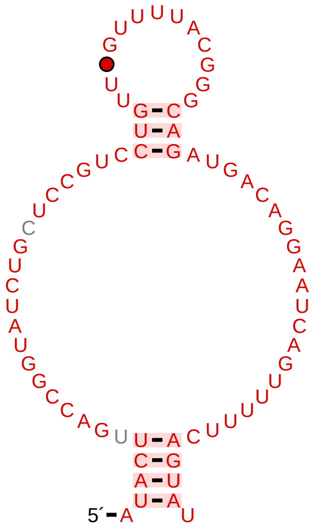

### SampleDMeta_503_1

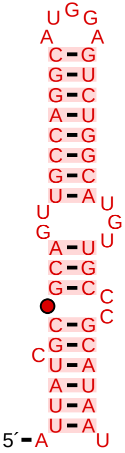

SampleDMeta_506_1

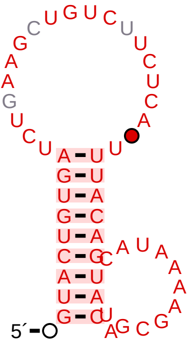

SampleDMeta_507_1

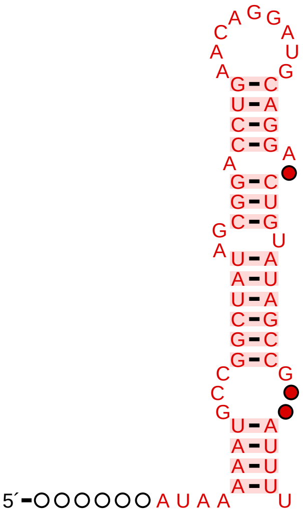

### SampleDMeta_508_1

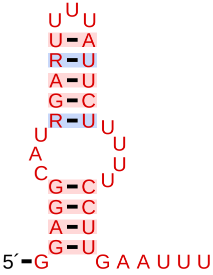

SampleDMeta_509_1

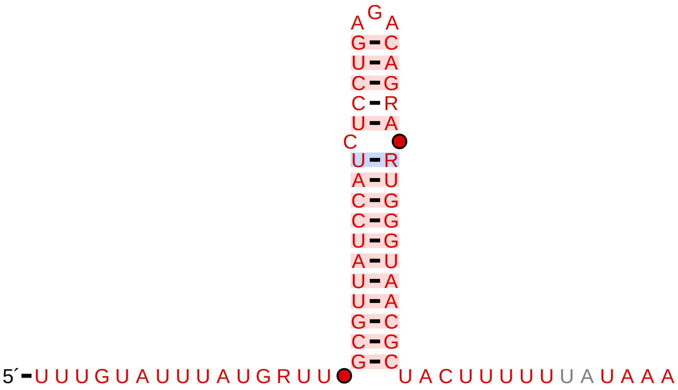

### SampleDMeta_51_1

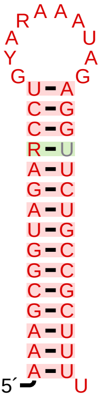

### SampleDMeta_512_1

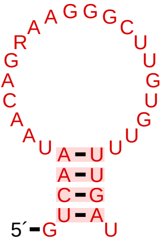

### SampleDMeta_513_1

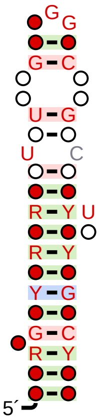

### SampleDMeta_516_1

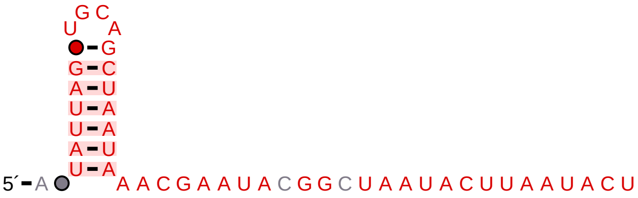

### SampleDMeta_517_1

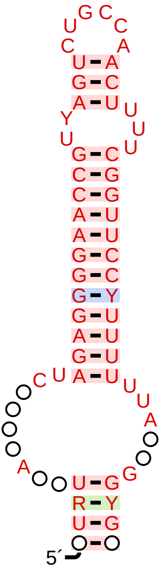

### SampleDMeta_519_1

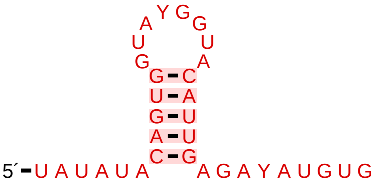

### SampleDMeta_52_1

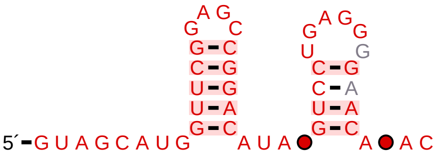

### SampleDMeta_520_1

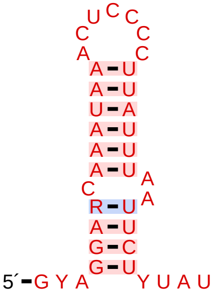

SampleDMeta_525_1

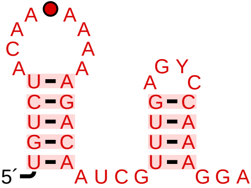

### SampleDMeta_527_1

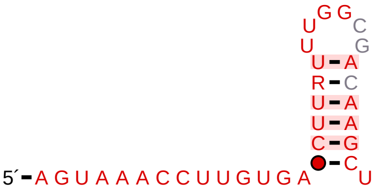

### SampleDMeta_528_1

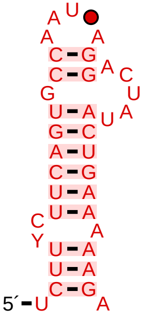

### SampleDMeta_529_1

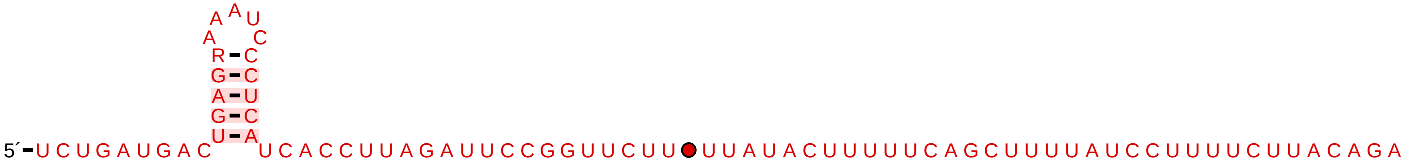

SampleDMeta_54_1

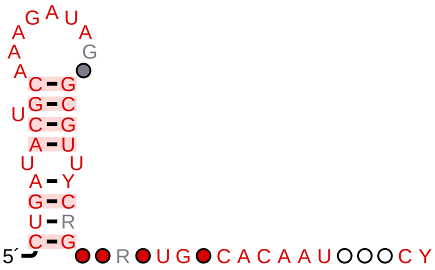

### SampleDMeta_55_1

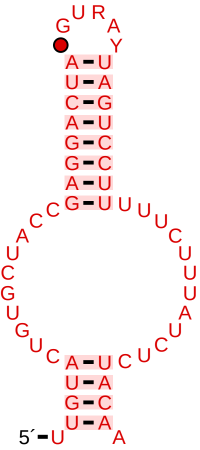

SampleDMeta_59_1

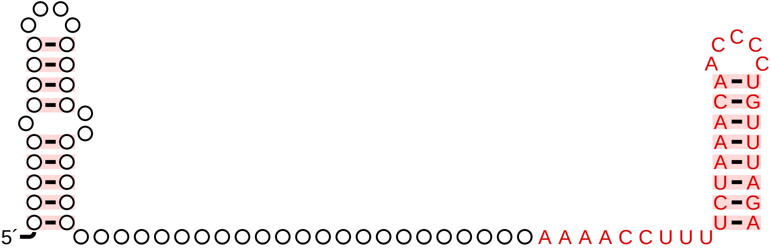

SampleDMeta_6_1

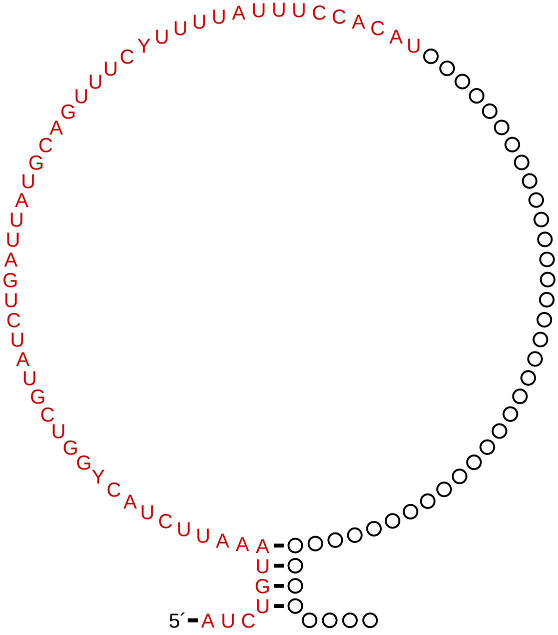

### SampleDMeta_60_1

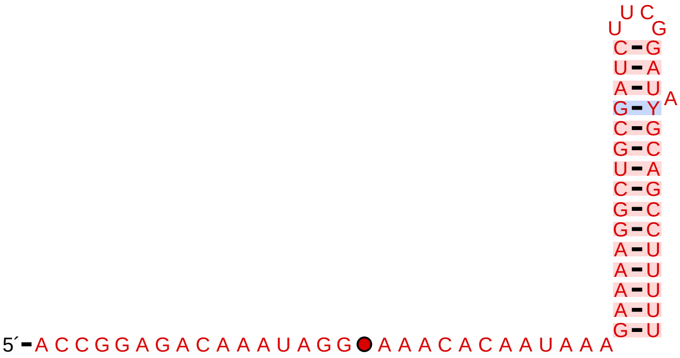

### SampleDMeta_61_1

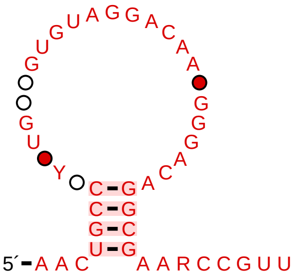

### SampleDMeta_64_1

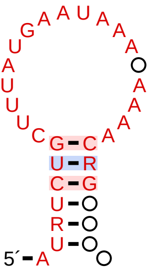

### SampleDMeta_66_1

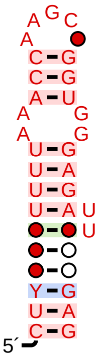

### SampleDMeta_71_1

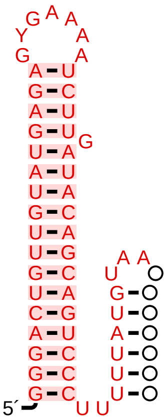

### SampleDMeta_88_1

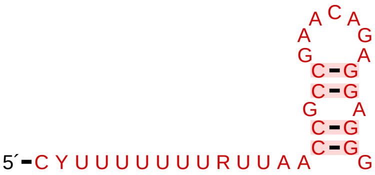

### SampleDMeta_91_1

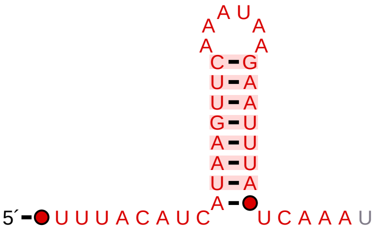
